# Protein structure-based gene expression signatures

**DOI:** 10.1101/2020.06.03.133066

**Authors:** R. Rahman, Y. Xiong, J. G. C. van Hasselt, J. Hansen, E. A. Sobie, M. R. Birtwistle, E. Azeloglu, R. Iyengar, A. Schlessinger

## Abstract

Gene expression signatures (GES) connect phenotypes to mRNA expression patterns, providing a powerful approach to define cellular identity, function, and the effects of perturbations. However, the use of GES has suffered from vague assessment criteria and limited reproducibility. The structure of proteins defines the functional capability of genes, and hence, we hypothesized that enrichment of structural features could be a generalizable representation of gene sets. We derive structural gene expression signatures (sGES) using features from various levels of protein structure (e.g. domain, fold) encoded by the transcribed genes in GES, to describe cellular phenotypes. Comprehensive analyses of data from the Genotype-Tissue Expression Project (GTEx), ARCHS4, and mRNA expression of drug effects on cardiomyocytes show that structural GES (sGES) are useful for identifying robust signatures of biological phenomena. sGES also enables the characterization of signatures across experimental platforms, facilitates the interoperability of expression datasets, and can describe drug action on cells.

## Introduction

Gene expression signatures (GES) are generally defined as a ranked list of genes whose differential expression is associated with a defined biological phenomenon (*1–3*). GES are typically obtained by measuring the transcriptional level of genes through RNA sequencing or previously by array-based experiments. Often, GES sets are determined, for example, by taking the top 100 or 200 highly expressed genes, or by using particular *p*-value cutoffs (*3*). Thousands of GES have been identified and claimed to characterize a wide variety of biological phenomenon (*1–3*). GES have been used to characterize subcellular and whole cell functions (*4, 5*), pathological states (*6, 7*) and cellular response to perturbagens (*8*). However, due to differences in technology, normalization protocols, and practices across laboratories, there is variability in identifying robust GES for a given phenotype, which has hindered its utility in the clinic (*9, 10*).

Determining the reproducibility of a signature for a phenotype of interest remains a challenge, often requiring meta-analyses of existing GES to validate a signature for a phenotype (2, 3, 10). This process includes analyses of thousands of independent samples to generate a robust signature for a single phenotype (*11*). For example, GES variability has led to the cancelation of clinical trials that linked endpoints to specific GES and can produce inconsistent results in the classification of patients for distinct subtypes of a cancer (*10, 12*). Numerous studies have analyzed the robustness of gene expression signatures across studies –further highlighting GES limitations (10, 12–16).

One way to improve the robustness of GES is to integrate multiple types of useful biological information (*1, 17*). Because genes encode proteins whose 3D structures execute functions, we hypothesize that enriched protein structures may define a generalizable representation of any given gene set. Particularly, one common structural characteristic of proteins is the overall structural family or “fold” of the protein and/or its individual domains, which have direct association with gene function (*18–20*). For example, incorporating structural information has enhanced the prediction of protein-protein interaction networks and disease pathways (*21, 22*). In this study, we derived higher order structural features from ranked gene lists to yield robust structural GES (sGES). We show that sGES can produce reliable signatures of distinct tissue types. Additionally, integration of sGES with GES, through an autoencoder, can be used to precisely identify outlier samples between distinct gene expression datasets, facilitating interoperability between experiments that use differing transcriptomic methodologies. Finally, we demonstrate that sGES can be used to characterize biological phenomena, such as cellular response to perturbagens, adding an additional dimension of insights to existing transcriptomics analysis.

## Results

### Quantitative metrics to evaluate GES and sGES reproducibility

To characterize reproducibility of signatures of given phenotype we defined relevant, quantitative properties to the describe the quality of a dataset. We posit that a reproducible GES would have three major properties: 1. consistency across independent samples and replications; 2. high predictive capacity for the phenotype using standard performance measures (*23*); and 3. robustness across different measurement platforms.

We use the Jaccard coefficient (J_C_), which measures the overlap between two distinct gene sets, to measure consistency between signatures characterizing the same phenotype in independent samples (Methods; **Fig. 1A**). A high J_C_ demonstrates that the gene signature is consistent across experimental samples. Furthermore, a low variance for a distribution JC values across several hundred independent samples may indicate signatures with high reproducibility.

**Fig. 1:**
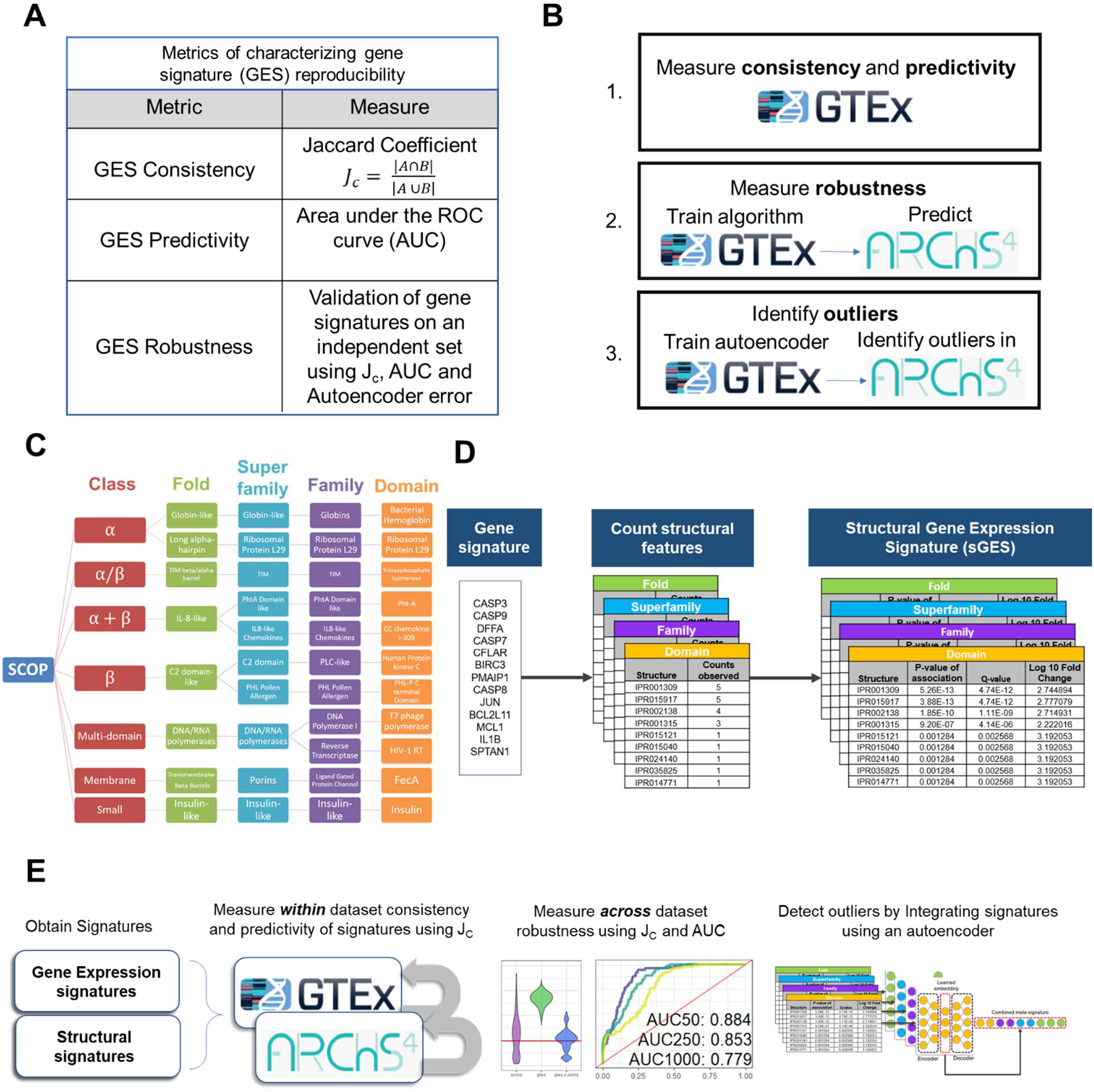
Study design. A) Evaluation metrics for GES (GS) consistency, predictivity, and robustness. B) Approach of measuring consistency, robustness and outlier detection. C) SCOPe hierarchy of protein structural features, with examples. D) Workflow to generate structural gene expression signatures (sGES). E) Workflow for evaluating the reproducibility of GES, structural signatures, and integrated signatures from GTEx and ARCHS4.

To evaluate the predictive performance of a signature to a phenotype, we used a standardized machine learning algorithm (a random forest) across all signatures to assess the baseline effectiveness of a given signature, in terms of area under the ROC curve (AUC), without any significant parameter optimizations or feature selection (Methods; **Fig. 1A**). To measure signature robustness, we computed both JC and AUC values of signatures across two independent datasets measuring an identical phenotype (Methods; **Fig. 1A**). We analyzed expression data from GTEx (v8.0), which categorizes 11,688 samples across 53 healthy tissues from 714 donors (**Table S1**)(*24*). We leverage ARCHS4 (*25*), a collection of GES mined from the Gene Expression Omnibus, as an independent, and nonoverlapping, collection of GES of tissue types analyzed in GTEx (**Fig. 1B,D**). The overall workflow is shown in **Fig. 1E**.

### Protein structure enrichment at any level captures relevant biological information from gene expression experiments

Protein structures encoded by the genes constituting the expressed genes may characterize the cell’s phenotype. Therefore, we hypothesize that using features derived from protein structures in GES can improve reproducibility across experimental platforms. A ‘structural gene expression signature’ (sGES) for each gene set was determined by identifying available structural features from the encoded protein of each gene (Methods; **Fig. 1C-D**). We define a structural feature as the structural hierarchy from the Structural Classification of Proteins extended [SCOPe] (*26*) and InterProscan (*27*) databases, where protein domains (10,637 domains, ex. Serine-threonine/tyrosine-protein kinase, catalytic domain) are categorized into families (4,919 families, ex. Protein kinases, catalytic subunit), which are categorized into superfamilies (2,026 superfamilies; Protein kinase-like) and further grouped into distinct folds (1,232; Protein kinase-like) (Methods; **Fig, 1C**). For a given gene set, each structural feature was evaluated for enrichment in the gene set, compared to the counts of the structural feature in the human proteome (Methods; **Fig, 1D**). sGES are defined as the complete set of structural features derived from a ranked list of genes, at each structural level (domain to fold levels).

To determine if protein structure enrichment captures biological information observed in GES, we utilized t-distributed Stochastic Network Embedding (t-SNE) (*28*) to cluster GTEx tissues samples based on top 250 highest expressed genes and their enriched structural features (**Fig. 2, Fig. S1**). We observed that sGES are capable of clustering tissue types at both the lowest structural level (domains) and, surprisingly, the highest structural level (folds). Importantly, the clusters at all structural levels capture functional and spatial relationships among tissues (**Fig. S1**).

**Fig. 2:**
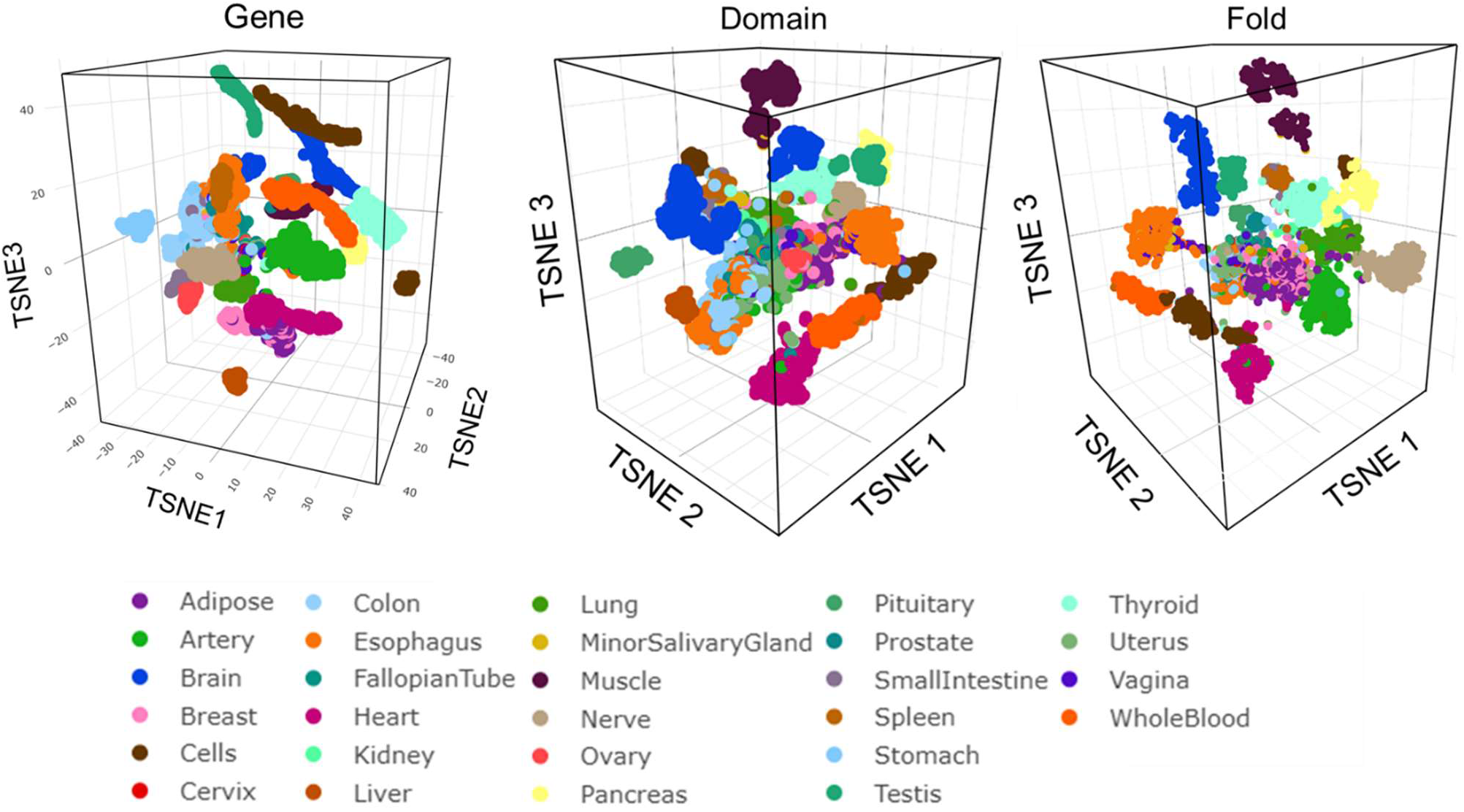
Protein structure enrichment clusters tissue-specific gene expression. The top 250 highest expressed genes from GTEx (in terms of transcripts per million) were obtained. Tissue samples were then clustered based on the presence or absence of the GES using t-SNE. sGES were then derived from the GES, and tissue samples were clustered by using t-SNE based on the presence or absence of structural features at the domain and fold levels. Each sample is colored by tissue type.

For example, ovary tissue sGES cluster near uterine signatures. Both tissues’ GES are enriched with protein domains related to sex hormone production such as Follistatin/Osteonectin EGF domain, Kazal domain, SPARC/Testican, and Fibrillar collagen domain (**Fig. S2**). Both tissues’ sGES also retain tissue-specific domain differences, such as ovarian tissue having structural signatures containing Glutathione transferase domains, which is a biomarker of oocyte viability and quality (*18*). While the uterine tissue enriching for proteins containing structural domains such as the Tubulin/FtsZ domain (*19*).

This result is surprising because conservation of high level protein structure (i.e., fold) is not necessarily always predictive of protein function (*20*), yet, using a representation of folds, domains and families, independently, can constitute an expression signature that captures tissue types.

### sGES improves within-dataset consistency of gene expression data

We computed the JC values for each pair of samples from the same tissue type, using both GES and sGES (**Fig. 3A**). Consistency of sGES, as measured by JC, at all structural levels, increases as higher order sGES are used (**Fig. 3A, Fig. S3**). For a gene set size of 250, sGES significantly increase the mean within-tissue JC compared to GES across all tissue types (**Fig. 3A, Table S2**). For example, at the fold level, the mean within tissue JC reaches a value of 0.75, while the mean across-tissue JC reaches a value of 0.54 (**Fig. 3B**). The increase in JC at higher structural levels indicates increasing consistency of signatures (**Fig. 3A**).

**Fig. 3:**
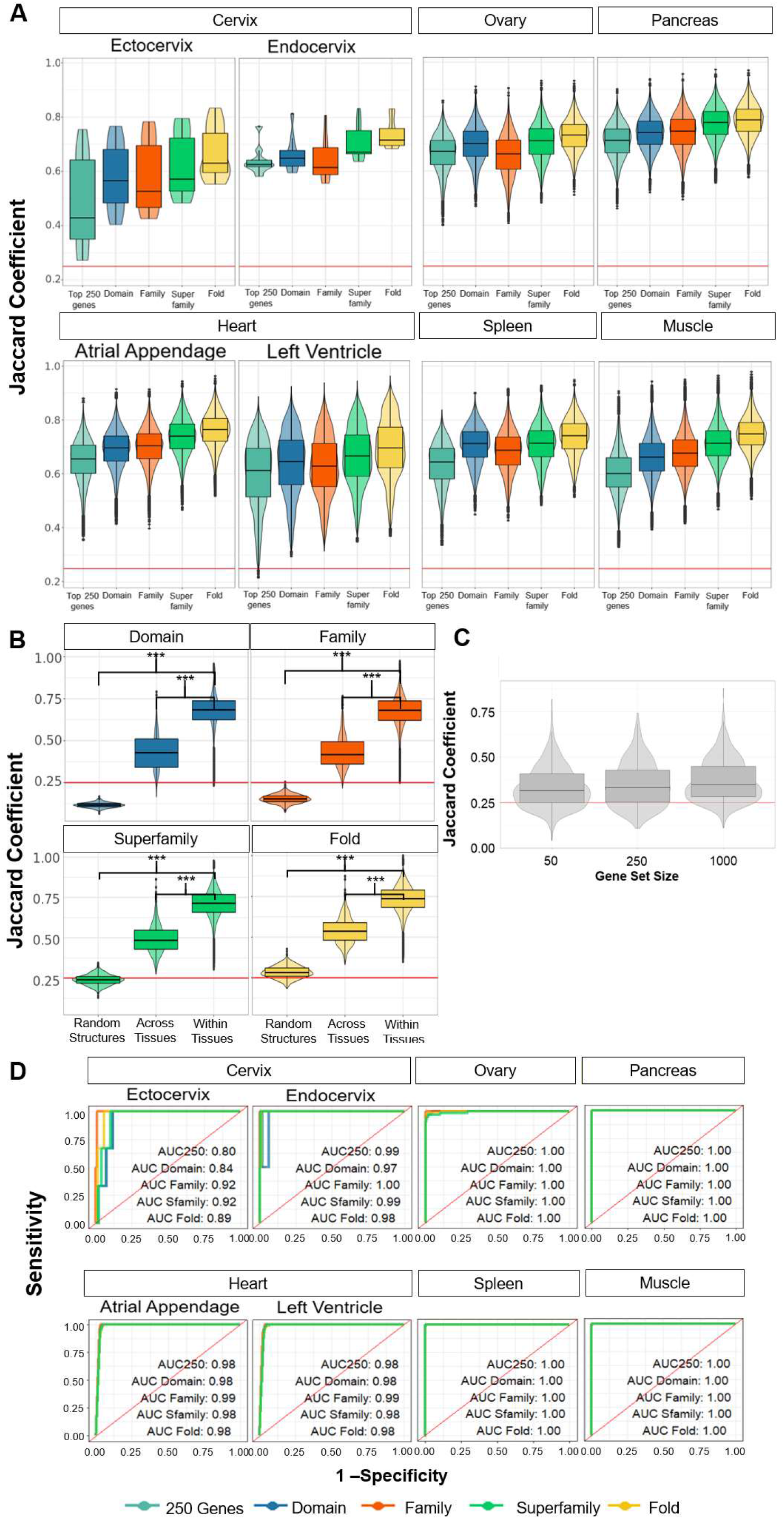
Signature consistency improves using protein structure. A) Distributions of Jaccard coefficient (JC) values within tissue types. For each pairs of samples, in each tissue type (as cataloged by GTEx), a JC was computed for the top 250 highly expressed genes (by TPM) and their derivative sGES at each structural level. The JC is defined as the intersection over the union of two sets and can be thought of as the percentage overlap of two sets. All distributions are statistically significant from each other using pairwise t-tests, with FDR correction (**Table S2**). The red line indicates a JC of 0.25. B) distributions if structures are randomly assigned to each gene (1,000 bootstraps). ‘Across tissues’ are JC distributions between unlike tissue types (1,000 bootstraps). ‘Within tissues’ are the JC distributions between the same tissue type. Within tissue comparisons are significantly higher than random structure comparisons and JC values between distinct tissue type. Red line indicates a JC = 0.25. C) Pairwise GES JC distributions across randomly selected, distinct tissues types, repeated 1,000 times. D) A random forest was trained using GES (of size 250) and sGES at different structural levels (Domain, Family, Superfamily [Sfamily], and Fold) for GTEx tissue expression data. Area under the curves (AUC) are displayed for each structural level.

One explanation for this improvement in consistency may be due to the number of possible structures diminishing as higher order structural features are used (**Fig. S4**). However, we observe that while the mean JC generally increases using sGES, disparate tissue types have significantly lower JC values at each structural level (**Fig. 3B**); retaining tissue-specific information, as observed from the t-SNE (**Fig. 2**). Importantly, the average consistency of dissimilar (or across-tissue) samples using sGES is similar to that of GES (for Domain and Family levels), even at increasing GES sizes (**Fig. 3B-C**). This result asserts that the higher average JC seen in sGES is unlikely to be an artifact of decreasing feature space sizes since GES have similar across-tissue JC values to sGES, despite a higher feature space.

### sGES accurately classifies cell type with a simple machine learning model

A major test of GES reproducibility is the ability of a GES for a phenotype, obtained in one sample, to accurately predict the phenotype for a gene signature derived in an independent sample measuring the same phenotype (i.e., predictivity, **Fig. 1A**). To evaluate the baseline predictivity of both GES and sGES across different tissues, and signature types (Methods), we trained a random forest to identify tissues from either GES of size of 250 or sGES from GTEx expression data. Notably, the parameters for the random forest were standardized and neither feature selection nor parameter optimization was performed on any model (Methods, **Fig. 3D, Fig. S5**).

We observe that both GES and sGES (at any structural level) have high predictivity for any given tissue type within the GTEx dataset, after 10-fold cross validation. For example, the best area under the ROC curve (AUC) values for each tissue range from 0.891 (ectocervix) to 1 (lung) (**Fig. S5**). Importantly, the tissue with the largest variance in JC distribution (i.e., ectocervix) has the lowest predictive performance, indicating a relationship between the two metrics. There are small differences in the predictivity among gene set sizes of 50, 250, and 1,000 across all tissue types within GTEx (**Fig. S6**).

### sGES enable the classification of robust expression signatures across databases

We used an independent validation set from the ARCHS4 (*25*) database to evaluate the robustness of tissue GES and sGES from GTEx data (**Fig. 4, Fig. S7-S10**). In brief, ARCHS4 is a collection of gene expression data derived from the Gene Expression Omnibus (GEO) (*29*), which collates gene expression data generated from a wide variety of sequencing technologies and platforms. Specifically, we evaluated GTEx signatures for consistency (**Fig. 4A-B, Fig. S7-S8**) and predictivity against ARCHS4 (**Fig. 4C-D, Fig. S9-S10**).

**Fig. 4:**
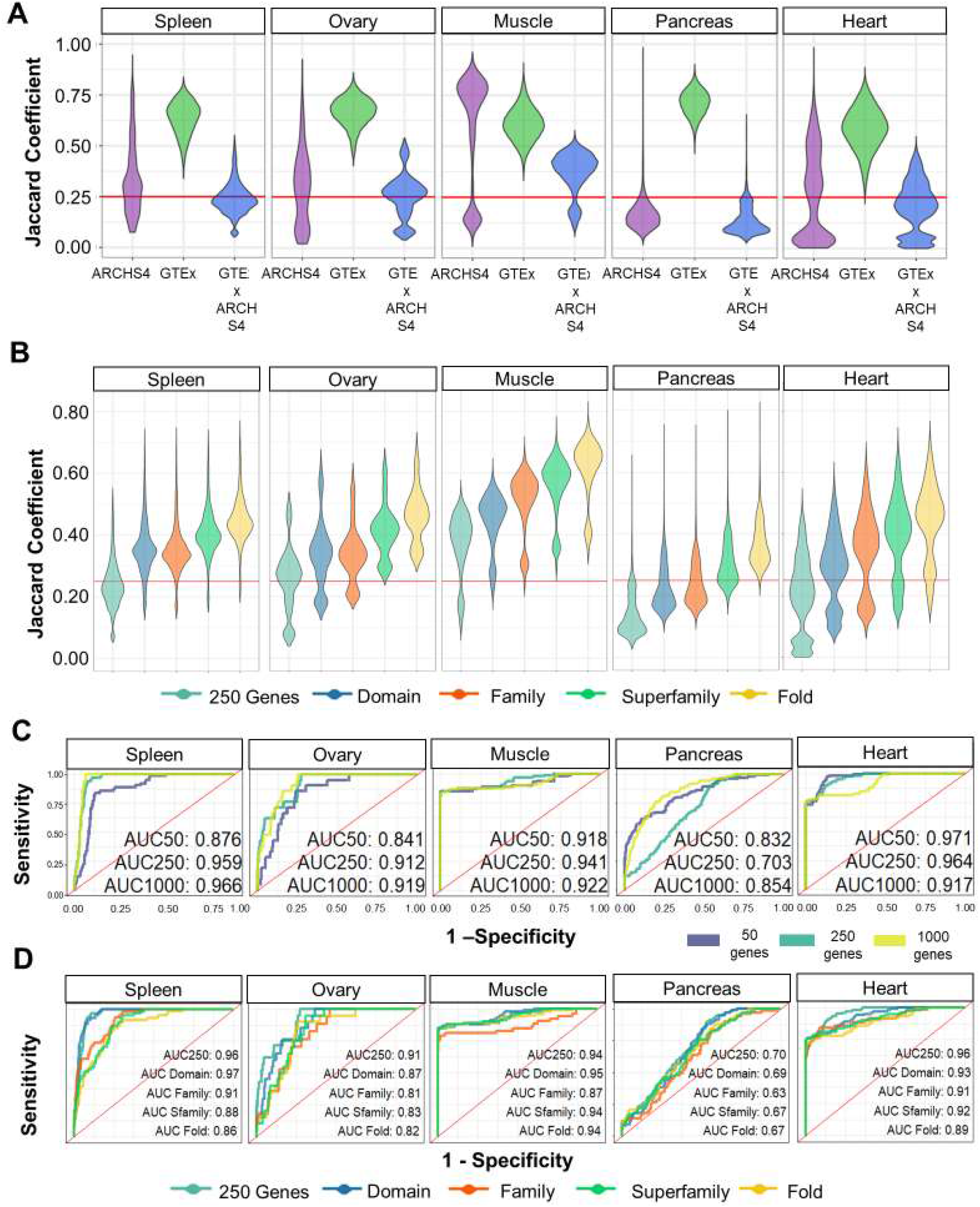
Robustness of GTEx GES using the ARCHS4 database. A) Distributions of JC values for a gene signature size of 250 for tissues within the ARCHS4 database (purple), the GTEx database (green), and across the ARCHS4 and GTEx databases (blue). Red line indicates a J_C_ = 0.25. B) Overlap of GTEx sGES with ARCHS4 signatures, across all structure levels. Red line indicates a J_C_ = 0.25. C) Predictive performance of a Random Forest model on GTEx gene sets of size 50, 250, and 1,000 highly expressed genes for predicting tissues from the ARCHS4 database, after 10-fold cross validation. D) Performance of a random forest classifier to predict ARCHS4 tissue type trained on GTEx top 250 GES or derived sGES.

In general, ARCHS4 GES consistency is much more variable across tissue types than that of GTEx GES (**Fig. 4A; purple**, and **Fig. S7**), likely because of the heterogeneity of the samples in ARCHS4. Samples in ARCHS4 can be obtained from both pathological and healthy tissues, or may characterize distinct subtypes of tissues, or may have artifacts due to differing sequencing methodologies. Importantly, measuring the consistency between ARCHS4 and GTEx by overlapping their GES alone demonstrated low JC values across most tissue types (**Fig. 4A; blue, Fig. S7**). Critically, we observed that for all tissues, sGES, at any structural level, increases the average JC overlap between GTEx and ARCHS4 signatures, and thus improves the consistency between the two datasets (**Fig. 4B, Fig. S8**).

Surprisingly, there is high predictivity of tissues from ARCHS4 using standardized models trained either on GES or sGES from GTEx (**Fig. 4C-D** and **Fig. S9-S10**). For example, AUC values for each tissue range from 0.70 (pancreas) to 0.999 (vagina) (**Fig. 4C, Fig. S9**). Importantly, decreasing GES size to 50 genes for many tissue types has significant effects on the performance of the classifier (**Fig. 4C, Fig. S9**). Several tissues such as pancreas and heart exhibit better performance using a small GES size. This indicates that much of the predictive performance of these signatures may be due to a select set of genes, rather than the signature as a whole (**Fig. 4C, Fig. S9-10**).

Using both metrics, we can identify that certain tissues such as pancreas, lung and esophagus, are not robust across GTEx and ARCHS4 due to relatively low AUC and JC values. The only potentially robust signature observed is muscle tissue, where high internal consistency within ARCHS4 and GTEx GES led to a relatively higher overlap JC distribution across the two datasets (**Fig. 4A-D**). We hypothesize that identifying and removing pathological and other atypical samples (‘outliers’) present in the ARCHS4 data will improve reproducibility across datasets. (**Fig. 4A**).

### Integration of sGES and GES enable high outlier detection

To identify potential outliers in GTEx samples, we used a neural network architecture called an autoencoder (Methods) (*30*). Autoencoders encode high dimensional data to a lower dimensional feature space that can regenerate the input of the network. The performance of an autoencoder is measured by the reconstruction error between the original inputs and the reconstructed output. Samples with high reconstruction error are often samples that are considered anomalies, or outliers, compared to the samples used to train the model.

We trained a stacked denoising autoencoder on 80% of GTEx GES. The remaining 20% of the GTEx GES was used to determine the baseline level of reconstruction error of the autoencoder (Fig. 5A; green, Fig. S11). We defined samples with reconstruction errors greater than two standard deviations of the reconstruction error (.00725) as outlier samples within GTEx. Importantly, based on this definition, very few GTEx samples can be considered outliers.

**Fig. 5:**
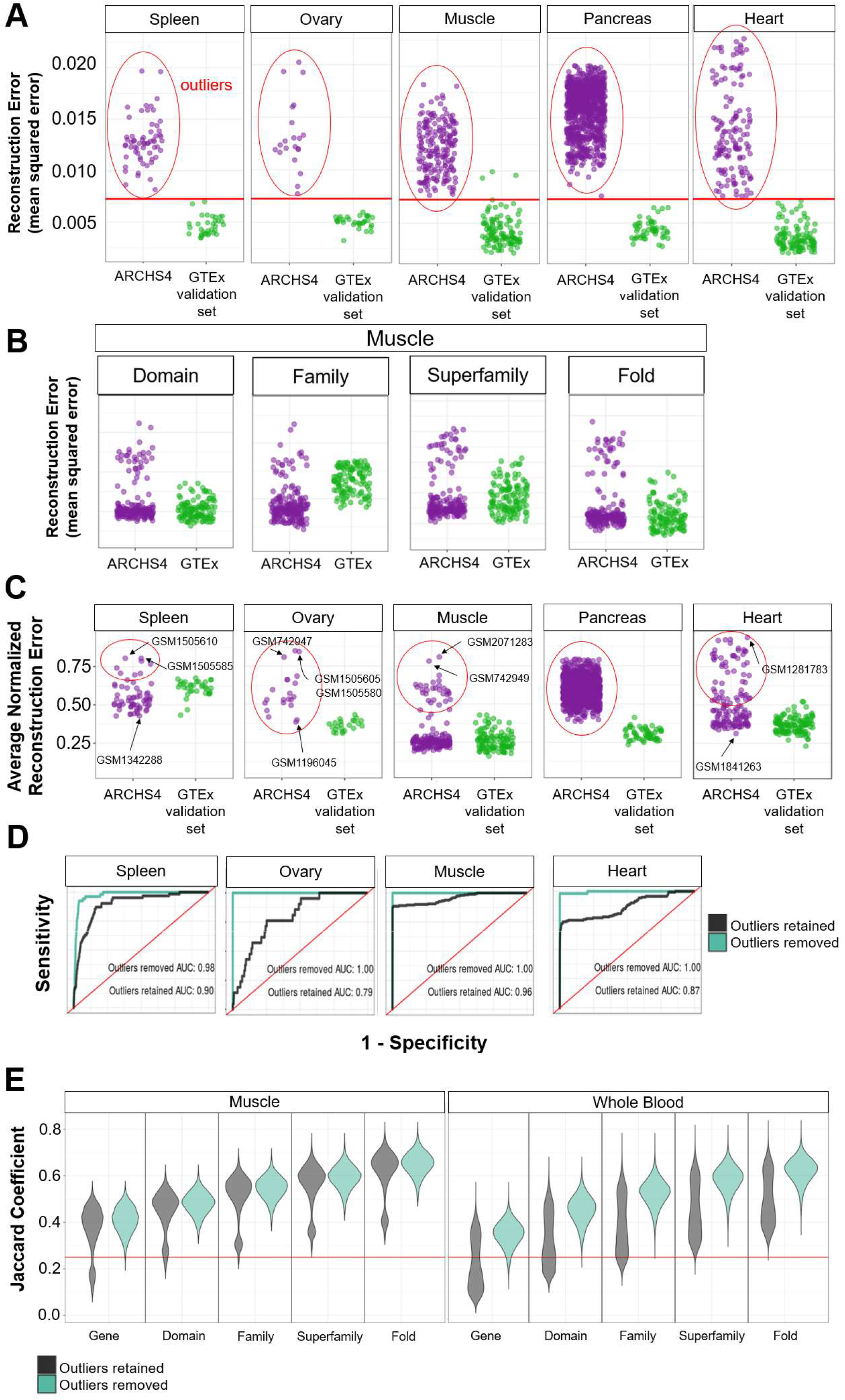
Integrated signatures enable identification of robust signatures across databases. A) Detection of outlier samples compared to GTEx gene signatures using a stacked denoising autoencoder trained to reconstruct gene signature membership from GTEx gene signatures (of size 250). Samples with high reconstruction error indicate that the sample is an outlier when compared to GTEx gene signatures. The red line indicates error values 2 standard deviations away from the mean of the distribution of errors reconstructing a validation GTEx set (error of .00725). Overlap of GTEx GES and structural signatures with ARCHS4 signatures, across tissues. B) Outlier detection using distinct structural signature levels. C) Outlier detection using integrated signatures. D) Predictive performance of GTEx GES to predict ARCHS4 tissue types, before and after outliers were removed. E) Consistency of GES and sGES of across ARCHS4 and GTEx for muscle and whole blood tissue types, before outlier removal (black) and after outlier removal (turquoise). Red line indicates a J_C_ = 0.25.

When using our trained GTEx model with ARCHS4 GES, the majority of ARCHS4 samples, within the same tissue type, are classified as ‘outliers’ (**Fig. 5A; purple, Fig. S11**). This result corroborates some results such as pancreas tissue – whose signature was demonstrated to be not robust across ARCHS4 and GTEx (**Fig. 4**). However, all ARCHS4 muscle tissue samples, which was shown to have some level of robustness (**Fig. 4**), can be considered wholly distinct datasets using this approach. Because sGES improve the overlap of JC scores across datasets and do not dramatically impact their predictivity (**Fig. 4**), we trained an autoencoder using sGES to see if outlier detection can be improved.

While we expected outlier detection to be less sensitive by ascending the structural hierarchy, as observed before (**Fig. 2-4**), surprisingly, distinct levels of structure have differing sensitivity to outliers (**Fig. 5B, Fig. S12**). For muscle tissue, the family and superfamily levels of sGES identified less outliers than those identified by domain, fold or gene level signatures. This indicates that distinct levels of the structure hierarchy characterize unique aspects of biological information present in GES.

We hypothesized that integrating GES and sGES would allow us to obtain a consensus classification of outlier vs non-outlier samples. To do so, we normalized and then averaged the reconstruction errors from autoencoders trained on GES and sGES (**Fig. 5C, Fig. S13**). Compared to either the GES (**Fig. 5A**) or sGES models (**Fig. 5B**), incorporating all signature information enables a clearer separation of true outliers in the data (**Fig. 5C, Fig. S13**). For example, this approach indicated that all pancreas tissue signatures from the ARCHS4 database can be considered outliers to GTEx pancreas signatures and thus, validated that the pancreas signature is not robust across datasets. However, for tissues such as muscle, ovary, heart, and spleen, outliers can be easily identified (**Fig. 5C**). For instance, GSM1281783, the sample with the largest reconstruction error in heart tissue in ARCHS4, characterizes dilated cardiomyopathy. Likewise, GSM2071283 (muscle) represents a sample from fetal skeletal muscle tissue, which is different from healthy adult muscle cells characterized in GTEx (**Fig. 5C**). Importantly, when identifying and removing outlier samples from ARCHS4, the predictivity and consistency of the signatures across GTEx and ARCHS4, for both GES and sGES, increased (**Fig. 5D-E, Fig. S14-S15**). We also observed increase in ARCHS4 internal GES consistency after outlier removal (**Fig. S16**).

After outlier removal, we were able to identify specific signature genes and sGES that are common across all ARCHS4 and GTEx samples (**Table S7-S8**). Table S8 shows signature genes and enriched domains, families, superfamilies, and folds seen across every ARCHS4 and GTEx whole blood samples, after outlier removal. While only two genes are consistently seen across both datasets (Actin Beta, Ferritin Light Chain), several domains (such as protein kinase domain, Immunoglobulin-like domain), families (such as C1 set domains, Pyruvate oxidase and decarboxylase PP module), superfamilies (such as EF-hand, Clathrin adaptor appendage domain), and folds (such as P-loop containing nucleoside triphosphate hydrolases, SH3-like barrel) were observed in the whole blood signature, demonstrating that sGES can illuminate additional biological information not present in GES alone.

Taken together, our results indicate that utilization and integration of both gene and protein structure information can dramatically improve the identification of outliers and enables the detection of robust expression signatures across datasets.

### sGES captures drug action on cardiomyocyte-like cell lines

We investigated if sGES alone can describe drug action on newly obtained transcriptomics data. We analyzed expression data from cardiomyocyte-like cell lines generated by the DToxS LINCS Center, to identify perturbagen specific cardiomyocyte response to specific drugs. We observed that certain over and underrepresented protein folds distinguish kinase inhibitor response from anthracycline drugs (**Fig. 6A-C**). For example, the kinase inhibitors nilotinib (NIL), regorafenib (REG), sorafenib (SOR), pazopanib (PAZ), and vemurafenib (VEM) have a characteristic underexpression of folds relating to metabolism (**Table S9**). Conversely, the anthracyclines drugs epirubicin (EPI) and doxorubicin (DOX), have characteristic overexpression of folds related to cytokine action (Table S9) as well as underexpression of folds relating to tRNA regulation such as: Proline tRNA ligase; Prolyl-tRNA synthetase; Aminoacyl-tRNA synthetases; Transmembrane ATPases; aminoacyl-tRNA synthetases; Anticodon-binding and Cortactin-binding protein (**Table S10**). Taken together, fold level sGES alone can further specify drug activity on cardiomyocytes, in addition to ranked lists of expressed genes.

**Fig. 6:**
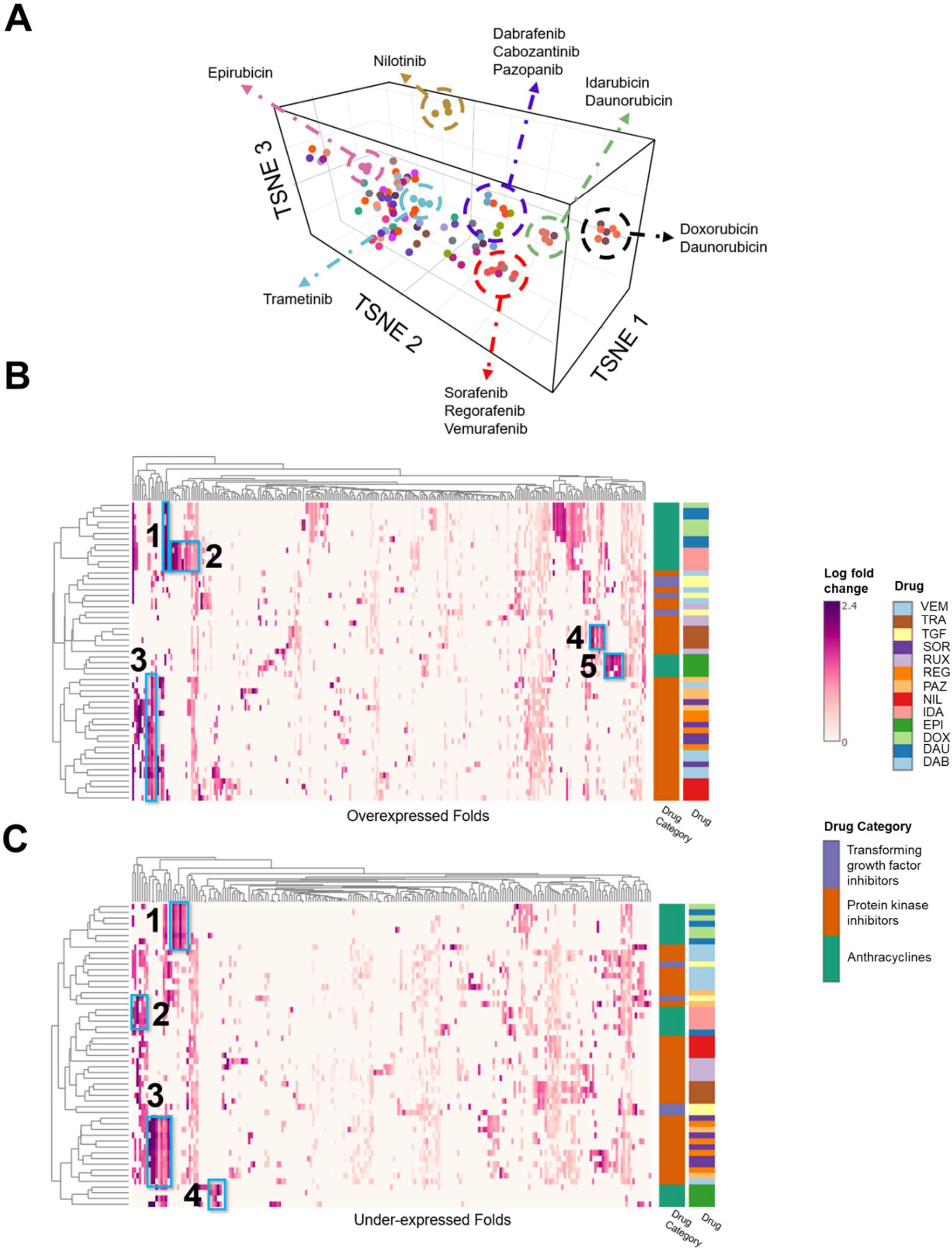
Characterization of kinase inhibitor activity using structural signatures. A) t-SNE clustering of fold signatures from distinct type of drugs on Promocell cardiomyocyte-like cell lines. Rows are labeled by Drug name, or level 3 ATC category. B) Overexpressed fold signatures for certain drugs. C) Under-expressed fold signatures for certain drugs. Distinct over and under-expressed clusters of folds are given numbers and are described in **Tables 2-3**.

## Discussion

In this study, we hypothesized that transforming gene signature space into protein structure space (e.g., domain, fold, superfamily) can characterize a robust, reproducible structural GES (sGES), and accurately define a phenotype. Additionally, integrating higher order structural features with ranked gene lists through an autoencoder, can be used to precisely identify outlier samples between distinct gene expression datasets, facilitating interoperability between experiments that use differing transcriptomic methodology. Three key findings emerge from this study. First, we define complementary metrics for evaluating the robustness of the GES: consistency, corresponds to the overlap of top ranked genes based on expression level (J_C_; **Fig. 1**); and predictivity assesses the predictive power of a phenotype using GES derived in different samples. (**Fig. 1,3-5**).

Second, we develop a new signature type termed structural gene expression signature (sGES), using features derived from various levels of protein structure (**Fig. 1**). The structural signature alone is able to characterize biological phenomena such as tissue type (**Fig. 2**). sGES overall improve the consistency of GES, while not impacting the predictive performance of signatures both within the same GES dataset and across gene expression datasets (**Fig. 3-4**). We also observed that integration of sGES and GES (using an autoencoder) facilitates the identification outliers among experimental samples enabling the filtering of unrelated samples to identify a robust expression signature and improve the reproducibility of transcriptomics analysis studies (**Fig. 5**).

Third, the structural signature was tested on multiple independent datasets, including a newly generated set of differentially expressed genes from DToxS (**Fig. 6**). This finding shows that distinct structural signatures can also be used to characterize the effects of perturbation. For example, structural signatures distinguish kinase inhibitors from anthracyclines, since anthracyclines down regulate several folds associated with tRNA regulatory factors. It has been shown that doxorubicin and its analogs bind to tRNA molecules which has been thought to contribute to their antitumor activity; however, explicit downregulation of tRNA molecules has not been previously reported, demonstrating a potential novel mechanism of anthracycline drug action (*31–33*). We expect that further investigation of sGES can lead to the identification of co-expressed structures which may reveal novel interactions between certain types proteins.

## Materials and Methods

### Computation of gene set consistency

Gene expression data was downloaded from the GTEx, version 7. For each experimental sample, each gene was sorted by expression level, in transcripts per million (TPM), and the top 50, 250, and 1,000 expressed genes were selected. For each pair of experimental samples from the same tissue subtype, the Jaccard coefficient (J_C_) was calculated to measure the overlap of GES between samples of the same tissue type. The Jaccard Coefficient JC was computed as follows:

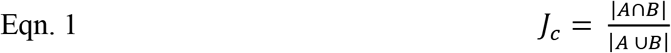

Where A and B are sets of genes names of size N (the top 50, 250, and 1,000 expressed genes). Distributions of JC for each gene set size, from selected tissues, were collected to measure robustness transcriptional signatures. A null distribution was generated by computing 1,000 bootstrap JC values between pairs of gene sets from distinct tissue types, without replacement, for gene sets of sizes 50, 250, and 1,000.

### Definition of a structural gene expression signature (sGES)

A structural signature for each gene set was determined by identifying available structural features from the encoded protein of each gene (**Fig. 1**). We defined a structural feature as a member of the structural hierarchy from the Structural Classification of Proteins extended (SCOPe) database (version 2.07). Here, structural features such as domains are categorized into families, which are categorized into superfamilies, which are further classified into distinct folds. We used HHpred (version 3.2.0) to annotate the SCOPe structural features of each protein, in the entire proteome. We used the following minimum threshold values for assigning SCOPe identifiers to proteins: length of alignment to a structure 30 residues, probability score: 50, overlap coverage: 80%, *p*-value: 1e-05, e-value: 1e-05, percent identity: 30%, coverage against template 30%.

In addition to the SCOPe hierarchy, we also obtained InterProScan (version 5.36) protein domain annotations for the human proteome, from UniProt (downloaded June 2018). For a given gene set, each structural feature was evaluated for enrichment in the gene set using a one-sided Fisher’s exact test, comparing the counts of the structural feature in the given gene set to the counts of each structural feature in the human proteome. For each gene set, a resulting structural signature is derived at the domain, family, superfamily, and fold levels, along with the log10x change, the *p*-value of enrichment (association), and the Bonferroni adjusted *p*-value (*q*-value) for each structural feature.

### A random forest algorithm for predicting tissue type

We trained a random forest classifier to predict tissue labels from gene set sizes of 50, 250, and 1,000 from GTEx expression data. We used a random forest model from the R package Ranger, for each sample from GTEx where each feature was a gene, and the value was the rank of the gene, based on the TPM observed from RNA sequencing. Genes not seen in a sample’s gene set were given a value of 0. The random forest model was trained using default parameters (mtry =20, ntree = 100). Ten-fold cross validation was used to measure performance of the GES, using a 50/50 testing-training, per tissue, split; meaning 50% of all samples, per tissue was used for either testing and training, with replacement. Receiver operator curves (ROC) were generated using the pROC package in the R programming language. Importantly, we did not perform parameter optimization for the random forest method since our goal was not optimal predictive performance, but rather to determine the baseline predictive performance of GES.

### Predictivity of random forest models on ARCHS4 gene sets

Gene expression datasets of the following tissues: adipose, brain, colon, esophagus, fallopian tube, heart, kidney, liver, lung, muscle, nerve, ovary, pancreas, prostate, small intestine, spleen, stomach, testis, thyroid, uterus, vagina, and whole blood, were downloaded from the ARCHS4 database (March 2019). The top 250 overexpressed genes from each of the samples of the tissue types were obtained by ranking the read counts of the Kalisto aligned expression data. We then predicted ARCHS4 tissue class using the random forest model trained on GTEx GES.

### Signature consistency

A structural signature was obtained for gene sets of sizes 50 to 1,000, across all GTEx tissue types. Pairwise Jaccard coefficients (*J_C_*; Eqn. 1) were then computed between structural signatures of the same gene set size, and the same tissue. The median *J_C_* at each gene set size per tissue defined the overall consistency.

### Clustering GTEx samples

For each tissue sample, the log10X change for each structure in the structural signature derived at 250 genes was used as input for t-distributed Stochastic Network Embedding (t-SNE) using the Rtsne package, using default perplexity (*28*) settings and was run for 1,000 iterations. For tissues where a structure was not observed, a value of 0 was used.

### sGES predictivity

As described above for GES, 10x cross validation was performed for predicting GTEx tissues class from the *p*-value of association of each structural feature in the signature. Structural signatures were generated from the ARCHS4 gene signature set and were used to validate the performance of the random forest classifier trained on GTEx structural signatures.

### Integration of GES and sGES

A stacked denoising autoencoder was used to embed structural signatures into a lower dimensionality matrix. We utilized a typical symmetrical autoencoder architecture of 3 dense (fully connected) encoding and decoding layers with 100, 50, 25 neurons and a bottleneck layer of 10 neurons using the Keras package in R. Each layer’s activation function was set to ‘relu’, except for the final layer whose activation function was set to ‘sigmoid’. We used the mean squared error between the input and output layers as the loss function for the model, ran the autoencoder for 50 epochs and utilized the ‘adam’ optimizer to update network weights. For each of the structural layers the bottleneck layer was selected and combined into a flattened matrix.

### Interoperability of GTEx and ARCHS4 GES

We then used two simple neural network models to predict 1) ARCHS4 tissue classes trained on the integrated signature of GTEx data, and 2) GTEx tissue classes from ARCHS4 integrated signatures to investigate interoperability of the two datasets. The neural network architecture is as follows: 3 densely connected hidden layers of 100 neurons each using the Keras package in R. The input layer and the first two hidden layers utilized the ‘relu’ activation function, while the final hidden layer used the ‘softmax’ activation function. The neural network used the ‘adam’ optimizer, and the ‘categorical_crossentropy’ loss function since the output layer consisted of 22 tissue categories.

### Experimental protocols for cell culture, drug treatment and transcriptomics

Details of the experimental protocols for cell culture, drug treatment and transcriptomics have been described as step-by-step standard operating procedures for the various experiments available on www.dtoxs.org.

### Classification of Promocell Cardiomyocyte cell lines

For each control Promocell cardiomyocyte sample that were not exposed to a perturbagen, a set of 250 top overexpressed genes were obtained. Structural signatures were generated and plotted against a t-SNE of GTEx samples, using the log10X change of each structural feature. Pairwise Euclidean distances were taken between each control Promocell sample and all other samples in GTEx to determine the tissue type Promocells were most similar to.

### Processing and exploratory analysis of gene expression data

The median log-transformed gene expression fold-change value was calculated across all cell lines for each individual small molecule drug. The resulting matrix of gene fold change values by drugs was used for the regression analysis. To obtain insight in the general patterns present in this drug-perturbed transcriptomics dataset, we generated rankings of the top 500 genes for each drug, by their absolute mean fold change value, i.e. whether positive or negative. For each of these drug-associated rankings we determined the frequency of these changes being also present in the ranking of other drugs, e.g. the similarity in genes present in the top 250 gene lists for each drug. This was visualized using the Jc, and by plotting the most highly drug-connected genes against the associated drugs. Principal component analysis for the first 3 principal components on the absolute mean fold-change values for each drug was performed to further assess similarity between drugs in their gene expression values.

### Structural characterization of DEGs from perturbagen studies

For each experimental sample from the DToxS set, the top 250 DEGs were obtained by ranking the observed *p*-value for each gene. Structural enrichment was performed for all DEGs combined, only overexpressed genes (by positive log10x change) or only underexpressed genes (by negative log10x change). The log10x change of each structure in the structural signature of the combined gene set was used for t-SNE clustering, where structures that were unseen for a given gene set were set to 0. Each drug is colored by their level 4 Anatomic Therapeutic Code (ATC), if available. Otherwise, drugs were manually assigned to an ATC code based on the known target of the drug tested.

### Clustering of kinase inhibitors

Selected kinase inhibitors were hierarchically clustered based on the log10x change of each structural feature from over- and under expressed gene sets using the Ward method from the hclust method in the R programming language.

## General

We would like to thank the Alexander Lachmann and Avi Ma’ayan (both Mount Sinai), and Burkhard Rost (Technical University Munich) for their insightful discussions.

## Author contributions

R.R., R.I. and A.S. conceptualized the study. R.R. curated the data, performed the formal analysis, developed the methodology for structural signatures, trained and validated the machine learning models used in the analysis. Y.X. performed the transcriptomics analysis of the Promocell data. E.S, M.B, and E.A provided critical analyses and insight into study design. R.R and A.S. wrote the analysis. R.R, A.S, E.S., M.B., E.A., R.I., J.H., J.G.C.H edited and reviewed the analysis. R.I and A.S provided funding for the analysis.

## Data and materials availability

All data is available on Github.

