## Supplemental Figures and Tables for "Protein structure-based gene expression signatures"

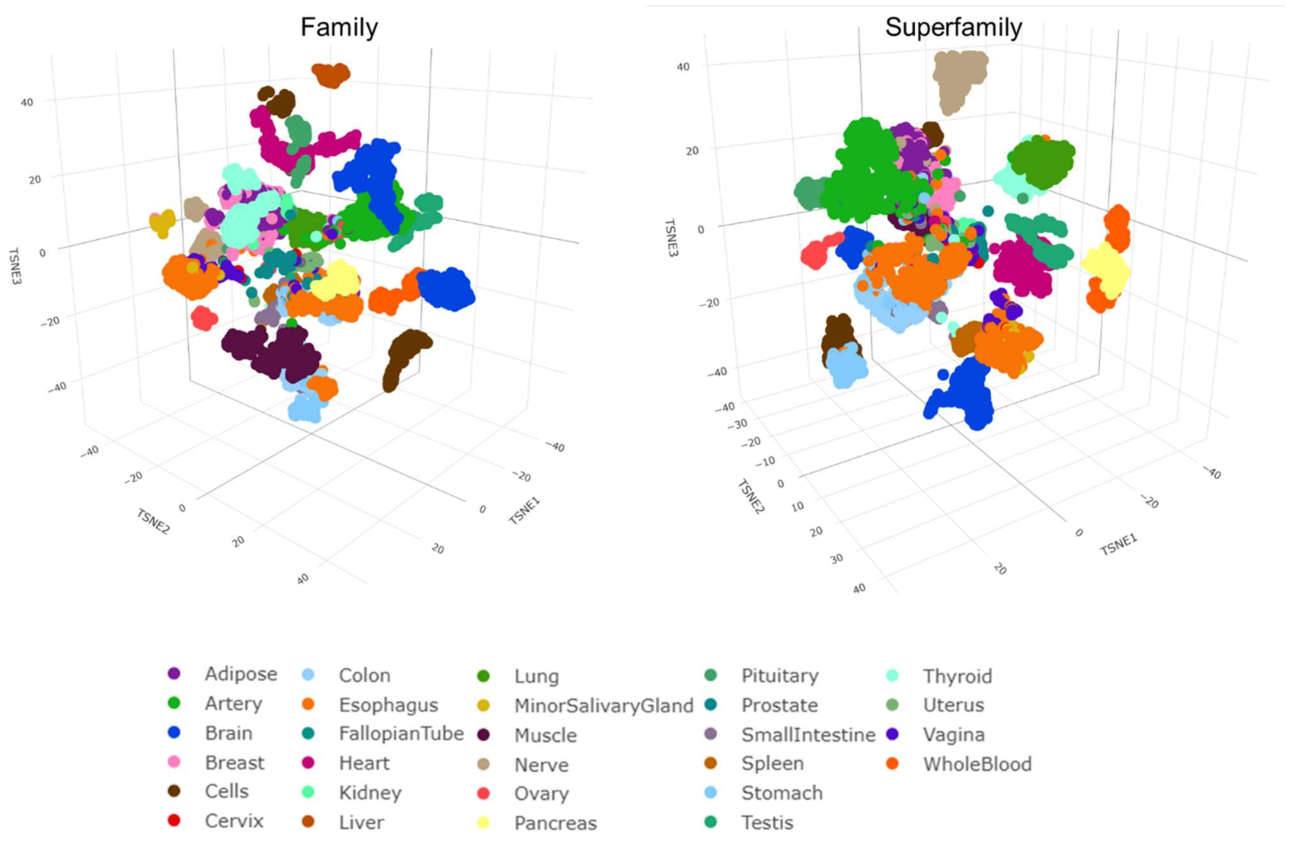


**Fig. S1: Protein structure enrichment clusters tissue-specific gene expression**. The top 250 highest expressed genes from GTEx (in terms of transcripts per million) were obtained. sGES were then derived from the GES, and tissue samples were clustered by using t-SNE based on the presence or absence of structural features at the family and superfamily levels. Each sample is colored by tissue type.

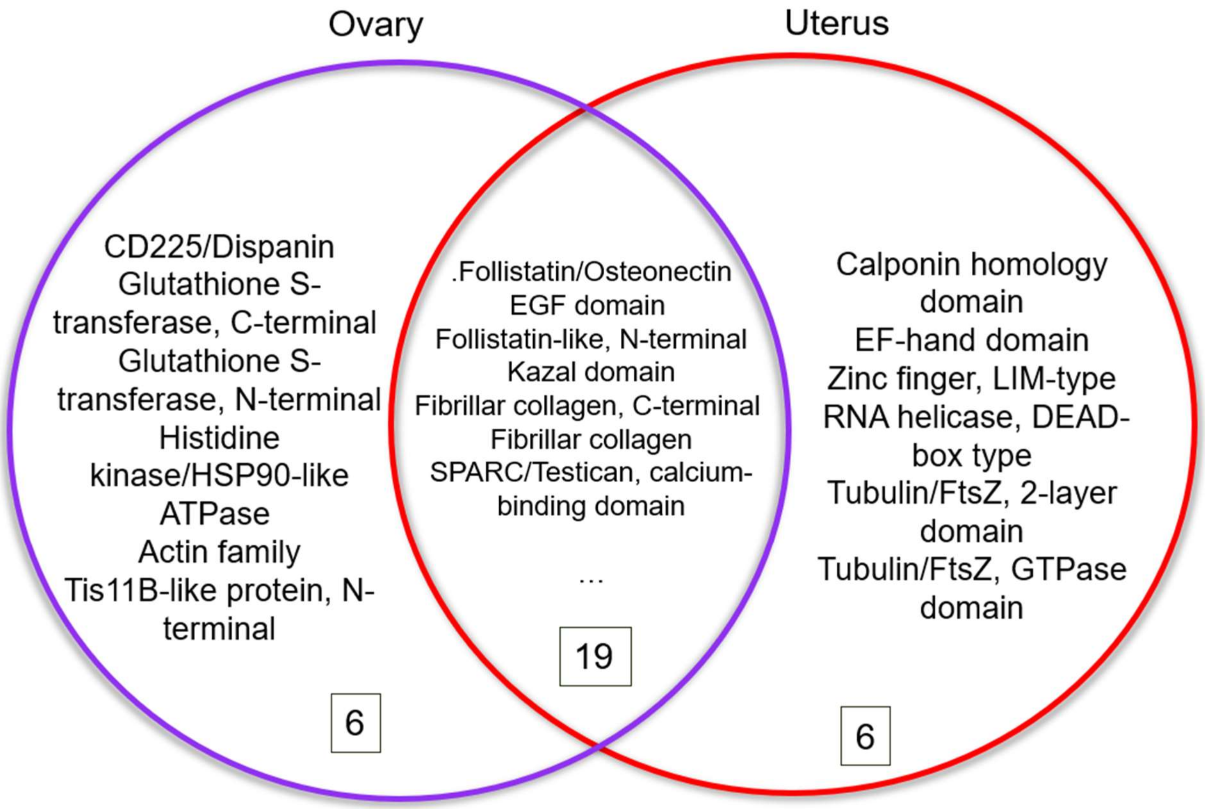


**Fig. S2:** Overlaps of overrepresented domains between Uterus and Ovary tissue types. Values indicate number of members in set.

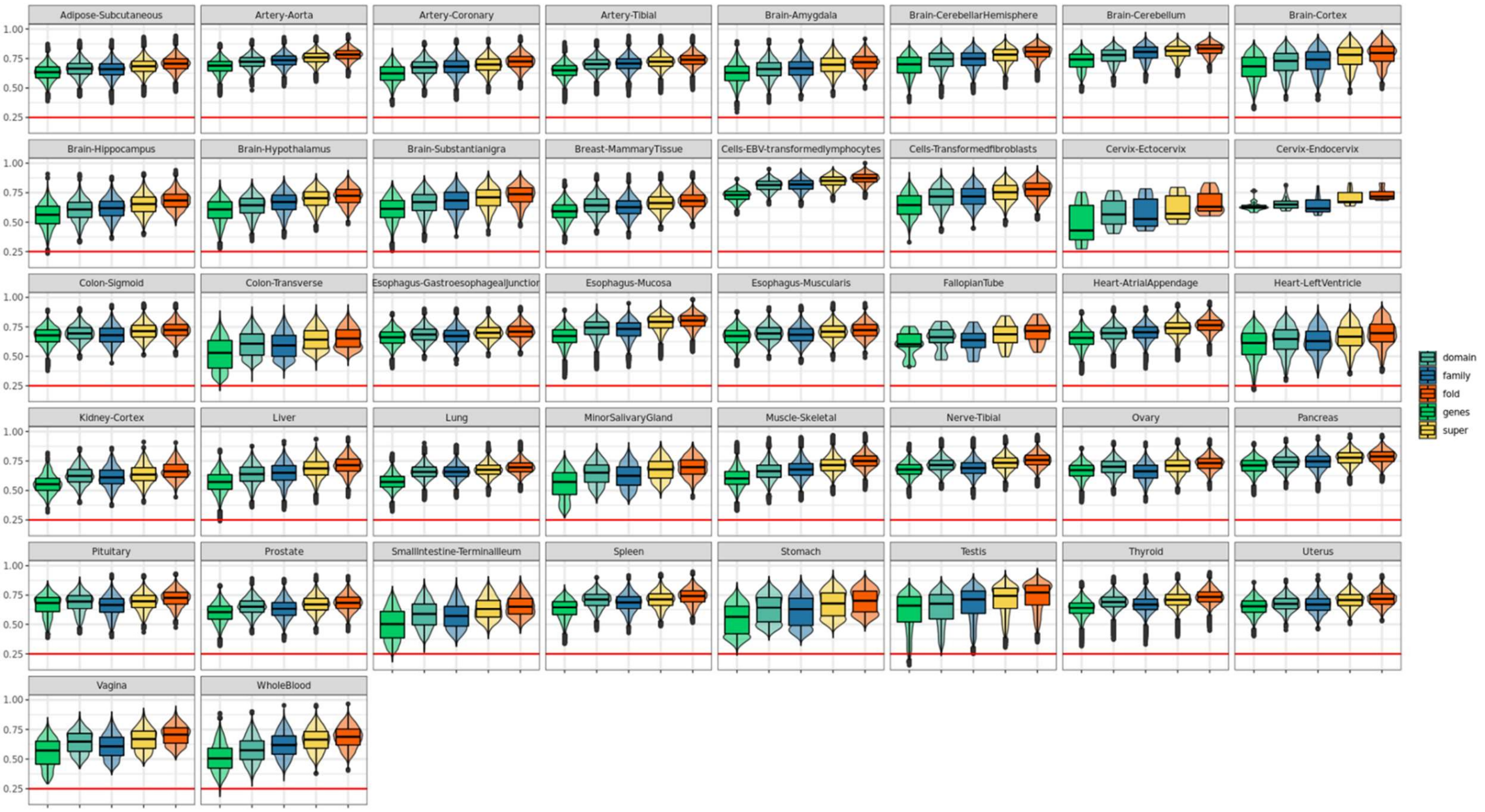


**Fig. S3:** Higher order structural signatures increases signature stability across tissues. 'Top' refers to top 250 genes, 'super' refers to superfamily structural signature level. Box plots are colored by signature type.

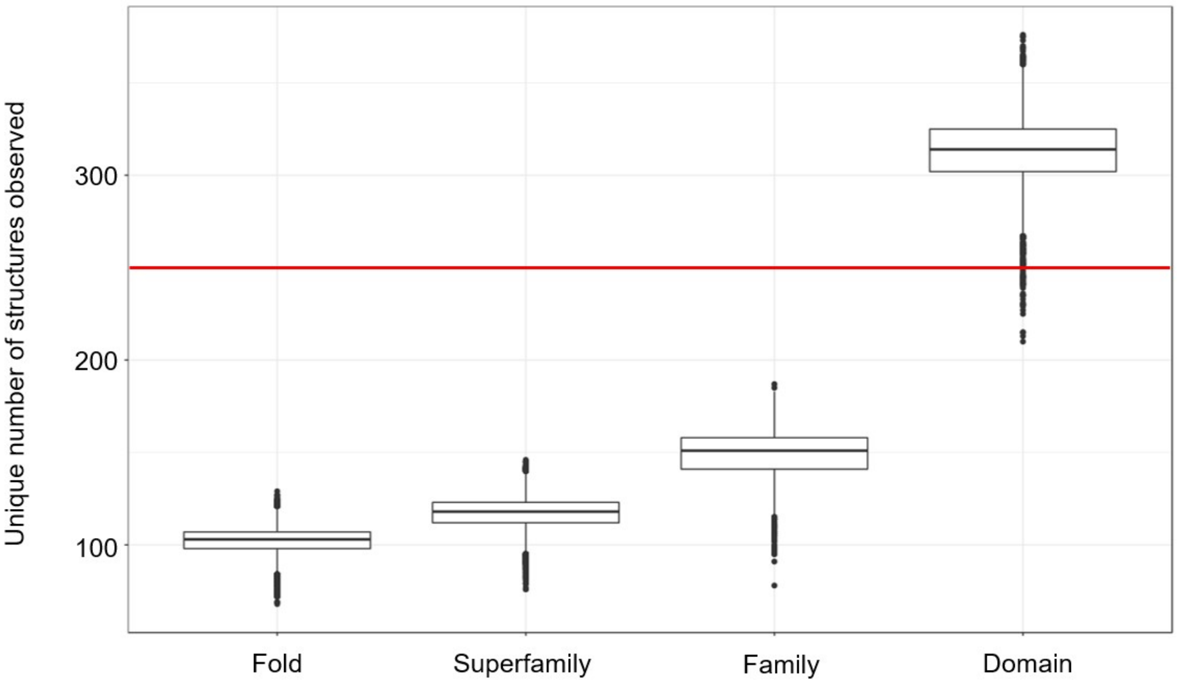


**Fig. S4**: Unique number of structures at each structural level, across all GTEx tissue samples, observed for gene signature size of 250. Red line delineates a size of 250.

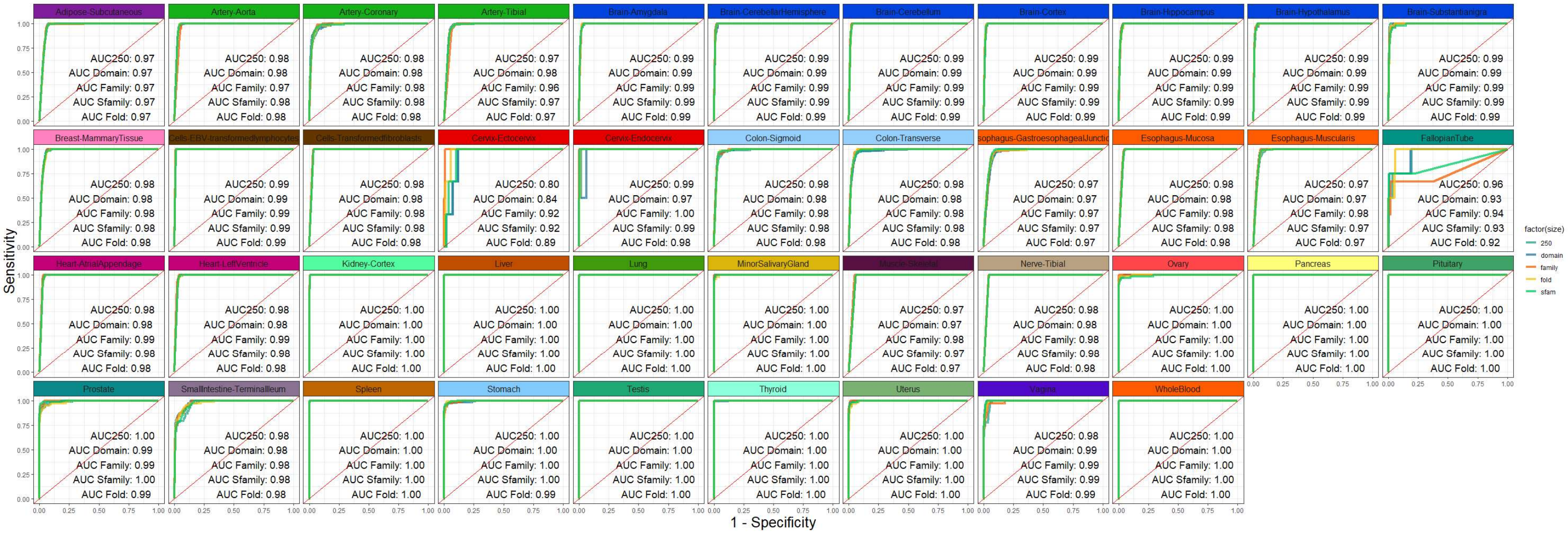


**Fig. S5.** Structural signatures are predictive of tissue type. ROC curves are shown for predicting GTEx tissue type from gene signature size 250 and at each structural signature level (fold, family, superfamily, domain) for all tissue types in GTEx.

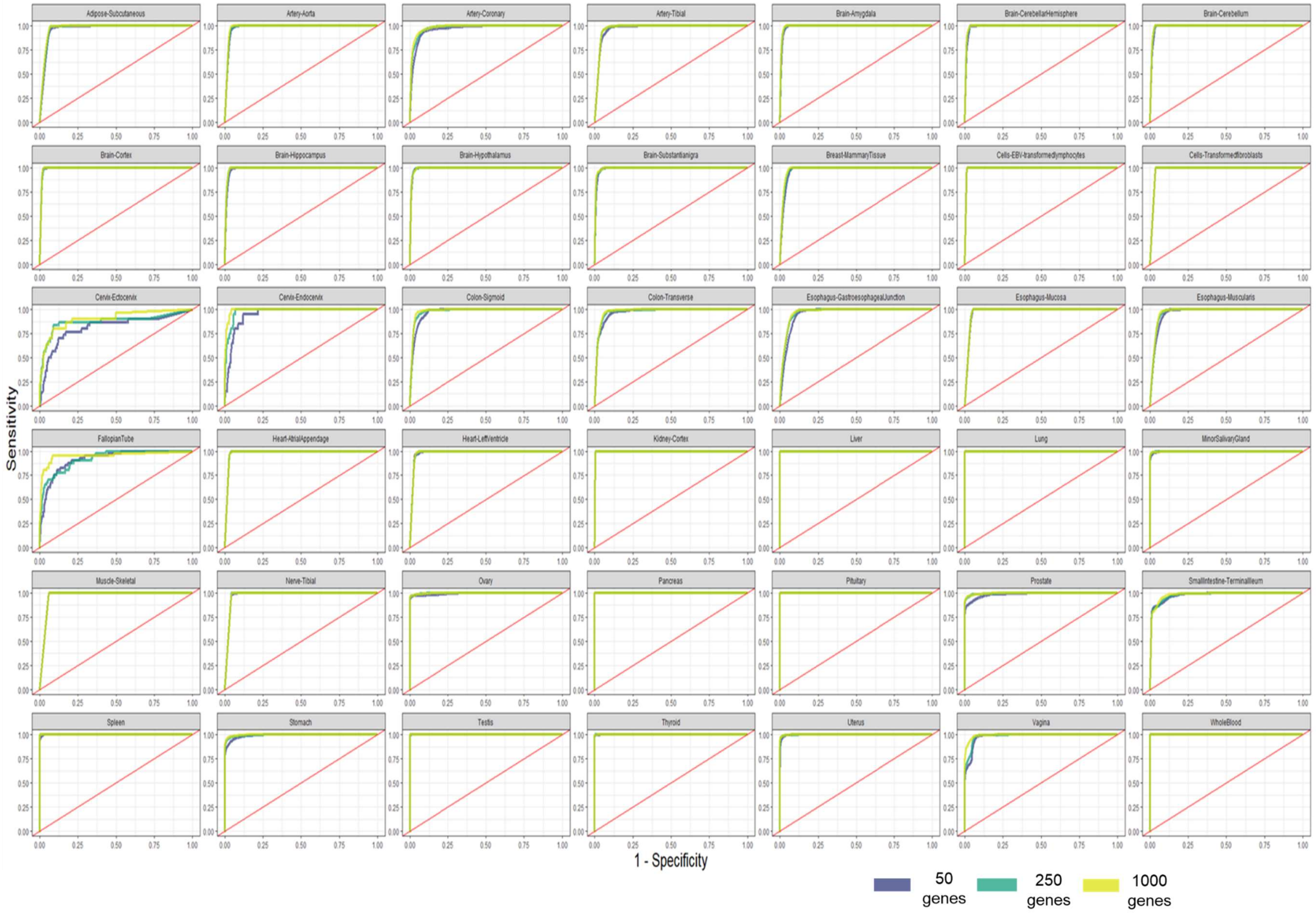
Adipose-Subcutaneous
Artery-Aorta
Artery-Coronary
Artery-Tibial
Brain-Amygdala
Brain-CerebellarHemisphere
Brain-Cerebellum
Brain-Cortex
Brain-Hippocampus
Brain-Hypothalamus
Brain-Substantianigra
Breast-MammaryTissue
Cells-EBV-transformedlymphocytes
Cells-Transformedfibroblasts
Cervix-Ectocervix
Cervix-Endocervix
Colon-Sigmoid
Colon-Transverse
Esophagus-GastroesophagealJunction
Esophagus-Mucosa
Esophagus-Muscularis
FallopianTube
Heart-AtrialAppendage
Heart-LeftVentricle
Kidney-Cortex
Liver
Lung
MinorSalivaryGland
Muscle-Skeletal
Nerve-Tibial
Ovary
Pancreas
Pituitary
Prostate
SmallIntestine-TerminalIleum
Spleen
Stomach
Testis
Thyroid
Uterus
Vagina
WholeBlood
Sensitivity
1 - Specificity
50 genes
250 genes
1000 genes

**Fig. S6.** ROC Performance of a standardized random forest classifier to predict GTEx tissue type using GES sizes of 50 (purple), 250 (green) and 1000 (yellow) genes.

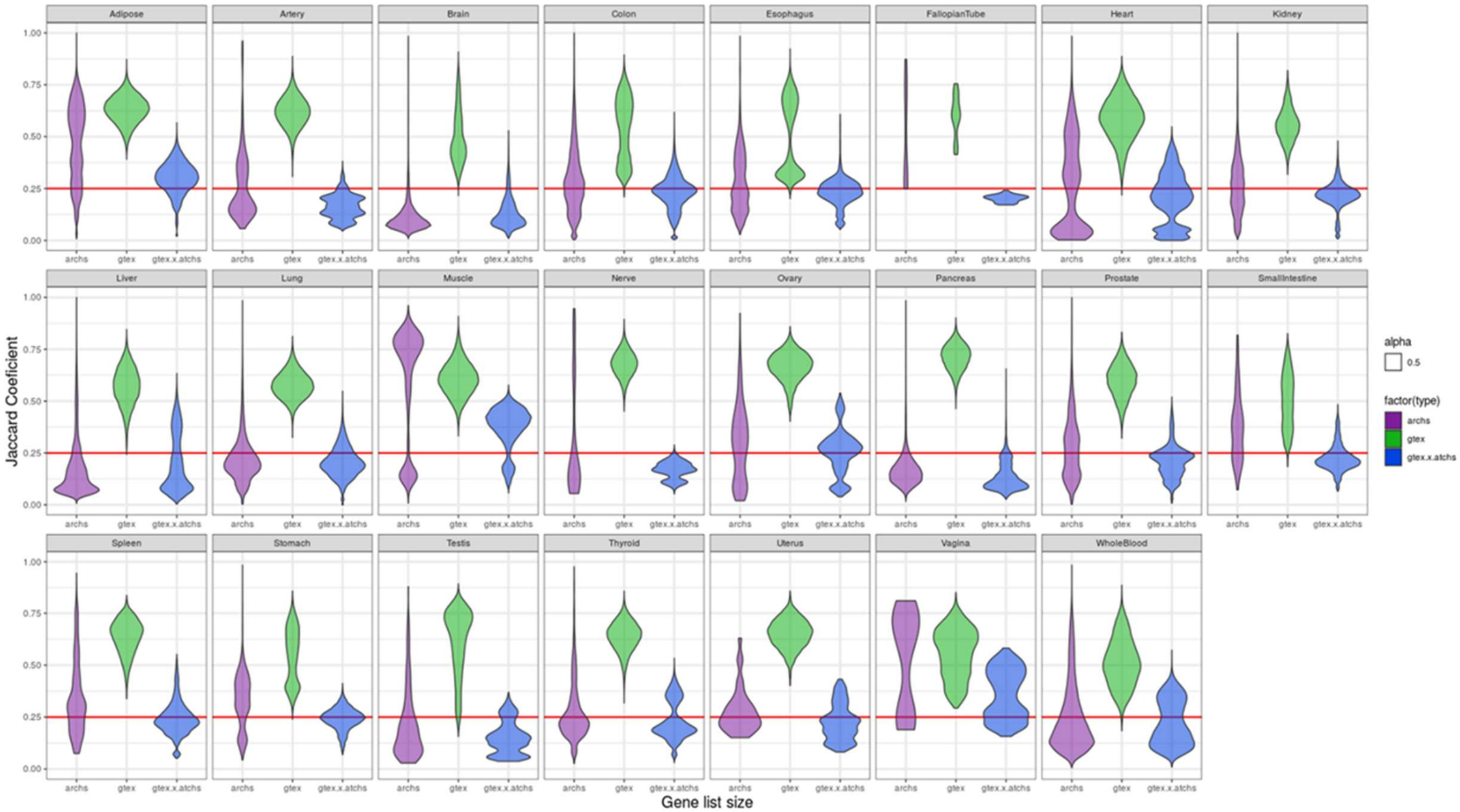


**Fig. S7:** Overlap distributions of JC values across all tissue types. Purple refers to GTEx internal consistency. Green refers to ARCHS internal consistency. Blue refers to ARCHS4 and GTEx cross consistency.

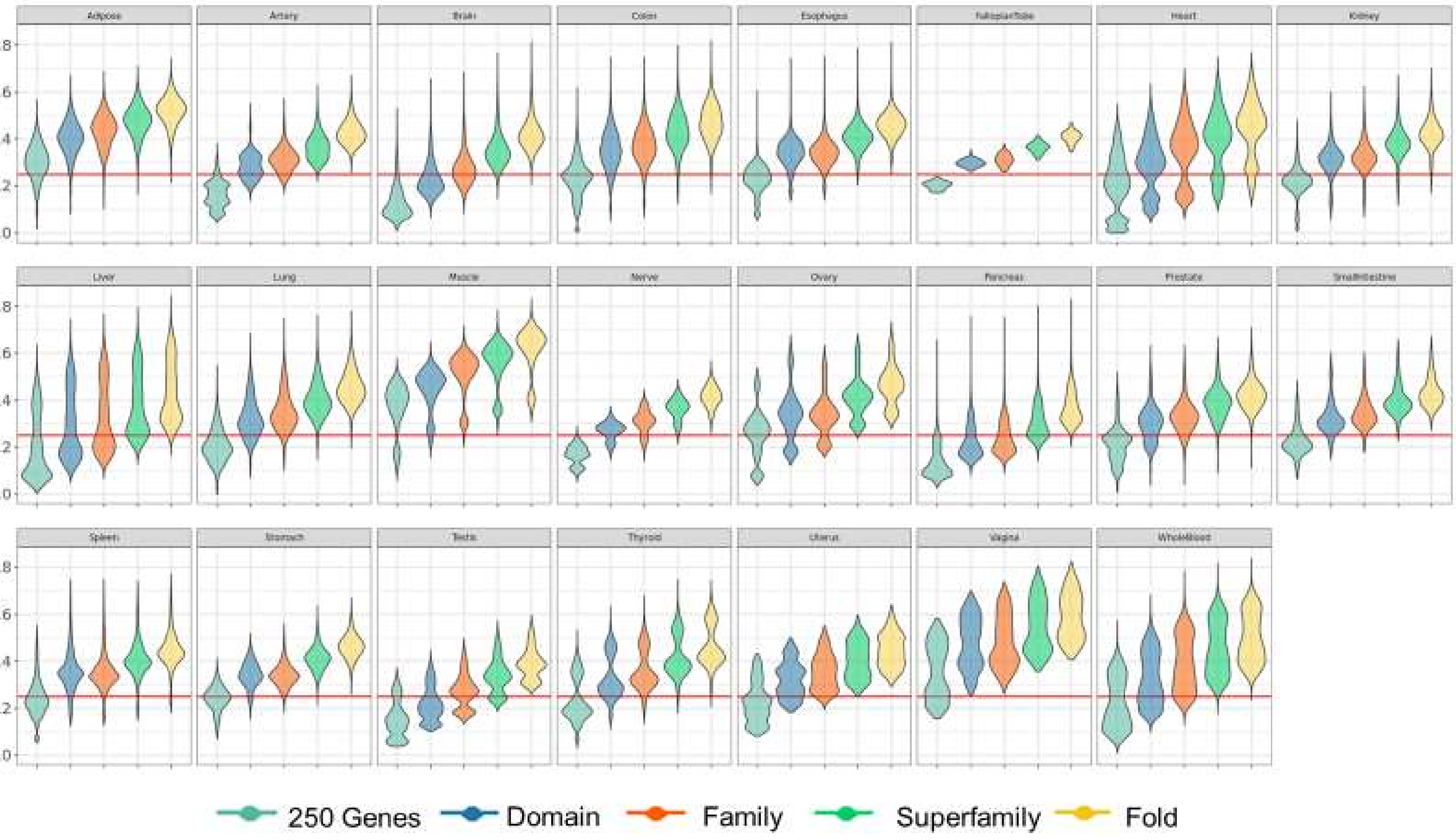


**Fig. S8:** Overlap distributions across GTEx and ARCHS4 of $J_C$ values across all tissue types using sGES.

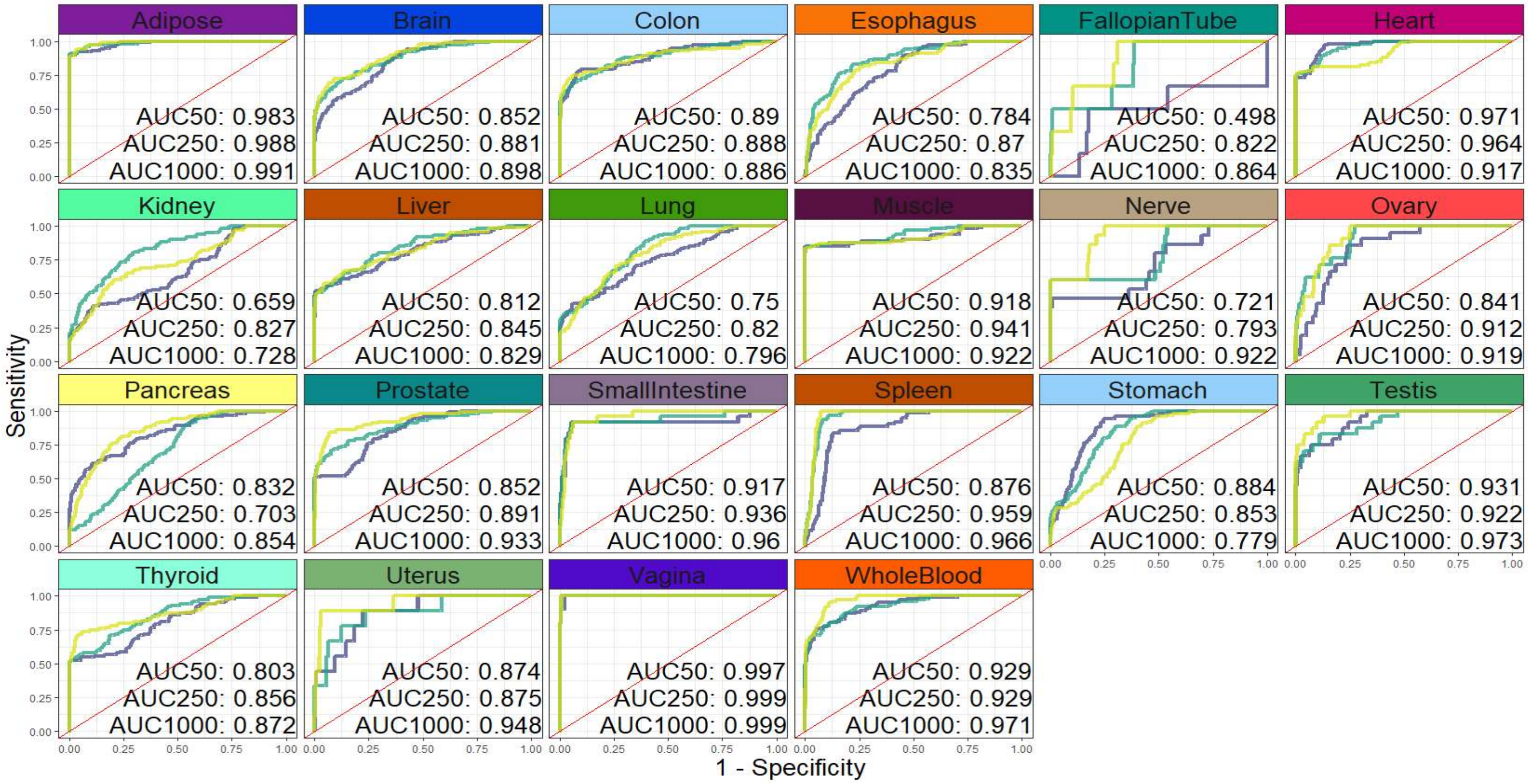


**Fig. S9:** Validation of GTEx signature model against ARCHS4 for all tissues, at distinct gene signature sizes, including 50 (purple), 250 (green), and 1,000 (yellow) genes.

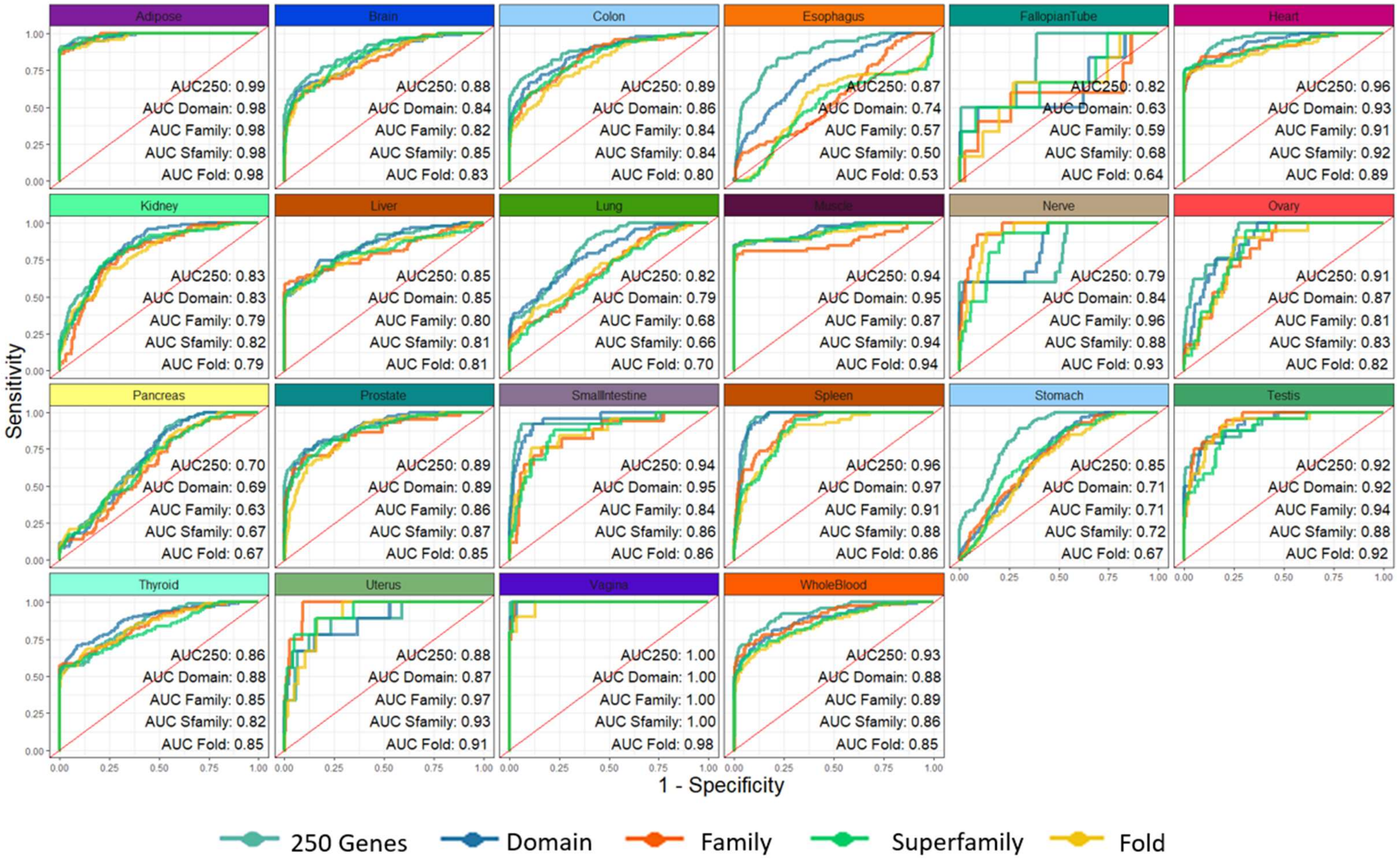


**Fig. S10**: Predictive performance of gene signatures (of size 250 gene) and sGES trained on GTEx to predict ARCHS4 tissues.

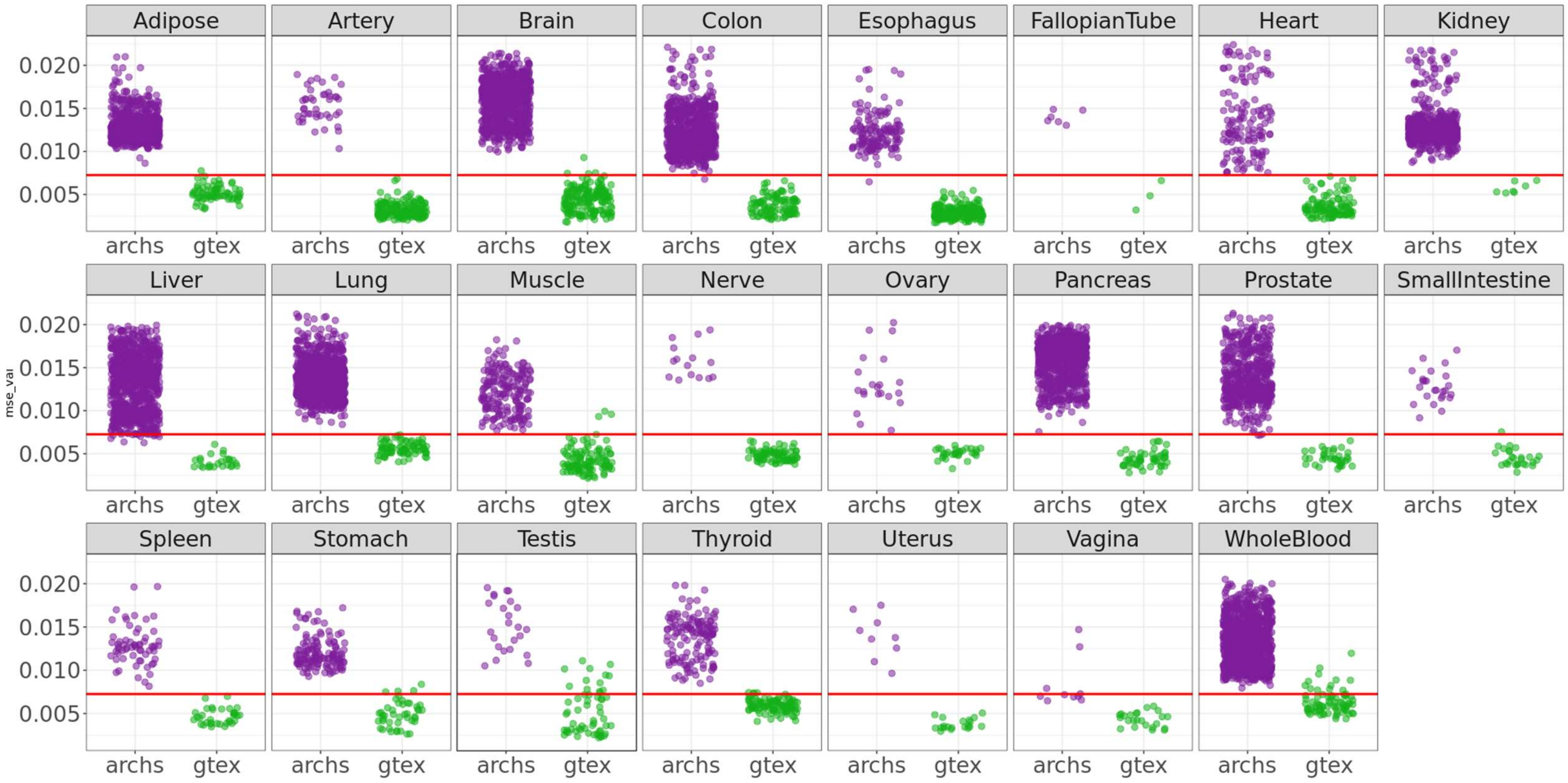


**Fig. S11:** Detection of outlier samples compared to GTEx gene signatures using a stacked denoising autoencoder trained to reconstruct gene signature membership from GTEx gene signatures (of size 250). Samples with high reconstruction error indicate that the sample is an outlier when compared to GTEx gene signatures. The red line indicates error values 4 2 standard deviations away from the mean of the distribution of errors reconstructing a validation GTEx set (error of .00725).

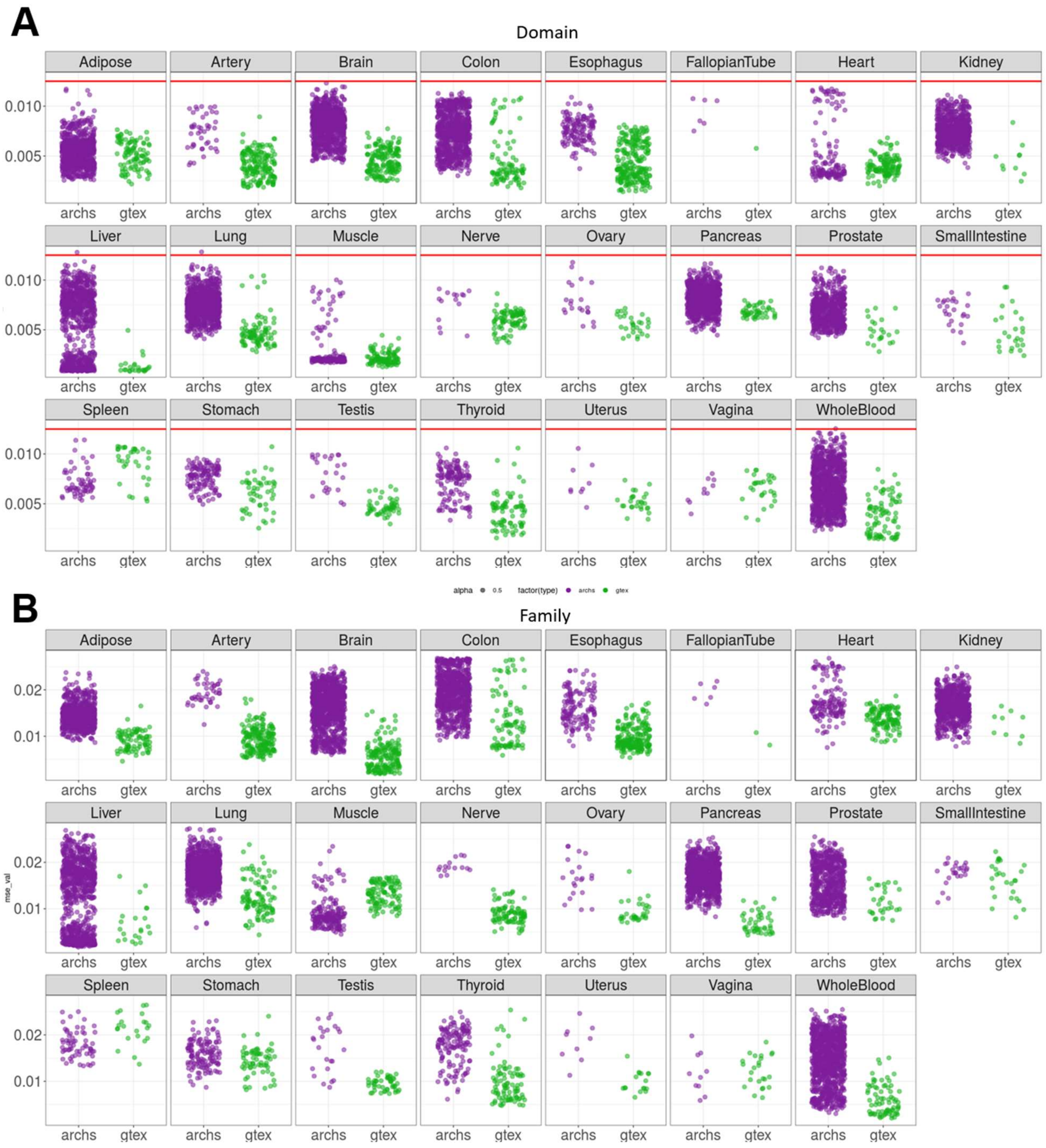
A
Domain
Adipose
Artery
Brain
Colon
Esophagus
FallopianTube
Heart
Kidney
Liver
Lung
Muscle
Nerve
Ovary
Pancreas
Prostate
SmallIntestine
Spleen
Stomach
Testis
Thyroid
Uterus
Vagina
WholeBlood
0.010
0.005
archs
gtex
alpha 0.5 factor(type) archs gtex
B
Family
0.02
0.01
mse_val

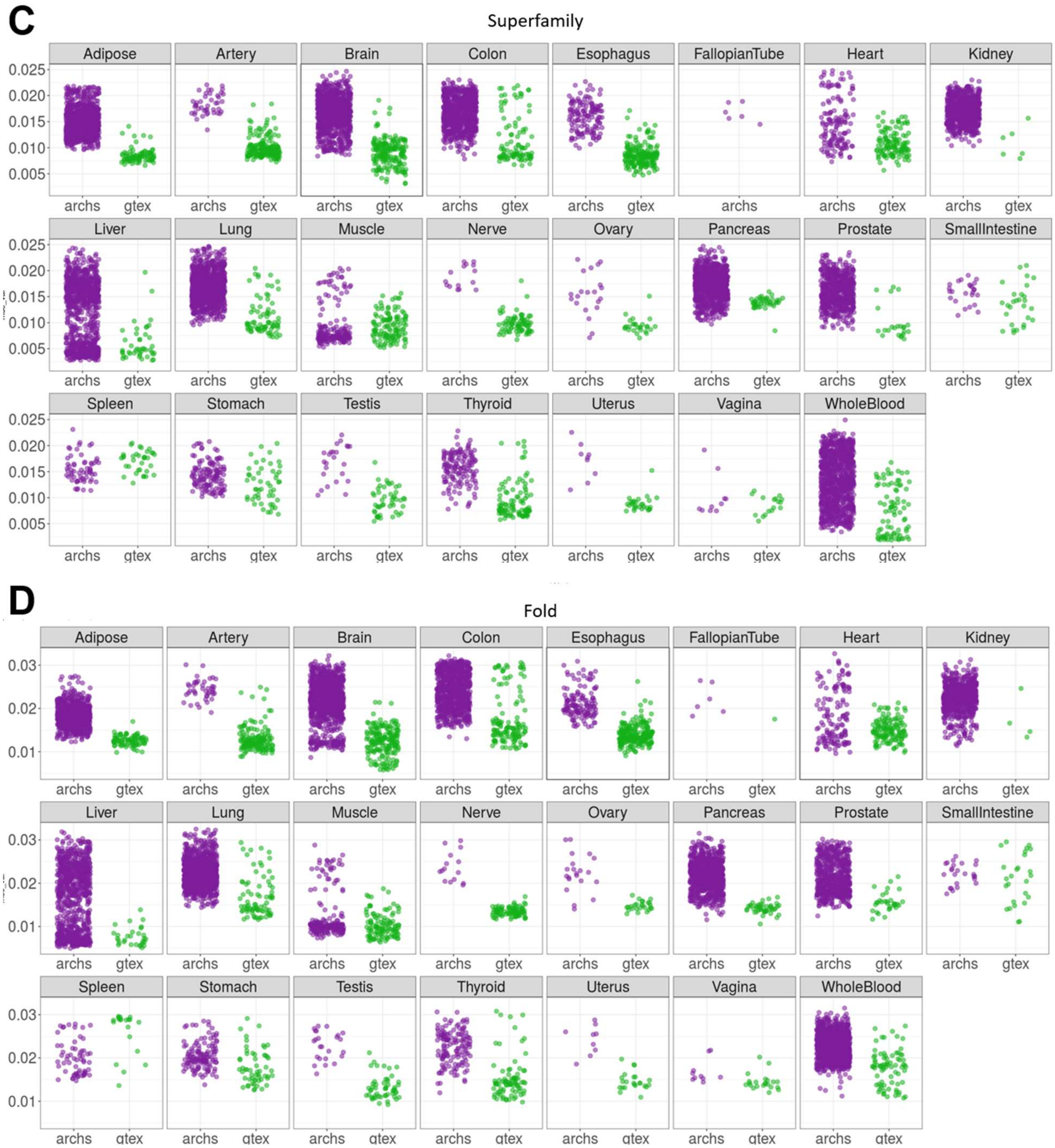
C
Superfamily
Adipose
Artery
Brain
Colon
Esophagus
FallopianTube
Heart
Kidney
Liver
Lung
Muscle
Nerve
Ovary
Pancreas
Prostate
SmallIntestine
Spleen
Stomach
Testis
Thyroid
Uterus
Vagina
WholeBlood
0.025
0.020
0.015
0.010
0.005
archs
gtex
D
Fold
0.03
0.02
0.01

**Fig. S12:** Reconstruction errors of ARCHS4 (purple) and GTEx (green) samples using an autoencoder trained on GTEx at each structural level: A) domain, B) Family, C) Superfamily, D) Fold.

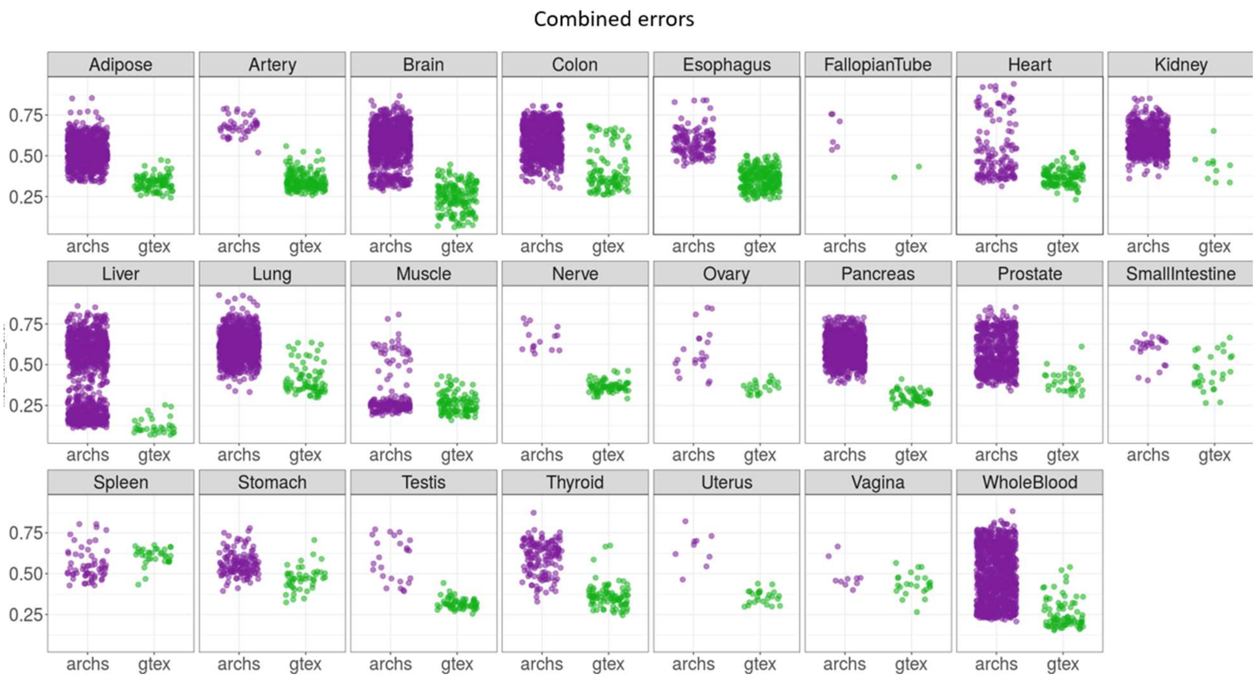


**Fig. S13:** Combined GES and structural signature reconstruction errors of ARCHS4 (purple) and GTEx (green) tissues types trained on GTEx, across all tissue types.

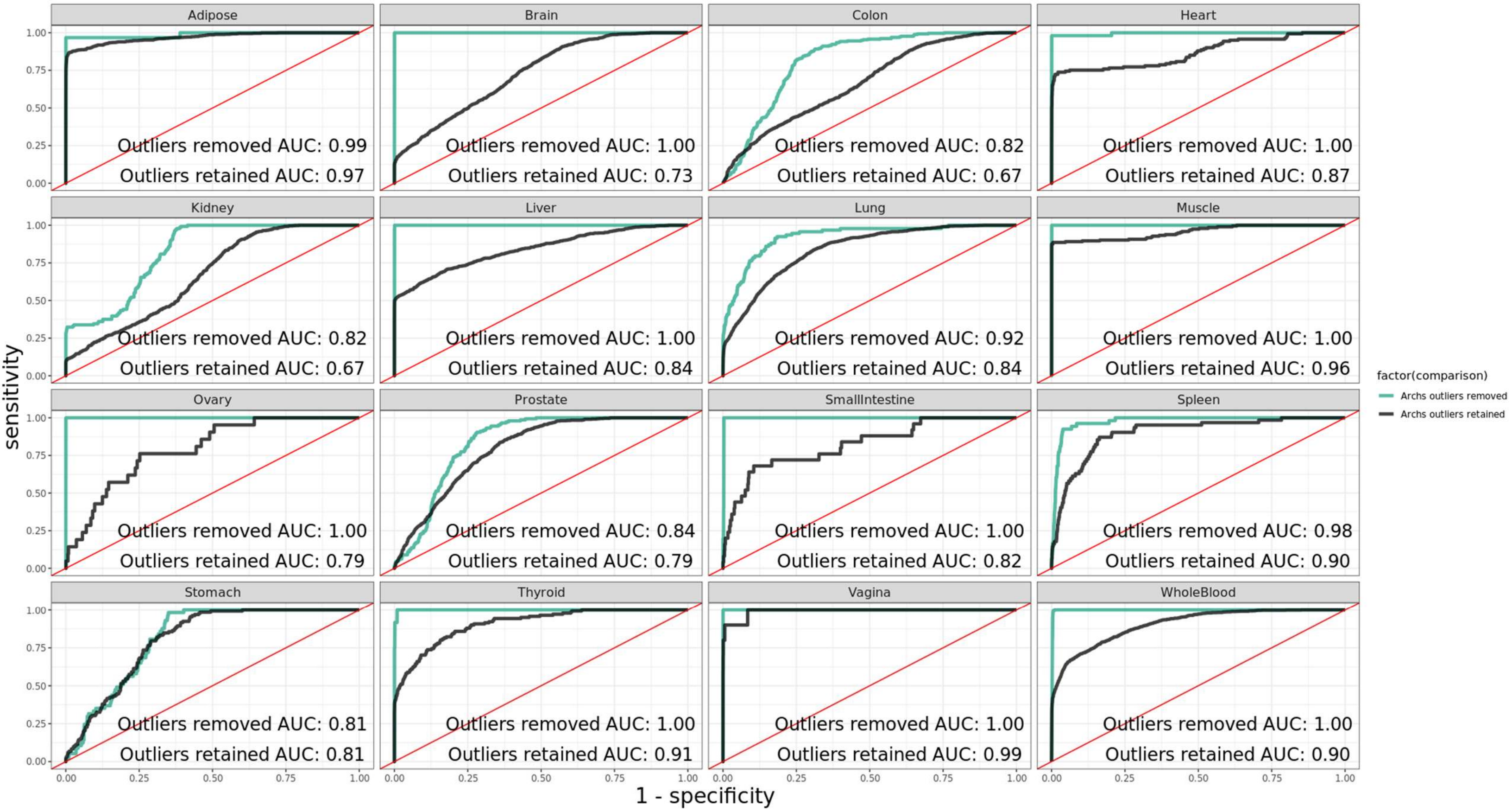


**Fig. S14:** Predicting ARCHS4 tissue type from GTEx trained models, where ARCHS4 outliers are included (black) vs removed (green), across all overlapping tissue types.

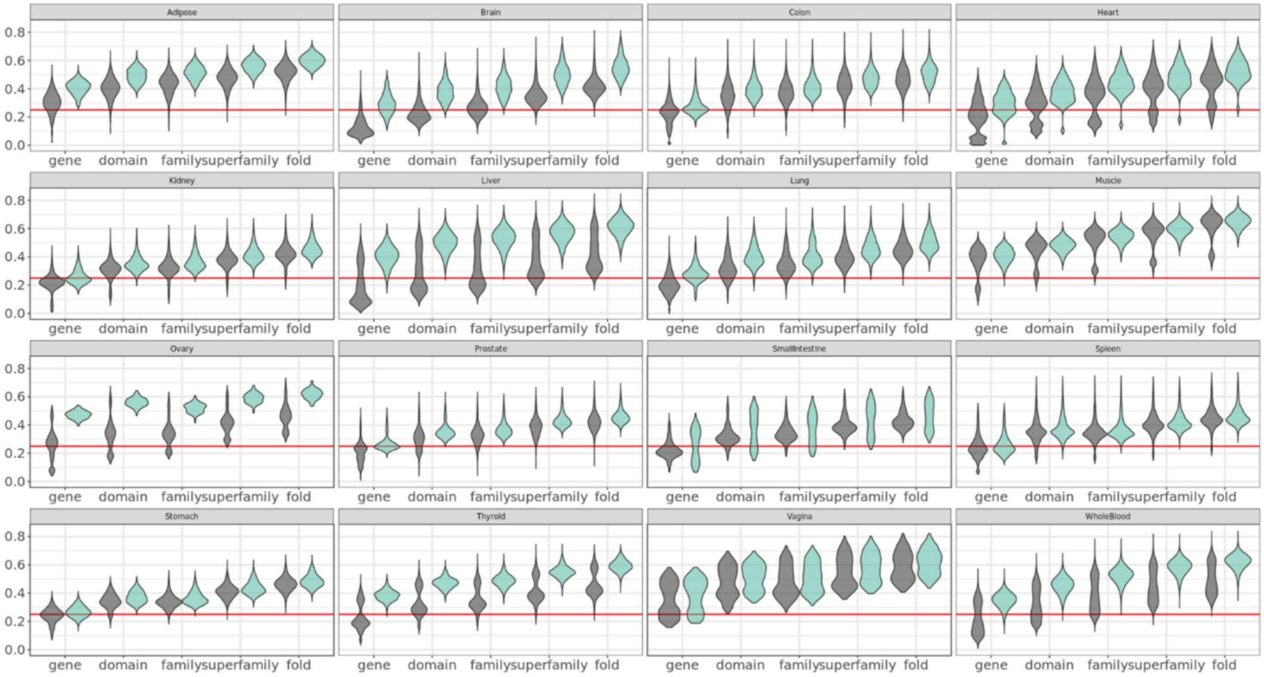


**Fig. S15:** Consistency of GES and sGES of across ARCHS4 and GTEx for all tissue types, before outlier removal (black) and after outlier removal (turquoise). Red line indicates a $J_C = 0.25$.

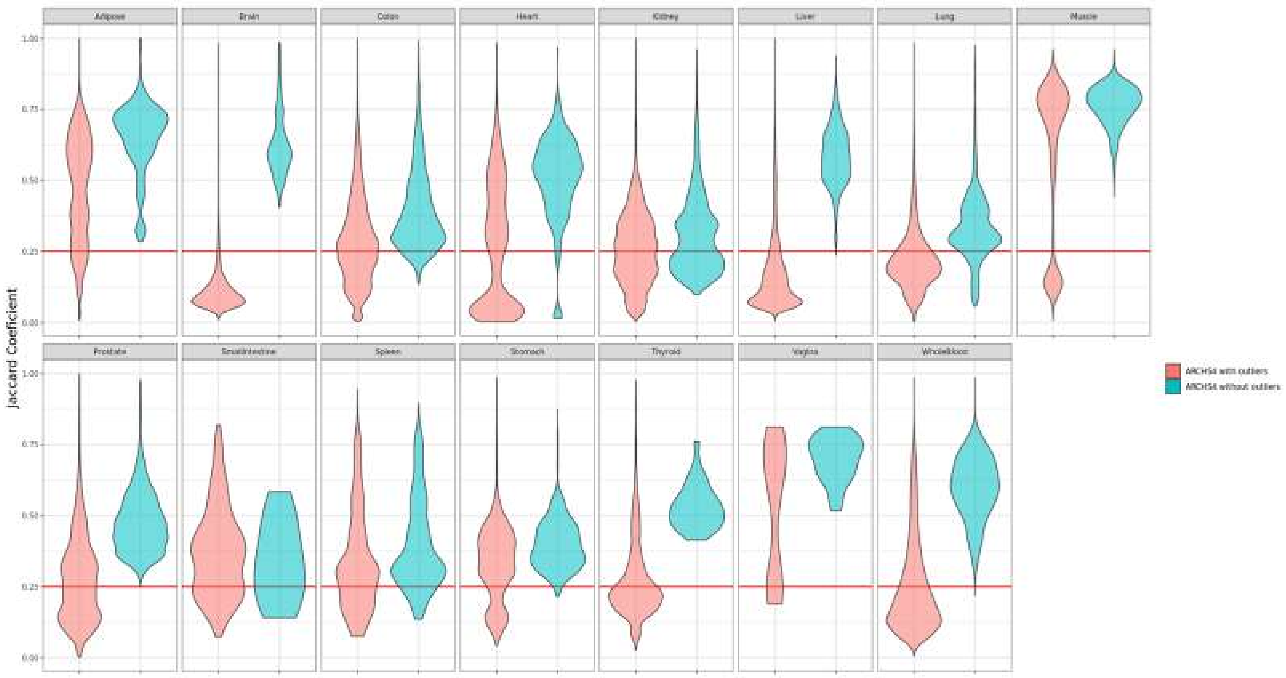


**Fig. S16:** Internal ARCHS4 GES tissue consistency before and after outlier removal.

### SUPPLEMENTAL TABLES

**Table S1:** Number of experimental samples per GTeX tissue type

| Tissue | Number of samples |
|---|---|
| **Adipose-Subcutaneous** | 442 |
| **Artery-Aorta** | 299 |
| **Artery-Coronary** | 173 |
| **Artery-Tibial** | 441 |
| **Brain-Amygdala** | 100 |
| **Brain-Cerebellar Hemisphere** | 136 |
| **Brain-Cerebellum** | 173 |
| **Brain-Cortex** | 158 |
| **Brain-Hippocampus** | 123 |
| **Brain-Hypothalamus** | 121 |
| **Brain-Substantianigra** | 88 |
| **Breast-Mammary Tissue** | 290 |
| **Cells-EBV-transformed lymphocytes** | 130 |
| **Cells-Transformed fibroblasts** | 343 |
| **Cervix-Ectocervix** | 6 |
| **Cervix-Endocervix** | 5 |
| **Colon-Sigmoid** | 233 |
| **Colon-Transverse** | 274 |
| **Esophagus-Gastroesophageal Junction** | 244 |
| **Esophagus-Mucosa** | 407 |
| **Esophagus-Muscularis** | 370 |
| **Fallopian Tube** | 7 |
| **Heart-Atrial Appendage** | 297 |
| **Heart-Left Ventricle** | 303 |
| **Kidney-Cortex** | 45 |
| **Liver** | 175 |
| **Lung** | 427 |
| **Minor Salivary Gland** | 97 |
| **Muscle-Skeletal** | 564 |
| **Nerve-Tibial** | 414 |
| **Ovary** | 133 |
| **Pancreas** | 248 |
| **Pituitary** | 183 |
| **Prostate** | 152 |
| **Small Intestine-Terminal Ileum** | 137 |
| **Spleen** | 162 |
| **Stomach** | 262 |
| **Testis** | 259 |
| **Thyroid** | 446 |
| **Uterus** | 111 |
| **Vagina** | 115 |
| **Whole Blood** | 406 |

**Table S2:** Significance testing of Jaccard Coefficient distributions between 50, 250, and 1000 gene sets. In bold are non-significant comparisons between gene set sizes.

| Sub-tissue | Tissue | Comparison | p-value | Adjusted p-value (Q-value) | Lower CI | Upper CI |
|---|---|---|---|---|---|---|
| Adipose-Subcutaneous | Adipose | 50 genes to 250 genes | 1.33E-303 | 3.98E-303 | -0.01386 | -0.01247 |
| Adipose-Subcutaneous | Adipose | 50 genes to 1000 genes | <2.2e-16 | <2.2e-16 | -0.04209 | -0.04077 |
| Adipose-Subcutaneous | Adipose | 250 genes to 1000 genes | <2.2e-16 | <2.2e-16 | -0.02881 | -0.02771 |
| Artery-Aorta | Artery | 50 genes to 250 genes | <2.2e-16 | <2.2e-16 | -0.03273 | -0.03073 |
| Artery-Aorta | Artery | 50 genes to 1000 genes | <2.2e-16 | <2.2e-16 | -0.05295 | -0.05104 |
| Artery-Aorta | Artery | 250 genes to 1000 genes | <2.2e-16 | <2.2e-16 | -0.02101 | -0.01952 |
| Artery-Coronary | Artery | 50 genes to 250 genes | 4.71E-215 | 1.41E-214 | -0.03606 | -0.03185 |
| Artery-Coronary | Artery | 50 genes to 1000 genes | <2.2e-16 | <2.2e-16 | -0.06327 | -0.05913 |
| Artery-Coronary | Artery | 250 genes to 1000 genes | 6.58E-216 | 1.97E-215 | -0.02893 | -0.02556 |
| Artery-Tibial | Artery | 50 genes to 250 genes | <2.2e-16 | <2.2e-16 | -0.02375 | -0.02237 |
| Artery-Tibial | Artery | 50 genes to 1000 genes | <2.2e-16 | <2.2e-16 | -0.06671 | -0.06534 |
| Artery-Tibial | Artery | 250 genes to 1000 genes | <2.2e-16 | <2.2e-16 | -0.04352 | -0.04241 |
| Brain-Amygdala | Brain | 50 genes to 250 genes | 3.40E-06 | 1.02E-05 | 0.005052 | 0.012423 |
| Brain-Amygdala | Brain | 50 genes to 1000 genes | 1.92E-50 | 5.77E-50 | -0.03013 | -0.02318 |
| Brain-Amygdala | Brain | 250 genes to 1000 genes | 3.35E-99 | 1.01E-98 | -0.03864 | -0.03215 |
| Brain-Cerebellar Hemisphere | Brain | 50 genes to 250 genes | 1.18E-215 | 3.55E-215 | -0.0511 | -0.04517 |
| Brain-Cerebellar Hemisphere | Brain | 50 genes to 1000 genes | <2.2e-16 | <2.2e-16 | -0.08009 | -0.07431 |
| Brain-Cerebellar Hemisphere | Brain | 250 genes to 1000 genes | 3.19E-116 | 9.58E-116 | -0.03153 | -0.0266 |
| Brain-Cerebellum | Brain | 50 genes to 250 genes | <2.2e-16 | <2.2e-16 | -0.08564 | -0.08186 |
| Brain-Cerebellum | Brain | 50 genes to 1000 genes | <2.2e-16 | <2.2e-16 | -0.10155 | -0.09796 |
| Brain-Cerebellum | Brain | 250 genes to 1000 genes | 1.06E-83 | 3.18E-83 | -0.01761 | -0.01439 |
| Brain-Cortex | Brain | 50 genes to 250 genes | 1.71E-36 | 5.12E-36 | -0.01943 | -0.01421 |
| Brain-Cortex | Brain | 50 genes to 1000 genes | <2.2e-16 | <2.2e-16 | -0.0585 | -0.05372 |
| Brain-Cortex | Brain | 250 genes to 1000 genes | 1.99E-204 | 5.97E-204 | -0.04179 | -0.03679 |
| Brain-Hippocampus | Brain | 50 genes to 250 genes | 5.06E-45 | 1.52E-44 | 0.020352 | 0.02691 |
| Brain-Hippocampus | Brain | 50 genes to 1000 genes | 1.03E-77 | 3.08E-77 | -0.03271 | -0.02653 |
| Brain-Hippocampus | Brain | 250 genes to 1000 genes | 1.98E-241 | 5.94E-241 | -0.05634 | -0.05016 |
| Brain-Hypothalamus | Brain | 50 genes to 250 genes | 6.29E-43 | 1.89E-42 | -0.02589 | -0.01944 |
| Brain-Hypothalamus | Brain | 50 genes to 1000 genes | <2.2e-16 | <2.2e-16 | -0.0734 | -0.06724 |
| Brain-Hypothalamus | Brain | 250 genes to 1000 genes | 9.31E-201 | 2.79E-200 | -0.0507 | -0.04462 |
| Brain-Substantianigra | Brain | 50 genes to 250 genes | 1.24E-39 | 3.72E-39 | 0.028412 | 0.038278 |

| | | | | | | |
|---|---|---|---|---|---|---|
| Brain-Substantianigra | Brain | 50 genes to 1000 genes | 0.012944721 | 0.038834164 | 0.001242 | 0.010504 |
| Brain-Substantianigra | Brain | 250 genes to 1000 genes | 9.34E-34 | 2.80E-33 | -0.0319 | -0.02305 |
| Breast-MammaryTissue | Breast | 50 genes to 250 genes | 2.48E-31 | 7.43E-31 | -0.00831 | -0.00592 |
| Breast-Mammary Tissue | Breast | 50 genes to 1000 genes | 8.99E-216 | 2.70E-215 | -0.01984 | -0.01751 |
| Breast-Mammary Tissue | Breast | 250 genes to 1000 genes | 3.41E-108 | 1.02E-107 | -0.01258 | -0.01053 |
| Cells-EBV-transformed lymphocytes | Cells | 50 genes to 250 genes | 3.08E-149 | 9.23E-149 | 0.020401 | 0.023687 |
| Cells-EBV-transformed lymphocytes | Cells | 50 genes to 1000 genes | 4.49E-238 | 1.35E-237 | 0.026857 | 0.030193 |
| Cells-EBV-transformed lymphocytes | Cells | 250 genes to 1000 genes | 1.57E-18 | 4.70E-18 | 0.005037 | 0.007925 |
| Cells-Transformed fibroblasts | Cells | 50 genes to 250 genes | 2.86E-159 | 8.58E-159 | 0.015474 | 0.017903 |
| Cells-Transformed fibroblasts | Cells | 50 genes to 1000 genes | 1.16E-82 | 3.49E-82 | -0.01238 | -0.01009 |
| Cells-Transformed fibroblasts | Cells | 250 genes to 1000 genes | <2.2e-16 | <2.2e-16 | -0.02898 | -0.02687 |
| Cervix-Ectocervix | Cervix | 50 genes to 250 genes | 0.187487245 | 0.562461734 | -0.21536 | 0.044177 |
| **Cervix-Ectocervix** | **Cervix** | **50 genes to 1000 genes** | **0.040312858** | **0.120938573** | **-0.24655** | **-0.006** |
| **Cervix-Ectocervix** | **Cervix** | **250 genes to 1000 genes** | **0.489284816** | **1** | **-0.15973** | **0.078362** |
| **Cervix-Endocervix** | **Cervix** | **50 genes to 250 genes** | **0.032633101** | **0.097899302** | **-0.17638** | **-0.00897** |
| **Cervix-Endocervix** | **Cervix** | **50 genes to 1000 genes** | **0.026960418** | **0.080881255** | **-0.17958** | **-0.01293** |
| **Cervix-Endocervix** | **Cervix** | **250 genes to 1000 genes** | **0.875118825** | **1** | **-0.05071** | **0.043557** |
| Colon-Sigmoid | Colon | 50 genes to 250 genes | 1.45E-183 | 4.34E-183 | 0.020263 | 0.023199 |
| Colon-Sigmoid | Colon | 50 genes to 1000 genes | <2.2e-16 | <2.2e-16 | 0.031268 | 0.034117 |
| Colon-Sigmoid | Colon | 250 genes to 1000 genes | 1.02E-74 | 3.06E-74 | 0.009788 | 0.012134 |
| Colon-Transverse | Colon | 50 genes to 250 genes | 7.64E-97 | 2.29E-96 | 0.019027 | 0.022962 |
| Colon-Transverse | Colon | 50 genes to 1000 genes | 2.69E-16 | 8.08E-16 | -0.01021 | -0.00626 |
| Colon-Transverse | Colon | 250 genes to 1000 genes | 2.29E-190 | 6.88E-190 | -0.03117 | -0.02729 |
| Esophagus-Gastroesophageal Junction | Esophagus | 50 genes to 250 genes | 2.89E-19 | 8.66E-19 | 0.004716 | 0.007351 |
| Esophagus-Gastroesophageal Junction | Esophagus | 50 genes to 1000 genes | 2.30E-17 | 6.91E-17 | 0.004356 | 0.006975 |
| **Esophagus-Gastroesophageal Junction** | **Esophagus** | **250 genes to 1000 genes** | **0.509019995** | **1** | **-0.00146** | **0.000724** |
| Esophagus-Mucosa | Esophagus | 50 genes to 250 genes | <2.2e-16 | <2.2e-16 | -0.02002 | -0.01826 |
| Esophagus-Mucosa | Esophagus | 50 genes to 1000 genes | 5.60447655845849e-313 | 1.68134296753755e-312 | -0.01721 | -0.01552 |
| Esophagus-Mucosa | Esophagus | 250 genes to 1000 genes | 5.63E-14 | 1.69E-13 | 0.002049 | 0.003494 |
| Esophagus-Muscularis | Esophagus | 50 genes to 250 genes | 1.37E-244 | 4.12E-244 | 0.013444 | 0.015116 |
| Esophagus-Muscularis | Esophagus | 50 genes to 1000 genes | 1.20E-240 | 3.60E-240 | 0.013351 | 0.015027 |
| **Esophagus-Muscularis** | **Esophagus** | **250 genes to 1000 genes** | **0.806306142** | **1** | **-0.00082** | **0.000636** |
| **Fallopian Tube** | **Fallopian Tube** | **50 genes to 250 genes** | **0.07081138** | **0.212434139** | **-0.11556** | **0.004919** |
| **Fallopian Tube** | **Fallopian Tube** | **50 genes to 1000 genes** | **0.115318163** | **0.345954489** | **-0.11275** | **0.012793** |
| **Fallopian Tube** | **Fallopian Tube** | **250 genes to 1000 genes** | **0.871501977** | **1** | **-0.06095** | **0.071632** |

| | | | | | | |
|---|---|---|---|---|---|---|
| Heart-Atrial Appendage | Heart | 50 genes to 250 genes | <2.2e-16 | <2.2e-16 | 0.031005 | 0.033443 |
| Heart-Atrial Appendage | Heart | 50 genes to 1000 genes | 1.00E-19 | 3.01E-19 | 0.004189 | 0.006492 |
| Heart-Atrial Appendage | Heart | 250 genes to 1000 genes | <2.2e-16 | <2.2e-16 | -0.02785 | -0.02592 |
| Heart-Left Ventricle | Heart | 50 genes to 250 genes | <2.2e-16 | <2.2e-16 | 0.043384 | 0.046803 |
| Heart-Left Ventricle | Heart | 50 genes to 1000 genes | 4.55E-72 | 1.36E-71 | 0.013376 | 0.016651 |
| Heart-Left Ventricle | Heart | 250 genes to 1000 genes | 6.030209545873 25e-318 | 1.80906286376197e-317 | -0.03162 | -0.02854 |
| Kidney-Cortex | Kidney | 50 genes to 250 genes | 2.52E-09 | 7.56E-09 | 0.014431 | 0.028488 |
| **Kidney-Cortex** | **Kidney** | **50 genes to 1000 genes** | **0.770840118** | **1** | **-0.00588** | **0.007934** |
| Kidney-Cortex | Kidney | 250 genes to 1000 genes | 8.80E-09 | 2.64E-08 | -0.02737 | -0.0135 |
| Liver | Liver | 50 genes to 250 genes | <2.2e-16 | <2.2e-16 | 0.0837 | 0.088044 |
| Liver | Liver | 50 genes to 1000 genes | 1.12E-201 | 3.35E-201 | 0.031125 | 0.035395 |
| Liver | Liver | 250 genes to 1000 genes | <2.2e-16 | <2.2e-16 | -0.0546 | -0.05062 |
| Lung | Lung | 50 genes to 250 genes | <2.2e-16 | <2.2e-16 | 0.016768 | 0.018231 |
| Lung | Lung | 50 genes to 1000 genes | 1.61E-58 | 4.84E-58 | -0.00673 | -0.00527 |
| Lung | Lung | 250 genes to 1000 genes | <2.2e-16 | <2.2e-16 | -0.02407 | -0.02292 |
| Minor Salivary Gland | Minor Salivary Gland | 50 genes to 250 genes | 1.42E-21 | 4.27E-21 | -0.034 | -0.02243 |
| Minor Salivary Gland | Minor Salivary Gland | 50 genes to 1000 genes | 3.98E-136 | 1.19E-135 | -0.07733 | -0.06621 |
| Minor Salivary Gland | Minor Salivary Gland | 250 genes to 1000 genes | 1.81E-73 | 5.43E-73 | -0.04823 | -0.03889 |
| Muscle-Skeletal | Muscle | 50 genes to 250 genes | <2.2e-16 | <2.2e-16 | 0.044315 | 0.045661 |
| Muscle-Skeletal | Muscle | 50 genes to 1000 genes | 1.93E-32 | 5.80E-32 | 0.003211 | 0.004482 |
| Muscle-Skeletal | Muscle | 250 genes to 1000 genes | <2.2e-16 | <2.2e-16 | -0.04166 | -0.04062 |
| Nerve-Tibial | Nerve | 50 genes to 250 genes | <2.2e-16 | <2.2e-16 | 0.025431 | 0.02683 |
| Nerve-Tibial | Nerve | 50 genes to 1000 genes | <2.2e-16 | <2.2e-16 | 0.026524 | 0.027925 |
| Nerve-Tibial | Nerve | 250 genes to 1000 genes | 0.000215816 | 0.000647447 | 0.000515 | 0.001674 |
| Ovary | Ovary | 50 genes to 250 genes | 1.98E-119 | 5.94E-119 | 0.034086 | 0.040299 |
| Ovary | Ovary | 50 genes to 1000 genes | 8.15E-229 | 2.44E-228 | 0.048333 | 0.054445 |
| Ovary | Ovary | 250 genes to 1000 genes | 1.79E-43 | 5.38E-43 | 0.012189 | 0.016203 |
| Pancreas | Pancreas | 50 genes to 250 genes | <2.2e-16 | <2.2e-16 | 0.052842 | 0.055092 |
| Pancreas | Pancreas | 50 genes to 1000 genes | <2.2e-16 | <2.2e-16 | 0.072408 | 0.07463 |
| Pancreas | Pancreas | 250 genes to 1000 genes | <2.2e-16 | <2.2e-16 | 0.018589 | 0.020515 |
| Pituitary | Pituitary | 50 genes to 250 genes | 2.23E-44 | 6.70E-44 | -0.01805 | -0.01362 |
| Pituitary | Pituitary | 50 genes to 1000 genes | 2.79E-13 | 8.38E-13 | -0.01038 | -0.00599 |
| Pituitary | Pituitary | 250 genes to 1000 genes | 4.72E-15 | 1.42E-14 | 0.005734 | 0.009558 |
| Prostate | Prostate | 50 genes to 250 genes | 1.27E-14 | 3.81E-14 | 0.007105 | 0.011946 |
| Prostate | Prostate | 50 genes to 1000 genes | 5.61E-52 | 1.68E-51 | 0.01616 | 0.020941 |

| | | | | | | |
|---|---|---|---|---|---|---|
| Prostate | Prostate | 250 genes to 1000 genes | 8.20E-18 | 2.46E-17 | 0.006969 | 0.011081 |
| Small Intestine-Terminal Ileum | Small Intestine | 50 genes to 250 genes | 3.67E-48 | 1.10E-47 | 0.026032 | 0.03409 |
| **Small Intestine-Terminal Ileum** | **Small Intestine** | **50 genes to 1000 genes** | **0.646645602** | **1** | **-0.00494** | **0.003065** |
| Small Intestine-Terminal Ileum | Small Intestine | 250 genes to 1000 genes | 1.68E-57 | 5.05E-57 | -0.03479 | -0.02721 |
| Spleen | Spleen | 50 genes to 250 genes | 3.27E-12 | 9.80E-12 | -0.01163 | -0.00652 |
| Spleen | Spleen | 50 genes to 1000 genes | 2.13E-284 | 6.40E-284 | -0.04739 | -0.04258 |
| Spleen | Spleen | 250 genes to 1000 genes | 7.905050333459 94e-323 | 2.37151510003798e-322 | -0.03771 | -0.0341 |
| Stomach | Stomach | 50 genes to 250 genes | 0.03121529 | 0.093645871 | -0.00417 | -0.0002 |
| Stomach | Stomach | 50 genes to 1000 genes | 2.43E-50 | 7.29E-50 | -0.01659 | -0.01274 |
| Stomach | Stomach | 250 genes to 1000 genes | 3.04E-38 | 9.12E-38 | -0.01438 | -0.01059 |
| Testis | Testis | 50 genes to 250 genes | 2.04E-207 | 6.13E-207 | 0.034003 | 0.038618 |
| Testis | Testis | 50 genes to 1000 genes | 2.40E-37 | 7.21E-37 | 0.012091 | 0.016472 |
| Testis | Testis | 250 genes to 1000 genes | 5.15E-82 | 1.54E-81 | -0.02428 | -0.01978 |
| Thyroid | Thyroid | 50 genes to 250 genes | <2.2e-16 | <2.2e-16 | -0.05261 | -0.0512 |
| Thyroid | Thyroid | 50 genes to 1000 genes | <2.2e-16 | <2.2e-16 | -0.05628 | -0.05492 |
| Thyroid | Thyroid | 250 genes to 1000 genes | 9.77E-35 | 2.93E-34 | -0.00428 | -0.00311 |
| Uterus | Uterus | 50 genes to 250 genes | 3.83E-108 | 1.15E-107 | -0.04233 | -0.03551 |
| Uterus | Uterus | 50 genes to 1000 genes | 3.90E-40 | 1.17E-39 | -0.02645 | -0.01967 |
| Uterus | Uterus | 250 genes to 1000 genes | 2.03E-36 | 6.09E-36 | 0.013401 | 0.018318 |
| Vagina | Vagina | 50 genes to 250 genes | 1.33E-243 | 3.99E-243 | -0.08224 | -0.07329 |
| Vagina | Vagina | 50 genes to 1000 genes | <2.2e-16 | <2.2e-16 | -0.10188 | -0.09346 |
| Vagina | Vagina | 250 genes to 1000 genes | 1.26E-24 | 3.79E-24 | -0.0237 | -0.0161 |
| Whole Blood | Whole Blood | 50 genes to 250 genes | 1.84E-08 | 5.53E-08 | -0.00434 | -0.0021 |
| Whole Blood | Whole Blood | 50 genes to 1000 genes | <2.2e-16 | <2.2e-16 | -0.0814 | -0.07937 |
| Whole Blood | Whole Blood | 250 genes to 1000 genes | <2.2e-16 | <2.2e-16 | -0.07818 | -0.07615 |

**Table S3:** Confusion matrix of validation set of GTeX model against ARCHS4 for gene signature size of 50.

| | Predicted Labels | | | | | | | | | | | | | | | | | |
|---|---|---|---|---|---|---|---|---|---|---|---|---|---|---|---|---|---|---|
| Observed Labels | Adipose | Artery | Brain | Cells | Colon | Esophagus | Heart | Liver | Lung | Muscle | Nerve | Ovary | Pancreas | SmallIntestine | Stomach | Testis | Thyroid | WholeBlood |
| Adipose | 79 | 11 | 1 | 5 | 0 | 0 | 0 | 0 | 0 | 0 | 0 | 0 | 0 | 0 | 0 | 0 | 0 | 4 |
| Brain | 0 | 9 | 57 | 1 | 0 | 1 | 0 | 0 | 0 | 0 | 0 | 1 | 0 | 0 | 0 | 0 | 0 | 31 |
| Colon | 0 | 7 | 14 | 10 | 43 | 10 | 0 | 0 | 0 | 0 | 0 | 0 | 0 | 1 | 0 | 0 | 0 | 15 |
| Esophagus | 0 | 27 | 28 | 11 | 0 | 27 | 0 | 0 | 0 | 0 | 0 | 0 | 0 | 0 | 0 | 0 | 0 | 6 |
| FallopianTube | 0 | 2 | 2 | 1 | 0 | 0 | 0 | 0 | 0 | 0 | 0 | 0 | 0 | 0 | 0 | 0 | 0 | 1 |
| Heart | 0 | 4 | 6 | 0 | 0 | 2 | 72 | 0 | 0 | 2 | 0 | 0 | 0 | 0 | 0 | 0 | 0 | 14 |
| Kidney | 0 | 10 | 43 | 17 | 0 | 5 | 0 | 0 | 0 | 0 | 0 | 22 | 0 | 0 | 0 | 0 | 0 | 3 |
| Liver | 0 | 11 | 11 | 10 | 0 | 0 | 0 | 50 | 0 | 0 | 0 | 0 | 0 | 0 | 0 | 0 | 0 | 18 |
| Lung | 0 | 5 | 39 | 19 | 0 | 6 | 0 | 0 | 19 | 0 | 0 | 0 | 0 | 0 | 0 | 0 | 0 | 12 |
| Muscle | 0 | 0 | 4 | 9 | 0 | 0 | 0 | 0 | 0 | 87 | 0 | 0 | 0 | 0 | 0 | 0 | 0 | 0 |
| Nerve | 0 | 11 | 3 | 0 | 0 | 0 | 0 | 0 | 0 | 0 | 1 | 0 | 0 | 0 | 0 | 0 | 0 | 0 |
| Ovary | 0 | 7 | 10 | 2 | 0 | 1 | 0 | 0 | 0 | 0 | 0 | 1 | 0 | 0 | 0 | 0 | 0 | 0 |
| Pancreas | 0 | 6 | 75 | 1 | 0 | 2 | 0 | 0 | 0 | 0 | 0 | 0 | 7 | 0 | 0 | 0 | 0 | 9 |
| Prostate | 0 | 13 | 41 | 17 | 0 | 22 | 0 | 0 | 0 | 0 | 0 | 0 | 0 | 0 | 0 | 0 | 0 | 6 |
| SmallIntestine | 0 | 17 | 3 | 1 | 1 | 1 | 0 | 0 | 0 | 0 | 0 | 0 | 0 | 1 | 0 | 0 | 0 | 1 |
| Spleen | 0 | 2 | 6 | 4 | 0 | 0 | 0 | 0 | 0 | 0 | 0 | 1 | 0 | 0 | 0 | 0 | 0 | 49 |
| Stomach | 0 | 52 | 13 | 12 | 0 | 12 | 0 | 0 | 0 | 0 | 0 | 1 | 0 | 0 | 1 | 0 | 0 | 9 |
| Testis | 0 | 0 | 13 | 1 | 0 | 1 | 0 | 0 | 0 | 0 | 0 | 0 | 0 | 0 | 0 | 6 | 0 | 3 |
| Thyroid | 0 | 34 | 18 | 6 | 0 | 0 | 0 | 0 | 0 | 2 | 0 | 0 | 0 | 0 | 0 | 0 | 35 | 4 |
| Uterus | 0 | 5 | 4 | 0 | 0 | 0 | 0 | 0 | 0 | 0 | 0 | 0 | 0 | 0 | 0 | 0 | 0 | 0 |
| Vagina | 0 | 0 | 0 | 0 | 0 | 10 | 0 | 0 | 0 | 0 | 0 | 0 | 0 | 0 | 0 | 0 | 0 | 0 |
| WholeBlood | 0 | 1 | 10 | 4 | 0 | 0 | 0 | 0 | 0 | 0 | 0 | 1 | 0 | 0 | 0 | 0 | 0 | 84 |

**Table S4:** Confusion matrix of validation set of GTeX model against ARCHS4 for gene signature size of 250.

| | Predicted Labels | | | | | | | | | | | | | | | | | |
|---|---|---|---|---|---|---|---|---|---|---|---|---|---|---|---|---|---|---|
| Observed Labels | Adipose | Artery | Brain | Cells | Colon | Esophagus | Heart | Liver | Lung | Muscle | Nerve | Pancreas | SmallIntestine | Spleen | Stomach | Testis | Thyroid | WholeBlood |
| Adipose | 83 | 2 | 2 | 12 | 0 | 0 | 0 | 0 | 0 | 0 | 0 | 0 | 0 | 0 | 0 | 0 | 0 | 1 |
| Brain | 0 | 1 | 65 | 4 | 0 | 0 | 0 | 0 | 0 | 0 | 0 | 0 | 0 | 0 | 0 | 0 | 0 | 30 |
| Colon | 0 | 2 | 3 | 34 | 40 | 5 | 0 | 0 | 0 | 0 | 0 | 0 | 3 | 0 | 0 | 0 | 0 | 13 |
| Esophagus | 0 | 3 | 25 | 28 | 0 | 41 | 0 | 0 | 0 | 0 | 0 | 0 | 0 | 0 | 0 | 0 | 0 | 2 |
| FallopianTube | 0 | 0 | 2 | 3 | 0 | 1 | 0 | 0 | 0 | 0 | 0 | 0 | 0 | 0 | 0 | 0 | 0 | 0 |
| Heart | 0 | 0 | 4 | 0 | 0 | 0 | 76 | 0 | 0 | 2 | 0 | 0 | 0 | 0 | 0 | 0 | 0 | 18 |
| Kidney | 0 | 3 | 30 | 66 | 0 | 1 | 0 | 0 | 0 | 0 | 0 | 0 | 0 | 0 | 0 | 0 | 0 | 0 |
| Liver | 0 | 0 | 8 | 23 | 0 | 5 | 0 | 50 | 0 | 0 | 0 | 0 | 0 | 0 | 0 | 0 | 0 | 14 |
| Lung | 0 | 0 | 6 | 47 | 0 | 15 | 0 | 0 | 13 | 0 | 0 | 0 | 0 | 0 | 0 | 0 | 0 | 19 |
| Muscle | 0 | 0 | 1 | 11 | 0 | 0 | 0 | 0 | 0 | 88 | 0 | 0 | 0 | 0 | 0 | 0 | 0 | 0 |
| Nerve | 0 | 2 | 5 | 6 | 0 | 0 | 0 | 0 | 0 | 0 | 2 | 0 | 0 | 0 | 0 | 0 | 0 | 0 |
| Ovary | 0 | 4 | 5 | 8 | 0 | 1 | 0 | 0 | 0 | 0 | 0 | 0 | 0 | 0 | 0 | 0 | 0 | 3 |
| Pancreas | 0 | 0 | 73 | 12 | 0 | 5 | 0 | 0 | 0 | 0 | 0 | 4 | 0 | 0 | 0 | 0 | 0 | 6 |
| Prostate | 0 | 10 | 31 | 43 | 0 | 11 | 0 | 0 | 0 | 0 | 0 | 0 | 0 | 0 | 0 | 0 | 0 | 4 |
| SmallIntestine | 0 | 5 | 0 | 9 | 0 | 7 | 0 | 0 | 0 | 0 | 0 | 0 | 3 | 0 | 0 | 0 | 0 | 1 |
| Spleen | 0 | 0 | 0 | 13 | 0 | 2 | 0 | 0 | 0 | 0 | 0 | 0 | 0 | 2 | 0 | 0 | 0 | 45 |
| Stomach | 0 | 4 | 14 | 67 | 0 | 13 | 0 | 0 | 0 | 0 | 0 | 0 | 0 | 0 | 1 | 0 | 0 | 1 |
| Testis | 0 | 0 | 6 | 6 | 0 | 0 | 0 | 0 | 0 | 0 | 0 | 0 | 0 | 0 | 0 | 9 | 0 | 3 |
| Thyroid | 0 | 2 | 22 | 21 | 0 | 1 | 0 | 0 | 0 | 5 | 0 | 0 | 0 | 0 | 0 | 0 | 47 | 1 |
| Uterus | 0 | 5 | 2 | 1 | 0 | 1 | 0 | 0 | 0 | 0 | 0 | 0 | 0 | 0 | 0 | 0 | 0 | 0 |
| Vagina | 0 | 0 | 0 | 0 | 0 | 10 | 0 | 0 | 0 | 0 | 0 | 0 | 0 | 0 | 0 | 0 | 0 | 0 |
| WholeBlood | 1 | 0 | 6 | 10 | 0 | 1 | 0 | 0 | 0 | 0 | 0 | 0 | 0 | 0 | 0 | 0 | 0 | 82 |

**Table S5:** Confusion matrix of validation set of GTeX model against ARCHS4 for gene signature size of 1,000.

| | Predicted Labels | | | | | | | | | | | | | | | | | | | |
|---|---|---|---|---|---|---|---|---|---|---|---|---|---|---|---|---|---|---|---|---|
| Observed Labels | Adipose | Artery | Brain | Cells | Colon | Esophagus | Heart | Kidney | Liver | Lung | Muscle | Nerve | Ovary | Pancreas | SmallIntestine | Spleen | Stomach | Testis | Thyroid | WholeBlood |
| Adipose | 85 | 0 | 1 | 13 | 0 | 0 | 0 | 0 | 0 | 0 | 0 | 0 | 0 | 0 | 0 | 0 | 0 | 0 | 0 | 1 |
| Brain | 0 | 0 | 66 | 7 | 0 | 0 | 0 | 0 | 0 | 0 | 0 | 0 | 0 | 0 | 0 | 0 | 0 | 0 | 0 | 27 |
| Colon | 0 | 0 | 3 | 36 | 48 | 4 | 0 | 0 | 0 | 2 | 0 | 0 | 0 | 0 | 2 | 0 | 0 | 0 | 0 | 5 |
| Esophagus | 0 | 0 | 40 | 35 | 0 | 23 | 0 | 0 | 0 | 0 | 0 | 0 | 0 | 0 | 0 | 0 | 0 | 0 | 0 | 1 |
| FallopianTube | 0 | 0 | 0 | 6 | 0 | 0 | 0 | 0 | 0 | 0 | 0 | 0 | 0 | 0 | 0 | 0 | 0 | 0 | 0 | 0 |
| Heart | 0 | 0 | 17 | 0 | 0 | 0 | 76 | 0 | 0 | 0 | 2 | 0 | 0 | 0 | 0 | 0 | 0 | 0 | 0 | 5 |
| Kidney | 0 | 0 | 7 | 82 | 0 | 0 | 0 | 10 | 1 | 0 | 0 | 0 | 0 | 0 | 0 | 0 | 0 | 0 | 0 | 0 |
| Liver | 0 | 0 | 0 | 49 | 0 | 1 | 0 | 0 | 45 | 1 | 0 | 0 | 0 | 0 | 0 | 0 | 0 | 0 | 0 | 4 |
| Lung | 0 | 0 | 4 | 74 | 0 | 0 | 0 | 0 | 0 | 14 | 0 | 0 | 0 | 0 | 0 | 0 | 0 | 0 | 0 | 8 |
| Muscle | 0 | 0 | 0 | 11 | 0 | 0 | 0 | 0 | 0 | 0 | 89 | 0 | 0 | 0 | 0 | 0 | 0 | 0 | 0 | 0 |
| Nerve | 0 | 0 | 3 | 6 | 0 | 0 | 0 | 0 | 0 | 0 | 0 | 6 | 0 | 0 | 0 | 0 | 0 | 0 | 0 | 0 |
| Ovary | 0 | 2 | 5 | 11 | 0 | 0 | 0 | 0 | 0 | 0 | 0 | 0 | 3 | 0 | 0 | 0 | 0 | 0 | 0 | 0 |
| Pancreas | 0 | 0 | 64 | 30 | 1 | 0 | 0 | 0 | 0 | 0 | 0 | 0 | 0 | 4 | 0 | 0 | 0 | 0 | 0 | 1 |
| Prostate | 0 | 5 | 7 | 68 | 0 | 18 | 0 | 0 | 0 | 1 | 0 | 0 | 0 | 0 | 0 | 0 | 0 | 0 | 0 | 0 |
| SmallIntestine | 0 | 1 | 0 | 16 | 5 | 0 | 0 | 0 | 0 | 0 | 0 | 0 | 0 | 0 | 3 | 0 | 0 | 0 | 0 | 0 |
| Spleen | 0 | 0 | 1 | 22 | 0 | 0 | 0 | 0 | 0 | 0 | 0 | 0 | 0 | 0 | 0 | 5 | 0 | 0 | 0 | 34 |
| Stomach | 0 | 0 | 27 | 65 | 2 | 2 | 2 | 0 | 0 | 0 | 0 | 0 | 0 | 0 | 0 | 0 | 2 | 0 | 0 | 0 |
| Testis | 0 | 0 | 5 | 5 | 0 | 0 | 0 | 0 | 0 | 0 | 0 | 0 | 0 | 0 | 0 | 0 | 0 | 14 | 0 | 0 |
| Thyroid | 0 | 0 | 7 | 33 | 0 | 0 | 0 | 0 | 0 | 1 | 5 | 0 | 0 | 0 | 0 | 0 | 0 | 0 | 53 | 0 |
| Uterus | 0 | 3 | 0 | 6 | 0 | 0 | 0 | 0 | 0 | 0 | 0 | 0 | 0 | 0 | 0 | 0 | 0 | 0 | 0 | 0 |
| Vagina | 0 | 0 | 0 | 0 | 0 | 10 | 0 | 0 | 0 | 0 | 0 | 0 | 0 | 0 | 0 | 0 | 0 | 0 | 0 | 0 |
| WholeBlood | 0 | 0 | 5 | 22 | 0 | 0 | 0 | 0 | 0 | 1 | 0 | 0 | 0 | 0 | 0 | 0 | 0 | 0 | 0 | 72 |

**Table S6:** Significance testing of Jaccard Coefficient ($J_C$) distributions between top 250 genes, domain enrichment, family enrichment, superfamily enrichment and fold enrichment. In bold are non-significant comparisons.

| *Sub-tissue* | *Tissue* | *Comparison* | *p-value* | *Adjusted p-value (Q-value)* | *Lower CI* | *Upper CI* |
|---|---|---|---|---|---|---|
| *Adipose-Subcutaneous* | Adipose | 250 genes to domain enrichment | <2.2e-16 | <2.2e-16 | -0.028807034 | -0.027639507 |
| *Adipose-Subcutaneous* | Adipose | 250 genes to family enrichment | <2.2e-16 | <2.2e-16 | -0.021555627 | -0.019817127 |
| *Adipose-Subcutaneous* | Adipose | 250 genes to superfamily enrichment | <2.2e-16 | <2.2e-16 | -0.054296326 | -0.053148901 |
| *Adipose-Subcutaneous* | Adipose | 250 genes to fold enrichment | <2.2e-16 | <2.2e-16 | -0.075513278 | -0.074382919 |
| *Artery-Aorta* | Artery | 250 genes to domain enrichment | <2.2e-16 | <2.2e-16 | -0.0468739 | -0.045978865 |
| *Artery-Coronary* | Artery | 250 genes to family enrichment | <2.2e-16 | <2.2e-16 | -0.058802219 | -0.057869389 |
| *Artery-Tibial* | Artery | 250 genes to superfamily enrichment | <2.2e-16 | <2.2e-16 | -0.075265138 | -0.074374203 |
| *Artery-Aorta* | Artery | 250 genes to fold enrichment | <2.2e-16 | <2.2e-16 | -0.094817887 | -0.093938713 |
| *Artery-Coronary* | Artery | 250 genes to domain enrichment | <2.2e-16 | <2.2e-16 | -0.0468739 | -0.045978865 |
| *Artery-Tibial* | Artery | 250 genes to family enrichment | <2.2e-16 | <2.2e-16 | -0.058802219 | -0.057869389 |
| *Artery-Aorta* | Artery | 250 genes to superfamily enrichment | <2.2e-16 | <2.2e-16 | -0.075265138 | -0.074374203 |
| *Artery-Coronary* | Artery | 250 genes to fold enrichment | <2.2e-16 | <2.2e-16 | -0.094817887 | -0.093938713 |
| *Artery-Tibial* | Artery | 250 genes to domain enrichment | <2.2e-16 | <2.2e-16 | -0.0468739 | -0.045978865 |
| *Artery-Aorta* | Artery | 250 genes to family enrichment | <2.2e-16 | <2.2e-16 | -0.058802219 | -0.057869389 |
| *Artery-Coronary* | Artery | 250 genes to superfamily enrichment | <2.2e-16 | <2.2e-16 | -0.075265138 | -0.074374203 |
| *Artery-Tibial* | Artery | 250 genes to fold enrichment | <2.2e-16 | <2.2e-16 | -0.094817887 | -0.093938713 |

| | | | | | | |
|---|---|---|---|---|---|---|
| *Brain-Amygdala* | Brain | 250 genes to domain enrichment | <2.2e-16 | <2.2e-16 | -0.039561682 | -0.037105672 |
| *Brain-CerebellarHemisphere* | Brain | 250 genes to family enrichment | <2.2e-16 | <2.2e-16 | -0.046056815 | -0.042740733 |
| *Brain-Cerebellum* | Brain | 250 genes to superfamily enrichment | <2.2e-16 | <2.2e-16 | -0.089830889 | -0.087479828 |
| *Brain-Cortex* | Brain | 250 genes to fold enrichment | <2.2e-16 | <2.2e-16 | -0.112712792 | -0.11043837 |
| *Brain-Hippocampus* | Brain | 250 genes to domain enrichment | <2.2e-16 | <2.2e-16 | -0.039561682 | -0.037105672 |
| *Brain-Hypothalamus* | Brain | 250 genes to family enrichment | <2.2e-16 | <2.2e-16 | -0.046056815 | -0.042740733 |
| *Brain-Substantianigra* | Brain | 250 genes to superfamily enrichment | <2.2e-16 | <2.2e-16 | -0.089830889 | -0.087479828 |
| *Brain-Amygdala* | Brain | 250 genes to fold enrichment | <2.2e-16 | <2.2e-16 | -0.112712792 | -0.11043837 |
| *Brain-CerebellarHemisphere* | Brain | 250 genes to domain enrichment | <2.2e-16 | <2.2e-16 | -0.039561682 | -0.037105672 |
| *Brain-Cerebellum* | Brain | 250 genes to family enrichment | <2.2e-16 | <2.2e-16 | -0.046056815 | -0.042740733 |
| *Brain-Cortex* | Brain | 250 genes to superfamily enrichment | <2.2e-16 | <2.2e-16 | -0.089830889 | -0.087479828 |
| *Brain-Hippocampus* | Brain | 250 genes to fold enrichment | <2.2e-16 | <2.2e-16 | -0.112712792 | -0.11043837 |
| *Brain-Hypothalamus* | Brain | 250 genes to domain enrichment | <2.2e-16 | <2.2e-16 | -0.039561682 | -0.037105672 |
| *Brain-Substantianigra* | Brain | 250 genes to family enrichment | <2.2e-16 | <2.2e-16 | -0.046056815 | -0.042740733 |
| *Brain-Amygdala* | Brain | 250 genes to superfamily enrichment | <2.2e-16 | <2.2e-16 | -0.089830889 | -0.087479828 |
| *Brain-CerebellarHemisphere* | Brain | 250 genes to fold enrichment | <2.2e-16 | <2.2e-16 | -0.112712792 | -0.11043837 |
| *Brain-Cerebellum* | Brain | 250 genes to domain enrichment | <2.2e-16 | <2.2e-16 | -0.039561682 | -0.037105672 |
| *Brain-Cortex* | Brain | 250 genes to family enrichment | <2.2e-16 | <2.2e-16 | -0.046056815 | -0.042740733 |
| *Brain-Hippocampus* | Brain | 250 genes to superfamily enrichment | <2.2e-16 | <2.2e-16 | -0.089830889 | -0.087479828 |
| *Brain-Hypothalamus* | Brain | 250 genes to fold enrichment | <2.2e-16 | <2.2e-16 | -0.112712792 | -0.11043837 |
| *Brain-Substantianigra* | Brain | 250 genes to domain enrichment | <2.2e-16 | <2.2e-16 | -0.039561682 | -0.037105672 |
| *Brain-Amygdala* | Brain | 250 genes to family enrichment | <2.2e-16 | <2.2e-16 | -0.046056815 | -0.042740733 |
| *Brain-CerebellarHemisphere* | Brain | 250 genes to superfamily enrichment | <2.2e-16 | <2.2e-16 | -0.089830889 | -0.087479828 |
| *Brain-Cerebellum* | Brain | 250 genes to fold enrichment | <2.2e-16 | <2.2e-16 | -0.112712792 | -0.11043837 |
| *Brain-Cortex* | Brain | 250 genes to domain enrichment | <2.2e-16 | <2.2e-16 | -0.039561682 | -0.037105672 |
| *Brain-Hippocampus* | Brain | 250 genes to family enrichment | <2.2e-16 | <2.2e-16 | -0.046056815 | -0.042740733 |
| *Brain-Hypothalamus* | Brain | 250 genes to superfamily enrichment | <2.2e-16 | <2.2e-16 | -0.089830889 | -0.087479828 |
| *Brain-Substantianigra* | Brain | 250 genes to fold enrichment | <2.2e-16 | <2.2e-16 | -0.112712792 | -0.11043837 |
| *Breast-MammaryTissue* | Breast | 250 genes to domain enrichment | <2.2e-16 | <2.2e-16 | -0.043287928 | -0.041182562 |
| *Breast-MammaryTissue* | Breast | 250 genes to family enrichment | 9.88131291682493e-324 | 3.95252516672997e-323 | -0.031679649 | -0.028652187 |

| | | | | | | |
|---|---|---|---|---|---|---|
| *Breast-MammaryTissue* | Breast | 250 genes to superfamily enrichment | <2.2e-16 | <2.2e-16 | -0.074543065 | -0.072507838 |
| *Breast-MammaryTissue* | Breast | 250 genes to fold enrichment | <2.2e-16 | <2.2e-16 | -0.09289164 | -0.090884954 |
| *Cells-EBV-transformedlymphocytes* | Cells | 250 genes to domain enrichment | <2.2e-16 | <2.2e-16 | -0.068004623 | -0.065970003 |
| *Cells-Transformedfibroblasts* | Cells | 250 genes to family enrichment | <2.2e-16 | <2.2e-16 | -0.075121186 | -0.072368665 |
| *Cells-EBV-transformedlymphocytes* | Cells | 250 genes to superfamily enrichment | <2.2e-16 | <2.2e-16 | -0.109772313 | -0.107829144 |
| *Cells-Transformedfibroblasts* | Cells | 250 genes to fold enrichment | <2.2e-16 | <2.2e-16 | -0.134335325 | -0.132432907 |
| *Cells-EBV-transformedlymphocytes* | Cells | 250 genes to domain enrichment | <2.2e-16 | <2.2e-16 | -0.068004623 | -0.065970003 |
| *Cells-Transformedfibroblasts* | Cells | 250 genes to family enrichment | <2.2e-16 | <2.2e-16 | -0.075121186 | -0.072368665 |
| *Cells-EBV-transformedlymphocytes* | Cells | 250 genes to superfamily enrichment | <2.2e-16 | <2.2e-16 | -0.109772313 | -0.107829144 |
| *Cells-Transformedfibroblasts* | Cells | 250 genes to fold enrichment | <2.2e-16 | <2.2e-16 | -0.134335325 | -0.132432907 |
| *Cervix-Ectocervix* | **Cervix** | **250 genes to domain enrichment** | **0.159101699** | **0.636406797** | **-0.133603425** | **0.022574698** |
| *Cervix-Endocervix* | **Cervix** | **250 genes to family enrichment** | **0.721023413** | **1** | **-0.116122132** | **0.081521534** |
| *Cervix-Ectocervix* | Cervix | 250 genes to superfamily enrichment | 0.006226107 | 0.024904428 | -0.181081778 | -0.031886319 |
| *Cervix-Endocervix* | Cervix | 250 genes to fold enrichment | 0.000200823 | 0.000803293 | -0.215481434 | -0.073501941 |
| *Cervix-Ectocervix* | **Cervix** | **250 genes to domain enrichment** | **0.159101699** | **0.636406797** | **-0.133603425** | **0.022574698** |
| *Cervix-Endocervix* | **Cervix** | **250 genes to family enrichment** | **0.721023413** | **1** | **-0.116122132** | **0.081521534** |
| *Cervix-Ectocervix* | Cervix | 250 genes to superfamily enrichment | 0.006226107 | 0.024904428 | -0.181081778 | -0.031886319 |
| *Cervix-Endocervix* | Cervix | 250 genes to fold enrichment | 0.000200823 | 0.000803293 | -0.215481434 | -0.073501941 |
| *Colon-Sigmoid* | Colon | 250 genes to domain enrichment | <2.2e-16 | <2.2e-16 | -0.056157182 | -0.053485776 |
| *Colon-Transverse* | Colon | 250 genes to family enrichment | <2.2e-16 | <2.2e-16 | -0.043428614 | -0.039792655 |
| *Colon-Sigmoid* | Colon | 250 genes to superfamily enrichment | <2.2e-16 | <2.2e-16 | -0.089366141 | -0.086834843 |
| *Colon-Transverse* | Colon | 250 genes to fold enrichment | <2.2e-16 | <2.2e-16 | -0.100017995 | -0.09751364 |
| *Colon-Sigmoid* | Colon | 250 genes to domain enrichment | <2.2e-16 | <2.2e-16 | -0.056157182 | -0.053485776 |
| *Colon-Transverse* | Colon | 250 genes to family enrichment | <2.2e-16 | <2.2e-16 | -0.043428614 | -0.039792655 |
| *Colon-Sigmoid* | Colon | 250 genes to superfamily enrichment | <2.2e-16 | <2.2e-16 | -0.089366141 | -0.086834843 |
| *Colon-Transverse* | Colon | 250 genes to fold enrichment | <2.2e-16 | <2.2e-16 | -0.100017995 | -0.09751364 |
| *Esophagus-GastroesophagealJunction* | Esophagus | 250 genes to domain enrichment | <2.2e-16 | <2.2e-16 | -0.045122373 | -0.044156298 |

| | | | | | | |
|---|---|---|---|---|---|---|
| *Esophagus-Mucosa* | Esophagus | 250 genes to family enrichment | <2.2e-16 | <2.2e-16 | -0.025930699 | -0.024458803 |
| *Esophagus-Muscularis* | Esophagus | 250 genes to superfamily enrichment | <2.2e-16 | <2.2e-16 | -0.076716433 | -0.075730846 |
| *Esophagus-GastroesophagealJunction* | Esophagus | 250 genes to fold enrichment | <2.2e-16 | <2.2e-16 | -0.088694059 | -0.087722381 |
| *Esophagus-Mucosa* | Esophagus | 250 genes to domain enrichment | <2.2e-16 | <2.2e-16 | -0.045122373 | -0.044156298 |
| *Esophagus-Muscularis* | Esophagus | 250 genes to family enrichment | <2.2e-16 | <2.2e-16 | -0.025930699 | -0.024458803 |
| *Esophagus-GastroesophagealJunction* | Esophagus | 250 genes to superfamily enrichment | <2.2e-16 | <2.2e-16 | -0.076716433 | -0.075730846 |
| *Esophagus-Mucosa* | Esophagus | 250 genes to fold enrichment | <2.2e-16 | <2.2e-16 | -0.088694059 | -0.087722381 |
| *Esophagus-Muscularis* | Esophagus | 250 genes to domain enrichment | <2.2e-16 | <2.2e-16 | -0.045122373 | -0.044156298 |
| *Esophagus-GastroesophagealJunction* | Esophagus | 250 genes to family enrichment | <2.2e-16 | <2.2e-16 | -0.025930699 | -0.024458803 |
| *Esophagus-Mucosa* | Esophagus | 250 genes to superfamily enrichment | <2.2e-16 | <2.2e-16 | -0.076716433 | -0.075730846 |
| *Esophagus-Muscularis* | Esophagus | 250 genes to fold enrichment | <2.2e-16 | <2.2e-16 | -0.088694059 | -0.087722381 |
| *FallopianTube* | **FallopianTube** | **250 genes to domain enrichment** | **0.12568065** | **0.5027226** | **-0.113075532** | **0.014414251** |
| *FallopianTube* | **FallopianTube** | **250 genes to family enrichment** | **0.60985602** | **1** | **-0.08560956** | **0.051052742** |
| *FallopianTube* | FallopianTube | 250 genes to superfamily enrichment | 0.008075886 | 0.032303544 | -0.147267663 | -0.023495004 |
| *FallopianTube* | FallopianTube | 250 genes to fold enrichment | 0.000807515 | 0.003230061 | -0.166843958 | -0.047485255 |
| *Heart-AtrialAppendage* | Heart | 250 genes to domain enrichment | <2.2e-16 | <2.2e-16 | -0.038998495 | -0.037067064 |
| *Heart-LeftVentricle* | Heart | 250 genes to family enrichment | <2.2e-16 | <2.2e-16 | -0.039063731 | -0.036109969 |
| *Heart-AtrialAppendage* | Heart | 250 genes to superfamily enrichment | <2.2e-16 | <2.2e-16 | -0.077874855 | -0.075975124 |
| *Heart-LeftVentricle* | Heart | 250 genes to fold enrichment | <2.2e-16 | <2.2e-16 | -0.105194512 | -0.103314252 |
| *Heart-AtrialAppendage* | Heart | 250 genes to domain enrichment | <2.2e-16 | <2.2e-16 | -0.038998495 | -0.037067064 |
| *Heart-LeftVentricle* | Heart | 250 genes to family enrichment | <2.2e-16 | <2.2e-16 | -0.039063731 | -0.036109969 |
| *Heart-AtrialAppendage* | Heart | 250 genes to superfamily enrichment | <2.2e-16 | <2.2e-16 | -0.077874855 | -0.075975124 |
| *Heart-LeftVentricle* | Heart | 250 genes to fold enrichment | <2.2e-16 | <2.2e-16 | -0.105194512 | -0.103314252 |
| *Kidney-Cortex* | Kidney | 250 genes to domain enrichment | 1.88E-75 | 7.53E-75 | -0.074462942 | -0.060660009 |
| *Kidney-Cortex* | Kidney | 250 genes to family enrichment | 7.56E-25 | 3.02E-24 | -0.077032098 | -0.054065743 |
| *Kidney-Cortex* | Kidney | 250 genes to superfamily enrichment | 2.06E-99 | 8.22E-99 | -0.089096801 | -0.074765947 |
| *Kidney-Cortex* | Kidney | 250 genes to fold enrichment | 2.01E-179 | 8.04E-179 | -0.118406062 | -0.104646526 |
| *Liver* | Liver | 250 genes to domain enrichment | <2.2e-16 | <2.2e-16 | -0.064480734 | -0.060497482 |
| *Liver* | Liver | 250 genes to family enrichment | <2.2e-16 | <2.2e-16 | -0.087962832 | -0.083274022 |

| | | | | | | |
|---|---|---|---|---|---|---|
| *Liver* | Liver | 250 genes to superfamily enrichment | <2.2e-16 | <2.2e-16 | -0.117701556 | -0.113826162 |
| *Liver* | Liver | 250 genes to fold enrichment | <2.2e-16 | <2.2e-16 | -0.14561381 | -0.141876525 |
| *Lung* | Lung | 250 genes to domain enrichment | <2.2e-16 | <2.2e-16 | -0.083758386 | -0.082640427 |
| *Lung* | Lung | 250 genes to family enrichment | <2.2e-16 | <2.2e-16 | -0.078875079 | -0.077446412 |
| *Lung* | Lung | 250 genes to superfamily enrichment | <2.2e-16 | <2.2e-16 | -0.101911186 | -0.100809623 |
| *Lung* | Lung | 250 genes to fold enrichment | <2.2e-16 | <2.2e-16 | -0.122258023 | -0.121179476 |
| *MinorSalivaryGland* | MinorSalivaryGland | 250 genes to domain enrichment | 3.48E-235 | 1.39E-234 | -0.082727959 | -0.073644683 |
| *MinorSalivaryGland* | MinorSalivaryGland | 250 genes to family enrichment | 5.07E-102 | 2.03E-101 | -0.080813601 | -0.067859799 |
| *MinorSalivaryGland* | MinorSalivaryGland | 250 genes to superfamily enrichment | <2.2e-16 | <2.2e-16 | -0.126535469 | -0.117813318 |
| *MinorSalivaryGland* | MinorSalivaryGland | 250 genes to fold enrichment | <2.2e-16 | <2.2e-16 | -0.147252287 | -0.138841466 |
| *Muscle-Skeletal* | Muscle | 250 genes to domain enrichment | <2.2e-16 | <2.2e-16 | -0.054885488 | -0.053796707 |
| *Muscle-Skeletal* | Muscle | 250 genes to family enrichment | <2.2e-16 | <2.2e-16 | -0.070677401 | -0.069092903 |
| *Muscle-Skeletal* | Muscle | 250 genes to superfamily enrichment | <2.2e-16 | <2.2e-16 | -0.111260136 | -0.110216908 |
| *Muscle-Skeletal* | Muscle | 250 genes to fold enrichment | <2.2e-16 | <2.2e-16 | -0.145671809 | -0.14467168 |
| *Nerve-Tibial* | Nerve | 250 genes to domain enrichment | <2.2e-16 | <2.2e-16 | -0.038903251 | -0.037768982 |
| *Nerve-Tibial* | Nerve | 250 genes to family enrichment | 1.46E-94 | 5.84E-94 | -0.009966979 | -0.00824097 |
| *Nerve-Tibial* | Nerve | 250 genes to superfamily enrichment | <2.2e-16 | <2.2e-16 | -0.0566804 | -0.055527858 |
| *Nerve-Tibial* | Nerve | 250 genes to fold enrichment | <2.2e-16 | <2.2e-16 | -0.08062669 | -0.079494631 |
| *Ovary* | Ovary | 250 genes to domain enrichment | 1.43E-234 | 5.71E-234 | -0.037024399 | -0.032897278 |
| *Ovary* | Ovary | 250 genes to family enrichment | 8.61E-07 | 3.44E-06 | 0.003454259 | 0.008024309 |
| *Ovary* | Ovary | 250 genes to superfamily enrichment | <2.2e-16 | <2.2e-16 | -0.045970557 | -0.041887542 |
| *Ovary* | Ovary | 250 genes to fold enrichment | <2.2e-16 | <2.2e-16 | -0.067918626 | -0.064043782 |
| *Pancreas* | Pancreas | 250 genes to domain enrichment | <2.2e-16 | <2.2e-16 | -0.028078048 | -0.026153428 |
| *Pancreas* | Pancreas | 250 genes to family enrichment | 1.82E-171 | 7.26E-171 | -0.021969829 | -0.019114486 |
| *Pancreas* | Pancreas | 250 genes to superfamily enrichment | <2.2e-16 | <2.2e-16 | -0.067030035 | -0.065088885 |
| *Pancreas* | Pancreas | 250 genes to fold enrichment | <2.2e-16 | <2.2e-16 | -0.077352793 | -0.075447578 |
| *Pituitary* | Pituitary | 250 genes to domain enrichment | 9.60E-163 | 3.84E-162 | -0.027944034 | -0.024205168 |
| *Pituitary* | Pituitary | 250 genes to family enrichment | 1.34E-39 | 5.36E-39 | 0.017828152 | 0.024015957 |
| *Pituitary* | Pituitary | 250 genes to superfamily enrichment | 1.45E-201 | 5.79E-201 | -0.029464173 | -0.02590701 |
| *Pituitary* | Pituitary | 250 genes to fold enrichment | <2.2e-16 | <2.2e-16 | -0.059752974 | -0.056274015 |
| *Prostate* | Prostate | 250 genes to domain enrichment | <2.2e-16 | <2.2e-16 | -0.049041903 | -0.045075021 |

| | | | | | | |
|---|---|---|---|---|---|---|
| *Prostate* | Prostate | 250 genes to family enrichment | 1.41E-85 | 5.65E-85 | -0.028453138 | -0.023321055 |
| *Prostate* | Prostate | 250 genes to superfamily enrichment | <2.2e-16 | <2.2e-16 | -0.076331219 | -0.072364723 |
| *Prostate* | Prostate | 250 genes to fold enrichment | <2.2e-16 | <2.2e-16 | -0.088899881 | -0.084991865 |
| *SmallIntestine-TerminalIleum* | SmallIntestine | 250 genes to domain enrichment | <2.2e-16 | <2.2e-16 | -0.080405483 | -0.07319127 |
| *SmallIntestine-TerminalIleum* | SmallIntestine | 250 genes to family enrichment | 7.04E-141 | 2.82E-140 | -0.078613338 | -0.067771239 |
| *SmallIntestine-TerminalIleum* | SmallIntestine | 250 genes to superfamily enrichment | <2.2e-16 | <2.2e-16 | -0.138490473 | -0.131853062 |
| *SmallIntestine-TerminalIleum* | SmallIntestine | 250 genes to fold enrichment | <2.2e-16 | <2.2e-16 | -0.159250776 | -0.152757324 |
| *Spleen* | Spleen | 250 genes to domain enrichment | <2.2e-16 | <2.2e-16 | -0.072420601 | -0.068747039 |
| *Spleen* | Spleen | 250 genes to family enrichment | 9.84E-126 | 3.93E-125 | -0.04082218 | -0.034767124 |
| *Spleen* | Spleen | 250 genes to superfamily enrichment | <2.2e-16 | <2.2e-16 | -0.078023803 | -0.074271895 |
| *Spleen* | Spleen | 250 genes to fold enrichment | <2.2e-16 | <2.2e-16 | -0.105034424 | -0.101336457 |
| *Stomach* | Stomach | 250 genes to domain enrichment | <2.2e-16 | <2.2e-16 | -0.083315216 | -0.079546887 |
| *Stomach* | Stomach | 250 genes to family enrichment | 4.04E-302 | 1.61E-301 | -0.064463674 | -0.058161435 |
| *Stomach* | Stomach | 250 genes to superfamily enrichment | <2.2e-16 | <2.2e-16 | -0.128771954 | -0.125110585 |
| *Stomach* | Stomach | 250 genes to fold enrichment | <2.2e-16 | <2.2e-16 | -0.153331671 | -0.149776966 |
| *Testis* | Testis | 250 genes to domain enrichment | 2.37E-125 | 9.49E-125 | -0.029887877 | -0.025351248 |
| *Testis* | Testis | 250 genes to family enrichment | 2.46E-228 | 9.84E-228 | -0.056428839 | -0.050051355 |
| *Testis* | Testis | 250 genes to superfamily enrichment | <2.2e-16 | <2.2e-16 | -0.100561561 | -0.096276241 |
| *Testis* | Testis | 250 genes to fold enrichment | <2.2e-16 | <2.2e-16 | -0.129837092 | -0.125643193 |
| *Thyroid* | Thyroid | 250 genes to domain enrichment | <2.2e-16 | <2.2e-16 | -0.052458599 | -0.051303203 |
| *Thyroid* | Thyroid | 250 genes to family enrichment | <2.2e-16 | <2.2e-16 | -0.025687564 | -0.023901421 |
| *Thyroid* | Thyroid | 250 genes to superfamily enrichment | <2.2e-16 | <2.2e-16 | -0.074593248 | -0.073399704 |
| *Thyroid* | Thyroid | 250 genes to fold enrichment | <2.2e-16 | <2.2e-16 | -0.098953571 | -0.097788015 |
| *Uterus* | Uterus | 250 genes to domain enrichment | 3.67E-88 | 1.47E-87 | -0.026995509 | -0.022191398 |
| *Uterus* | Uterus | 250 genes to family enrichment | 8.15E-25 | 3.26E-24 | -0.018947883 | -0.012905564 |
| *Uterus* | Uterus | 250 genes to superfamily enrichment | <2.2e-16 | <2.2e-16 | -0.059834732 | -0.055038936 |
| *Uterus* | Uterus | 250 genes to fold enrichment | <2.2e-16 | <2.2e-16 | -0.069582113 | -0.064861377 |
| *Vagina* | Vagina | 250 genes to domain enrichment | 6.96957813932128e-315 | 2.78783125572851e-314 | -0.079401918 | -0.07180799 |
| *Vagina* | Vagina | 250 genes to family enrichment | 1.76E-62 | 7.04E-62 | -0.049541807 | -0.039264979 |
| *Vagina* | Vagina | 250 genes to superfamily enrichment | <2.2e-16 | <2.2e-16 | -0.114075988 | -0.106733154 |
| *Vagina* | Vagina | 250 genes to fold enrichment | <2.2e-16 | <2.2e-16 | -0.150140249 | -0.143079298 |

| | | | | | | |
|---|---|---|---|---|---|---|
| *WholeBlood* | WholeBlood | 250 genes to domain enrichment | <2.2e-16 | <2.2e-16 | -0.071853819 | -0.069687404 |
| *WholeBlood* | WholeBlood | 250 genes to family enrichment | <2.2e-16 | <2.2e-16 | -0.107745428 | -0.105580052 |
| *WholeBlood* | WholeBlood | 250 genes to superfamily enrichment | <2.2e-16 | <2.2e-16 | -0.15306531 | -0.151005687 |
| *WholeBlood* | WholeBlood | 250 genes to fold enrichment | <2.2e-16 | <2.2e-16 | -0.176427353 | -0.17440875 |
| *random* | random | 250 genes to domain enrichment | 3.67E-67 | 1.47E-66 | -0.099354361 | -0.079824366 |
| *random* | random | 250 genes to family enrichment | 1.34E-66 | 5.37E-66 | -0.093805628 | -0.075313232 |
| *random* | random | 250 genes to superfamily enrichment | 4.65E-198 | 1.86E-197 | -0.167803858 | -0.149501713 |
| *random* | random | 250 genes to fold enrichment | 8.32E-280 | 3.33E-279 | -0.204149845 | -0.186365438 |

**Table S7:** Robust GES and sGES of tissues across ARCHS4 and GTEx after outlier removal from ARCHS.

| signature | tissue | Signature Type |
|---|---|---|
| B2M | Adipose | gene |
| EEF1A1 | Adipose | gene |
| EEF1G | Adipose | gene |
| FTH1 | Adipose | gene |
| FTL | Adipose | gene |
| MT-ATP6 | Adipose | gene |
| MT-CO1 | Adipose | gene |
| MT-CO2 | Adipose | gene |
| MT-CO3 | Adipose | gene |
| MT-CYB | Adipose | gene |
| MT-ND1 | Adipose | gene |
| MT-ND4 | Adipose | gene |
| PSAP | Adipose | gene |
| RPL10 | Adipose | gene |
| RPL11 | Adipose | gene |
| RPL13A | Adipose | gene |
| RPL19 | Adipose | gene |
| RPL3 | Adipose | gene |
| RPL4 | Adipose | gene |
| RPL6 | Adipose | gene |
| RPL7 | Adipose | gene |
| RPL7A | Adipose | gene |
| RPLP0 | Adipose | gene |
| RPS11 | Adipose | gene |
| RPS20 | Adipose | gene |
| RPS6 | Adipose | gene |
| TMSB4X | Adipose | gene |

| | | |
|---|---|---|
| TPT1 | Adipose | gene |
| VIM | Adipose | gene |
| ACTG1 | Brain | gene |
| CALM2 | Brain | gene |
| GAPDH | Brain | gene |
| PEBP1 | Brain | gene |
| PSAP | Brain | gene |
| ACTB | Esophagus | gene |
| ACTG1 | Esophagus | gene |
| EEF1A1 | Esophagus | gene |
| EEF1G | Esophagus | gene |
| MT-ATP6 | Esophagus | gene |
| MT-CO1 | Esophagus | gene |
| MT-CO2 | Esophagus | gene |
| MT-CO3 | Esophagus | gene |
| MT-CYB | Esophagus | gene |
| MT-ND1 | Esophagus | gene |
| MT-ND2 | Esophagus | gene |
| MT-ND3 | Esophagus | gene |
| MT-ND4 | Esophagus | gene |
| RPL10 | Esophagus | gene |
| RPL13 | Esophagus | gene |
| RPL13A | Esophagus | gene |
| RPL14 | Esophagus | gene |
| RPL17 | Esophagus | gene |
| RPL27 | Esophagus | gene |
| RPL7A | Esophagus | gene |
| RPS18 | Esophagus | gene |
| RPS27 | Esophagus | gene |
| RPS27A | Esophagus | gene |

| | | |
|---|---|---|
| RPS6 | Esophagus | gene |
| RPS7 | Esophagus | gene |
| RPS8 | Esophagus | gene |
| TPT1 | Esophagus | gene |
| EEF1A1 | Kidney | gene |
| ALB | Liver | gene |
| AMBP | Liver | gene |
| APOA2 | Liver | gene |
| APOC3 | Liver | gene |
| APOH | Liver | gene |
| C3 | Liver | gene |
| CFB | Liver | gene |
| FGA | Liver | gene |
| FGB | Liver | gene |
| FGG | Liver | gene |
| FTL | Liver | gene |
| HP | Liver | gene |
| HPX | Liver | gene |
| MT-CO1 | Liver | gene |
| RBP4 | Liver | gene |
| SERPINA1 | Liver | gene |
| TF | Liver | gene |
| VTN | Liver | gene |
| ALDOA | Muscle | gene |
| ATP5B | Muscle | gene |
| BIN1 | Muscle | gene |
| CKM | Muscle | gene |
| DES | Muscle | gene |
| EEF1A2 | Muscle | gene |
| EEF1G | Muscle | gene |

| | | |
|---|---|---|
| EEF2 | Muscle | gene |
| GAPDH | Muscle | gene |
| HSP90AB1 | Muscle | gene |
| KLHL41 | Muscle | gene |
| MT-CO1 | Muscle | gene |
| MT-CO2 | Muscle | gene |
| MT-CO3 | Muscle | gene |
| MT-CYB | Muscle | gene |
| MT-ND1 | Muscle | gene |
| MT-ND4 | Muscle | gene |
| MT-ND5 | Muscle | gene |
| MYBPC1 | Muscle | gene |
| MYOZ1 | Muscle | gene |
| PSAP | Muscle | gene |
| RPL10 | Muscle | gene |
| RPL13A | Muscle | gene |
| RPL4 | Muscle | gene |
| RPLP0 | Muscle | gene |
| RPLP1 | Muscle | gene |
| TPT1 | Muscle | gene |
| YBX1 | Muscle | gene |
| YBX3 | Muscle | gene |
| ACTB | Ovary | gene |
| ACTG1 | Ovary | gene |
| B2M | Ovary | gene |
| BTF3 | Ovary | gene |
| EEF1A1 | Ovary | gene |
| EEF1B2 | Ovary | gene |
| EEF1G | Ovary | gene |
| EEF2 | Ovary | gene |

| | | |
|---|---|---|
| FTH1 | Ovary | gene |
| FTL | Ovary | gene |
| GAPDH | Ovary | gene |
| HSP90AB1 | Ovary | gene |
| IGFBP4 | Ovary | gene |
| IGFBP5 | Ovary | gene |
| MT-ATP6 | Ovary | gene |
| MT-CO1 | Ovary | gene |
| MT-CO2 | Ovary | gene |
| MT-CO3 | Ovary | gene |
| MT-CYB | Ovary | gene |
| MT-ND2 | Ovary | gene |
| MT-ND3 | Ovary | gene |
| MT-ND4 | Ovary | gene |
| PSAP | Ovary | gene |
| PTMA | Ovary | gene |
| RPL10 | Ovary | gene |
| RPL10A | Ovary | gene |
| RPL11 | Ovary | gene |
| RPL12 | Ovary | gene |
| RPL13 | Ovary | gene |
| RPL13A | Ovary | gene |
| RPL14 | Ovary | gene |
| RPL17 | Ovary | gene |
| RPL19 | Ovary | gene |
| RPL21 | Ovary | gene |
| RPL23 | Ovary | gene |
| RPL24 | Ovary | gene |
| RPL26 | Ovary | gene |
| RPL27 | Ovary | gene |

| | | |
|---|---|---|
| RPL29 | Ovary | gene |
| RPL3 | Ovary | gene |
| RPL32 | Ovary | gene |
| RPL35A | Ovary | gene |
| RPL4 | Ovary | gene |
| RPL41 | Ovary | gene |
| RPL5 | Ovary | gene |
| RPL6 | Ovary | gene |
| RPL7 | Ovary | gene |
| RPL7A | Ovary | gene |
| RPLP0 | Ovary | gene |
| RPS11 | Ovary | gene |
| RPS12 | Ovary | gene |
| RPS13 | Ovary | gene |
| RPS18 | Ovary | gene |
| RPS20 | Ovary | gene |
| RPS24 | Ovary | gene |
| RPS25 | Ovary | gene |
| RPS27 | Ovary | gene |
| RPS27A | Ovary | gene |
| RPS3A | Ovary | gene |
| RPS4X | Ovary | gene |
| RPS6 | Ovary | gene |
| RPS7 | Ovary | gene |
| RPS8 | Ovary | gene |
| RPSA | Ovary | gene |
| SERPING1 | Ovary | gene |
| TMSB10 | Ovary | gene |
| TMSB4X | Ovary | gene |
| TPT1 | Ovary | gene |

| | | |
|---|---|---|
| VIM | Ovary | gene |
| EEF1A1 | Prostate | gene |
| EEF1G | Prostate | gene |
| GAPDH | Prostate | gene |
| HSP90AB1 | Prostate | gene |
| RPL19 | Prostate | gene |
| RPL3 | Prostate | gene |
| RPL4 | Prostate | gene |
| RPL41 | Prostate | gene |
| RPL6 | Prostate | gene |
| RPL7A | Prostate | gene |
| RPLP0 | Prostate | gene |
| RPS18 | Prostate | gene |
| RPS6 | Prostate | gene |
| ACTB | SmallIntestine | gene |
| EEF1A1 | SmallIntestine | gene |
| FTH1 | SmallIntestine | gene |
| MT-CO1 | SmallIntestine | gene |
| MT-CO3 | SmallIntestine | gene |
| TPT1 | SmallIntestine | gene |
| ACTB | Spleen | gene |
| ACTG1 | Spleen | gene |
| B2M | Spleen | gene |
| EEF1A1 | Spleen | gene |
| HLA-B | Spleen | gene |
| HLA-E | Spleen | gene |
| MT-CO3 | Spleen | gene |
| MT-ND4 | Spleen | gene |
| RPL13A | Spleen | gene |
| RPL3 | Spleen | gene |

| | | |
|---|---|---|
| RPL4 | Spleen | gene |
| TMSB4X | Spleen | gene |
| TPT1 | Spleen | gene |
| EEF1A1 | Stomach | gene |
| MT-ATP6 | Stomach | gene |
| MT-CO1 | Stomach | gene |
| MT-CO3 | Stomach | gene |
| MT-CYB | Stomach | gene |
| MT-ND1 | Stomach | gene |
| MT-ND3 | Stomach | gene |
| MT-ND4 | Stomach | gene |
| RPL10 | Stomach | gene |
| RPL13 | Stomach | gene |
| RPL13A | Stomach | gene |
| RPL41 | Stomach | gene |
| RPS18 | Stomach | gene |
| RPS27 | Stomach | gene |
| RPS27A | Stomach | gene |
| RPS3A | Stomach | gene |
| RPS6 | Stomach | gene |
| ACTB | Thyroid | gene |
| ACTG1 | Thyroid | gene |
| B2M | Thyroid | gene |
| EEF1A1 | Thyroid | gene |
| EEF1G | Thyroid | gene |
| EEF2 | Thyroid | gene |
| FTH1 | Thyroid | gene |
| GAPDH | Thyroid | gene |
| HSP90AB1 | Thyroid | gene |
| MT-ATP6 | Thyroid | gene |

| | | |
|---|---|---|
| MT-CO1 | Thyroid | gene |
| MT-CO2 | Thyroid | gene |
| MT-CO3 | Thyroid | gene |
| MT-CYB | Thyroid | gene |
| MT-ND1 | Thyroid | gene |
| MT-ND2 | Thyroid | gene |
| MT-ND3 | Thyroid | gene |
| MT-ND4 | Thyroid | gene |
| PSAP | Thyroid | gene |
| PTMA | Thyroid | gene |
| RPL10 | Thyroid | gene |
| RPL13 | Thyroid | gene |
| RPL13A | Thyroid | gene |
| RPL17 | Thyroid | gene |
| RPL19 | Thyroid | gene |
| RPL3 | Thyroid | gene |
| RPL4 | Thyroid | gene |
| RPL5 | Thyroid | gene |
| RPL6 | Thyroid | gene |
| RPL7A | Thyroid | gene |
| RPL8 | Thyroid | gene |
| RPLP0 | Thyroid | gene |
| RPS11 | Thyroid | gene |
| RPS4X | Thyroid | gene |
| RPS6 | Thyroid | gene |
| RPS8 | Thyroid | gene |
| TPT1 | Thyroid | gene |
| ACTB | Vagina | gene |
| ACTG1 | Vagina | gene |
| EEF1A1 | Vagina | gene |

| | | |
|---|---|---|
| EEF1G | Vagina | gene |
| EEF2 | Vagina | gene |
| GAPDH | Vagina | gene |
| GSTP1 | Vagina | gene |
| HSPB1 | Vagina | gene |
| MT-ATP6 | Vagina | gene |
| MT-ATP8 | Vagina | gene |
| MT-CO1 | Vagina | gene |
| MT-CO2 | Vagina | gene |
| MT-CO3 | Vagina | gene |
| MT-CYB | Vagina | gene |
| MT-ND1 | Vagina | gene |
| MT-ND2 | Vagina | gene |
| MT-ND3 | Vagina | gene |
| MT-ND4 | Vagina | gene |
| MT-ND5 | Vagina | gene |
| RPL10 | Vagina | gene |
| RPL10A | Vagina | gene |
| RPL11 | Vagina | gene |
| RPL13 | Vagina | gene |
| RPL13A | Vagina | gene |
| RPL17 | Vagina | gene |
| RPL19 | Vagina | gene |
| RPL26 | Vagina | gene |
| RPL29 | Vagina | gene |
| RPL3 | Vagina | gene |
| RPL35 | Vagina | gene |
| RPL4 | Vagina | gene |
| RPL41 | Vagina | gene |
| RPL5 | Vagina | gene |

| | | |
|---|---|---|
| RPL6 | Vagina | gene |
| RPL7 | Vagina | gene |
| RPL7A | Vagina | gene |
| RPL8 | Vagina | gene |
| RPLP0 | Vagina | gene |
| RPLP1 | Vagina | gene |
| RPLP2 | Vagina | gene |
| RPS10 | Vagina | gene |
| RPS11 | Vagina | gene |
| RPS12 | Vagina | gene |
| RPS14 | Vagina | gene |
| RPS18 | Vagina | gene |
| RPS2 | Vagina | gene |
| RPS27 | Vagina | gene |
| RPS3A | Vagina | gene |
| RPS4X | Vagina | gene |
| RPS5 | Vagina | gene |
| RPS6 | Vagina | gene |
| RPS7 | Vagina | gene |
| RPS8 | Vagina | gene |
| RPSA | Vagina | gene |
| TPT1 | Vagina | gene |
| ACTB | WholeBlood | gene |
| FTL | WholeBlood | gene |
| IPR000077 | Adipose | domain |
| IPR000196 | Adipose | domain |
| IPR000298 | Adipose | domain |
| IPR000568 | Adipose | domain |
| IPR000719 | Adipose | domain |
| IPR000836 | Adipose | domain |

| | | |
|---|---|---|
| IPR000961 | Adipose | domain |
| IPR001133 | Adipose | domain |
| IPR001152 | Adipose | domain |
| IPR001662 | Adipose | domain |
| IPR001694 | Adipose | domain |
| IPR001790 | Adipose | domain |
| IPR002048 | Adipose | domain |
| IPR002350 | Adipose | domain |
| IPR002429 | Adipose | domain |
| IPR003599 | Adipose | domain |
| IPR004000 | Adipose | domain |
| IPR004038 | Adipose | domain |
| IPR004045 | Adipose | domain |
| IPR004160 | Adipose | domain |
| IPR004161 | Adipose | domain |
| IPR005225 | Adipose | domain |
| IPR005568 | Adipose | domain |
| IPR005797 | Adipose | domain |
| IPR005798 | Adipose | domain |
| IPR005822 | Adipose | domain |
| IPR005824 | Adipose | domain |
| IPR007110 | Adipose | domain |
| IPR007740 | Adipose | domain |
| IPR008331 | Adipose | domain |
| IPR009040 | Adipose | domain |
| IPR009703 | Adipose | domain |
| IPR010987 | Adipose | domain |
| IPR011759 | Adipose | domain |
| IPR012988 | Adipose | domain |
| IPR014222 | Adipose | domain |

| | | |
|---|---|---|
| IPR016082 | Adipose | domain |
| IPR016180 | Adipose | domain |
| IPR018499 | Adipose | domain |
| IPR023616 | Adipose | domain |
| IPR027486 | Adipose | domain |
| IPR028139 | Adipose | domain |
| IPR029099 | Adipose | domain |
| IPR031309 | Adipose | domain |
| IPR031310 | Adipose | domain |
| IPR032440 | Adipose | domain |
| IPR034737 | Adipose | domain |
| IPR035808 | Adipose | domain |
| No-Domain | Adipose | domain |
| IPR000719 | Brain | domain |
| IPR000836 | Brain | domain |
| IPR002048 | Brain | domain |
| IPR003594 | Brain | domain |
| IPR004000 | Brain | domain |
| IPR004160 | Brain | domain |
| IPR004161 | Brain | domain |
| IPR005225 | Brain | domain |
| IPR008331 | Brain | domain |
| IPR008914 | Brain | domain |
| IPR009040 | Brain | domain |
| IPR013087 | Brain | domain |
| IPR020828 | Brain | domain |
| IPR020829 | Brain | domain |
| IPR023410 | Brain | domain |
| IPR029099 | Brain | domain |

| | | |
|---|---|---|
| No-Domain | Brain | domain |
| IPR002048 | Colon | domain |
| IPR005225 | Colon | domain |
| No-Domain | Colon | domain |
| IPR000298 | Esophagus | domain |
| IPR000440 | Esophagus | domain |
| IPR000554 | Esophagus | domain |
| IPR000568 | Esophagus | domain |
| IPR000592 | Esophagus | domain |
| IPR000626 | Esophagus | domain |
| IPR000719 | Esophagus | domain |
| IPR000836 | Esophagus | domain |
| IPR000961 | Esophagus | domain |
| IPR001063 | Esophagus | domain |
| IPR001133 | Esophagus | domain |
| IPR001152 | Esophagus | domain |
| IPR001662 | Esophagus | domain |
| IPR001694 | Esophagus | domain |
| IPR001750 | Esophagus | domain |
| IPR002048 | Esophagus | domain |
| IPR002429 | Esophagus | domain |
| IPR002784 | Esophagus | domain |
| IPR002906 | Esophagus | domain |
| IPR004000 | Esophagus | domain |
| IPR004038 | Esophagus | domain |
| IPR004045 | Esophagus | domain |
| IPR004160 | Esophagus | domain |
| IPR004161 | Esophagus | domain |
| IPR005225 | Esophagus | domain |

| | | |
|---|---|---|
| IPR005797 | Esophagus | domain |
| IPR005798 | Esophagus | domain |
| IPR005822 | Esophagus | domain |
| IPR006802 | Esophagus | domain |
| IPR007110 | Esophagus | domain |
| IPR008331 | Esophagus | domain |
| IPR009040 | Esophagus | domain |
| IPR010933 | Esophagus | domain |
| IPR010987 | Esophagus | domain |
| IPR011759 | Esophagus | domain |
| IPR014222 | Esophagus | domain |
| IPR016180 | Esophagus | domain |
| IPR020040 | Esophagus | domain |
| IPR021131 | Esophagus | domain |
| IPR022309 | Esophagus | domain |
| IPR023616 | Esophagus | domain |
| IPR028139 | Esophagus | domain |
| IPR029099 | Esophagus | domain |
| IPR034737 | Esophagus | domain |
| No-Domain | Esophagus | domain |
| IPR000719 | Heart | domain |
| IPR001606 | Heart | domain |
| IPR001609 | Heart | domain |
| IPR002048 | Heart | domain |
| IPR003599 | Heart | domain |
| IPR004161 | Heart | domain |
| IPR005225 | Heart | domain |
| IPR007110 | Heart | domain |
| IPR020683 | Heart | domain |

| | | |
|---|---|---|
| No-Domain | Heart | domain |
| IPR004160 | Kidney | domain |
| IPR004161 | Kidney | domain |
| IPR005225 | Kidney | domain |
| No-Domain | Kidney | domain |
| IPR000436 | Liver | domain |
| IPR000566 | Liver | domain |
| IPR000568 | Liver | domain |
| IPR000585 | Liver | domain |
| IPR001164 | Liver | domain |
| IPR001254 | Liver | domain |
| IPR001590 | Liver | domain |
| IPR001750 | Liver | domain |
| IPR001762 | Liver | domain |
| IPR002035 | Liver | domain |
| IPR002181 | Liver | domain |
| IPR002223 | Liver | domain |
| IPR002870 | Liver | domain |
| IPR005225 | Liver | domain |
| IPR006586 | Liver | domain |
| IPR006801 | Liver | domain |
| IPR007110 | Liver | domain |
| IPR008278 | Liver | domain |
| IPR008331 | Liver | domain |
| IPR008403 | Liver | domain |
| IPR009040 | Liver | domain |
| IPR012290 | Liver | domain |
| IPR012617 | Liver | domain |
| IPR013649 | Liver | domain |

| | | |
|---|---|---|
| IPR014760 | Liver | domain |
| IPR015104 | Liver | domain |
| IPR018484 | Liver | domain |
| IPR018485 | Liver | domain |
| IPR023616 | Liver | domain |
| IPR023796 | Liver | domain |
| IPR025160 | Liver | domain |
| IPR002048 | Lung | domain |
| IPR005225 | Lung | domain |
| No-Domain | Lung | domain |
| IPR000194 | Muscle | domain |
| IPR000210 | Muscle | domain |
| IPR000298 | Muscle | domain |
| IPR000719 | Muscle | domain |
| IPR000741 | Muscle | domain |
| IPR000836 | Muscle | domain |
| IPR001133 | Muscle | domain |
| IPR001452 | Muscle | domain |
| IPR001516 | Muscle | domain |
| IPR001606 | Muscle | domain |
| IPR001662 | Muscle | domain |
| IPR001694 | Muscle | domain |
| IPR001715 | Muscle | domain |
| IPR001750 | Muscle | domain |
| IPR001781 | Muscle | domain |
| IPR001790 | Muscle | domain |
| IPR001870 | Muscle | domain |
| IPR001878 | Muscle | domain |
| IPR001978 | Muscle | domain |
| IPR002048 | Muscle | domain |

| | | |
|---|---|---|
| IPR002068 | Muscle | domain |
| IPR002429 | Muscle | domain |
| IPR003090 | Muscle | domain |
| IPR003593 | Muscle | domain |
| IPR003594 | Muscle | domain |
| IPR003599 | Muscle | domain |
| IPR003877 | Muscle | domain |
| IPR003961 | Muscle | domain |
| IPR004000 | Muscle | domain |
| IPR004045 | Muscle | domain |
| IPR004100 | Muscle | domain |
| IPR004148 | Muscle | domain |
| IPR004160 | Muscle | domain |
| IPR004161 | Muscle | domain |
| IPR005225 | Muscle | domain |
| IPR005797 | Muscle | domain |
| IPR005798 | Muscle | domain |
| IPR005822 | Muscle | domain |
| IPR007110 | Muscle | domain |
| IPR007740 | Muscle | domain |
| IPR008438 | Muscle | domain |
| IPR010934 | Muscle | domain |
| IPR010987 | Muscle | domain |
| IPR011129 | Muscle | domain |
| IPR011705 | Muscle | domain |
| IPR011759 | Muscle | domain |
| IPR013098 | Muscle | domain |
| IPR014222 | Muscle | domain |
| IPR016180 | Muscle | domain |
| IPR019410 | Muscle | domain |

| | | |
|---|---|---|
| IPR020828 | Muscle | domain |
| IPR020829 | Muscle | domain |
| IPR022413 | Muscle | domain |
| IPR022414 | Muscle | domain |
| IPR023616 | Muscle | domain |
| IPR027534 | Muscle | domain |
| IPR028139 | Muscle | domain |
| IPR029099 | Muscle | domain |
| IPR029426 | Muscle | domain |
| IPR034737 | Muscle | domain |
| No-Domain | Muscle | domain |
| IPR000077 | Ovary | domain |
| IPR000196 | Ovary | domain |
| IPR000218 | Ovary | domain |
| IPR000298 | Ovary | domain |
| IPR000440 | Ovary | domain |
| IPR000504 | Ovary | domain |
| IPR000554 | Ovary | domain |
| IPR000568 | Ovary | domain |
| IPR000592 | Ovary | domain |
| IPR000626 | Ovary | domain |
| IPR000716 | Ovary | domain |
| IPR000719 | Ovary | domain |
| IPR000836 | Ovary | domain |
| IPR000867 | Ovary | domain |
| IPR000961 | Ovary | domain |
| IPR001063 | Ovary | domain |
| IPR001133 | Ovary | domain |
| IPR001134 | Ovary | domain |
| IPR001147 | Ovary | domain |

| | | |
|---|---|---|
| IPR001152 | Ovary | domain |
| IPR001515 | Ovary | domain |
| IPR001593 | Ovary | domain |
| IPR001650 | Ovary | domain |
| IPR001662 | Ovary | domain |
| IPR001750 | Ovary | domain |
| IPR001780 | Ovary | domain |
| IPR001790 | Ovary | domain |
| IPR001870 | Ovary | domain |
| IPR002035 | Ovary | domain |
| IPR002048 | Ovary | domain |
| IPR002429 | Ovary | domain |
| IPR002673 | Ovary | domain |
| IPR002784 | Ovary | domain |
| IPR002906 | Ovary | domain |
| IPR002942 | Ovary | domain |
| IPR003594 | Ovary | domain |
| IPR003599 | Ovary | domain |
| IPR003877 | Ovary | domain |
| IPR004000 | Ovary | domain |
| IPR004038 | Ovary | domain |
| IPR004045 | Ovary | domain |
| IPR004160 | Ovary | domain |
| IPR004161 | Ovary | domain |
| IPR004931 | Ovary | domain |
| IPR004977 | Ovary | domain |
| IPR005225 | Ovary | domain |
| IPR005568 | Ovary | domain |
| IPR005797 | Ovary | domain |
| IPR005798 | Ovary | domain |

| | | |
|---|---|---|
| IPR005822 | Ovary | domain |
| IPR005824 | Ovary | domain |
| IPR006574 | Ovary | domain |
| IPR006802 | Ovary | domain |
| IPR007110 | Ovary | domain |
| IPR007740 | Ovary | domain |
| IPR007836 | Ovary | domain |
| IPR008331 | Ovary | domain |
| IPR009040 | Ovary | domain |
| IPR009703 | Ovary | domain |
| IPR010579 | Ovary | domain |
| IPR010933 | Ovary | domain |
| IPR010987 | Ovary | domain |
| IPR011017 | Ovary | domain |
| IPR011161 | Ovary | domain |
| IPR011759 | Ovary | domain |
| IPR012606 | Ovary | domain |
| IPR012988 | Ovary | domain |
| IPR013843 | Ovary | domain |
| IPR013845 | Ovary | domain |
| IPR014001 | Ovary | domain |
| IPR014014 | Ovary | domain |
| IPR014038 | Ovary | domain |
| IPR014222 | Ovary | domain |
| IPR016082 | Ovary | domain |
| IPR016180 | Ovary | domain |
| IPR018472 | Ovary | domain |
| IPR018940 | Ovary | domain |
| IPR019410 | Ovary | domain |
| IPR020040 | Ovary | domain |

| | | |
|---|---|---|
| IPR020783 | Ovary | domain |
| IPR020784 | Ovary | domain |
| IPR020828 | Ovary | domain |
| IPR020829 | Ovary | domain |
| IPR021131 | Ovary | domain |
| IPR022309 | Ovary | domain |
| IPR023616 | Ovary | domain |
| IPR023796 | Ovary | domain |
| IPR026146 | Ovary | domain |
| IPR027486 | Ovary | domain |
| IPR028139 | Ovary | domain |
| IPR028364 | Ovary | domain |
| IPR029099 | Ovary | domain |
| IPR029426 | Ovary | domain |
| IPR031309 | Ovary | domain |
| IPR031310 | Ovary | domain |
| IPR032277 | Ovary | domain |
| IPR032440 | Ovary | domain |
| IPR034737 | Ovary | domain |
| IPR035808 | Ovary | domain |
| No-Domain | Ovary | domain |
| IPR000077 | Prostate | domain |
| IPR000196 | Prostate | domain |
| IPR000719 | Prostate | domain |
| IPR000836 | Prostate | domain |
| IPR000961 | Prostate | domain |
| IPR001650 | Prostate | domain |
| IPR001662 | Prostate | domain |
| IPR001790 | Prostate | domain |
| IPR002048 | Prostate | domain |

| | | |
|---|---|---|
| IPR003594 | Prostate | domain |
| IPR004000 | Prostate | domain |
| IPR004038 | Prostate | domain |
| IPR004045 | Prostate | domain |
| IPR004160 | Prostate | domain |
| IPR004161 | Prostate | domain |
| IPR005225 | Prostate | domain |
| IPR005568 | Prostate | domain |
| IPR006802 | Prostate | domain |
| IPR007740 | Prostate | domain |
| IPR007836 | Prostate | domain |
| IPR010987 | Prostate | domain |
| IPR014001 | Prostate | domain |
| IPR014014 | Prostate | domain |
| IPR020828 | Prostate | domain |
| IPR020829 | Prostate | domain |
| IPR029099 | Prostate | domain |
| No-Domain | Prostate | domain |
| IPR000298 | SmallIntestine | domain |
| IPR000716 | SmallIntestine | domain |
| IPR000719 | SmallIntestine | domain |
| IPR002048 | SmallIntestine | domain |
| IPR004000 | SmallIntestine | domain |
| IPR004045 | SmallIntestine | domain |
| IPR004160 | SmallIntestine | domain |
| IPR004161 | SmallIntestine | domain |
| IPR005225 | SmallIntestine | domain |
| IPR007110 | SmallIntestine | domain |
| IPR008331 | SmallIntestine | domain |
| IPR009040 | SmallIntestine | domain |

| | | |
|---|---|---|
| IPR010987 | SmallIntestine | domain |
| IPR023616 | SmallIntestine | domain |
| IPR034737 | SmallIntestine | domain |
| No-Domain | SmallIntestine | domain |
| IPR000077 | Spleen | domain |
| IPR000298 | Spleen | domain |
| IPR000504 | Spleen | domain |
| IPR000719 | Spleen | domain |
| IPR001133 | Spleen | domain |
| IPR001152 | Spleen | domain |
| IPR001650 | Spleen | domain |
| IPR002048 | Spleen | domain |
| IPR003599 | Spleen | domain |
| IPR004000 | Spleen | domain |
| IPR004160 | Spleen | domain |
| IPR004161 | Spleen | domain |
| IPR005225 | Spleen | domain |
| IPR005822 | Spleen | domain |
| IPR007110 | Spleen | domain |
| IPR007740 | Spleen | domain |
| IPR010579 | Spleen | domain |
| IPR011161 | Spleen | domain |
| IPR014001 | Spleen | domain |
| IPR028139 | Spleen | domain |
| IPR034737 | Spleen | domain |
| No-Domain | Spleen | domain |
| IPR000298 | Stomach | domain |
| IPR000440 | Stomach | domain |
| IPR000568 | Stomach | domain |

| | | |
|---|---|---|
| IPR000592 | Stomach | domain |
| IPR000626 | Stomach | domain |
| IPR000719 | Stomach | domain |
| IPR000961 | Stomach | domain |
| IPR001133 | Stomach | domain |
| IPR001593 | Stomach | domain |
| IPR001694 | Stomach | domain |
| IPR001750 | Stomach | domain |
| IPR002906 | Stomach | domain |
| IPR004000 | Stomach | domain |
| IPR004038 | Stomach | domain |
| IPR004160 | Stomach | domain |
| IPR004161 | Stomach | domain |
| IPR005225 | Stomach | domain |
| IPR005797 | Stomach | domain |
| IPR005798 | Stomach | domain |
| IPR005822 | Stomach | domain |
| IPR006802 | Stomach | domain |
| IPR007110 | Stomach | domain |
| IPR007836 | Stomach | domain |
| IPR016180 | Stomach | domain |
| IPR021131 | Stomach | domain |
| IPR023616 | Stomach | domain |
| IPR028139 | Stomach | domain |
| No-Domain | Stomach | domain |
| IPR000077 | Thyroid | domain |
| IPR000196 | Thyroid | domain |
| IPR000298 | Thyroid | domain |
| IPR000440 | Thyroid | domain |
| IPR000568 | Thyroid | domain |

| | | |
|---|---|---|
| IPR000719 | Thyroid | domain |
| IPR000836 | Thyroid | domain |
| IPR000961 | Thyroid | domain |
| IPR001063 | Thyroid | domain |
| IPR001133 | Thyroid | domain |
| IPR001662 | Thyroid | domain |
| IPR001694 | Thyroid | domain |
| IPR001750 | Thyroid | domain |
| IPR001790 | Thyroid | domain |
| IPR002048 | Thyroid | domain |
| IPR002429 | Thyroid | domain |
| IPR002942 | Thyroid | domain |
| IPR003594 | Thyroid | domain |
| IPR003599 | Thyroid | domain |
| IPR004000 | Thyroid | domain |
| IPR004038 | Thyroid | domain |
| IPR004045 | Thyroid | domain |
| IPR004160 | Thyroid | domain |
| IPR004161 | Thyroid | domain |
| IPR004931 | Thyroid | domain |
| IPR005225 | Thyroid | domain |
| IPR005568 | Thyroid | domain |
| IPR005797 | Thyroid | domain |
| IPR005798 | Thyroid | domain |
| IPR005822 | Thyroid | domain |
| IPR005824 | Thyroid | domain |
| IPR007110 | Thyroid | domain |
| IPR007740 | Thyroid | domain |
| IPR008331 | Thyroid | domain |
| IPR009040 | Thyroid | domain |

| | | |
|---|---|---|
| IPR010933 | Thyroid | domain |
| IPR010987 | Thyroid | domain |
| IPR011759 | Thyroid | domain |
| IPR013843 | Thyroid | domain |
| IPR013845 | Thyroid | domain |
| IPR014222 | Thyroid | domain |
| IPR016180 | Thyroid | domain |
| IPR018472 | Thyroid | domain |
| IPR019410 | Thyroid | domain |
| IPR020683 | Thyroid | domain |
| IPR020828 | Thyroid | domain |
| IPR020829 | Thyroid | domain |
| IPR022309 | Thyroid | domain |
| IPR023616 | Thyroid | domain |
| IPR028139 | Thyroid | domain |
| IPR029099 | Thyroid | domain |
| IPR029426 | Thyroid | domain |
| IPR032277 | Thyroid | domain |
| IPR032440 | Thyroid | domain |
| IPR034737 | Thyroid | domain |
| No-Domain | Thyroid | domain |
| IPR000077 | Vagina | domain |
| IPR000196 | Vagina | domain |
| IPR000298 | Vagina | domain |
| IPR000440 | Vagina | domain |
| IPR000504 | Vagina | domain |
| IPR000554 | Vagina | domain |
| IPR000568 | Vagina | domain |
| IPR000592 | Vagina | domain |
| IPR000716 | Vagina | domain |

| | | |
|---|---|---|
| IPR000719 | Vagina | domain |
| IPR000836 | Vagina | domain |
| IPR000961 | Vagina | domain |
| IPR001063 | Vagina | domain |
| IPR001133 | Vagina | domain |
| IPR001254 | Vagina | domain |
| IPR001421 | Vagina | domain |
| IPR001516 | Vagina | domain |
| IPR001593 | Vagina | domain |
| IPR001662 | Vagina | domain |
| IPR001694 | Vagina | domain |
| IPR001750 | Vagina | domain |
| IPR001780 | Vagina | domain |
| IPR001790 | Vagina | domain |
| IPR001971 | Vagina | domain |
| IPR002048 | Vagina | domain |
| IPR002068 | Vagina | domain |
| IPR002429 | Vagina | domain |
| IPR002673 | Vagina | domain |
| IPR002942 | Vagina | domain |
| IPR004000 | Vagina | domain |
| IPR004038 | Vagina | domain |
| IPR004045 | Vagina | domain |
| IPR004160 | Vagina | domain |
| IPR004161 | Vagina | domain |
| IPR005225 | Vagina | domain |
| IPR005326 | Vagina | domain |
| IPR005568 | Vagina | domain |
| IPR005797 | Vagina | domain |
| IPR005798 | Vagina | domain |

| | | |
|---|---|---|
| IPR005822 | Vagina | domain |
| IPR005824 | Vagina | domain |
| IPR006802 | Vagina | domain |
| IPR007110 | Vagina | domain |
| IPR007740 | Vagina | domain |
| IPR007836 | Vagina | domain |
| IPR010933 | Vagina | domain |
| IPR010934 | Vagina | domain |
| IPR010987 | Vagina | domain |
| IPR011759 | Vagina | domain |
| IPR012988 | Vagina | domain |
| IPR013843 | Vagina | domain |
| IPR013845 | Vagina | domain |
| IPR014222 | Vagina | domain |
| IPR016082 | Vagina | domain |
| IPR016180 | Vagina | domain |
| IPR018472 | Vagina | domain |
| IPR019410 | Vagina | domain |
| IPR020040 | Vagina | domain |
| IPR020828 | Vagina | domain |
| IPR020829 | Vagina | domain |
| IPR021131 | Vagina | domain |
| IPR022309 | Vagina | domain |
| IPR023573 | Vagina | domain |
| IPR023616 | Vagina | domain |
| IPR023798 | Vagina | domain |
| IPR027534 | Vagina | domain |
| IPR028139 | Vagina | domain |
| IPR028364 | Vagina | domain |
| IPR029099 | Vagina | domain |

| | | |
|---|---|---|
| IPR029426 | Vagina | domain |
| IPR031309 | Vagina | domain |
| IPR031310 | Vagina | domain |
| IPR032277 | Vagina | domain |
| IPR032440 | Vagina | domain |
| IPR034737 | Vagina | domain |
| IPR035808 | Vagina | domain |
| No-Domain | Vagina | domain |
| IPR000719 | WholeBlood | domain |
| IPR002048 | WholeBlood | domain |
| IPR004000 | WholeBlood | domain |
| IPR005225 | WholeBlood | domain |
| IPR007110 | WholeBlood | domain |
| IPR008331 | WholeBlood | domain |
| IPR009040 | WholeBlood | domain |
| a.135.1.1 | Adipose | family |
| a.25.1.0 | Adipose | family |
| a.39.1.5 | Adipose | family |
| a.45.1.1 | Adipose | family |
| b.1.1.2 | Adipose | family |
| b.34.5.7 | Adipose | family |
| b.40.4.5 | Adipose | family |
| b.43.3.1 | Adipose | family |
| b.44.1.0 | Adipose | family |
| b.6.1.3 | Adipose | family |
| b.88.1.2 | Adipose | family |
| c.21.1.1 | Adipose | family |
| c.37.1.0 | Adipose | family |
| c.47.1.5 | Adipose | family |
| c.55.1.1 | Adipose | family |

| | | |
|---|---|---|
| c.61.1.0 | Adipose | family |
| c.61.1.2 | Adipose | family |
| d.144.1.0 | Adipose | family |
| d.41.4.1 | Adipose | family |
| d.58.46.1 | Adipose | family |
| d.58.62.1 | Adipose | family |
| d.64.1.1 | Adipose | family |
| d.79.3.1 | Adipose | family |
| f.17.2.1 | Adipose | family |
| f.18.1.1 | Adipose | family |
| f.21.1.2 | Adipose | family |
| f.25.1.1 | Adipose | family |
| f.32.1.1 | Adipose | family |
| g.68.1.1 | Adipose | family |
| No-Fold | Adipose | family |
| a.118.7.1 | Brain | family |
| a.25.1.0 | Brain | family |
| a.39.1.5 | Brain | family |
| b.17.1.1 | Brain | family |
| b.43.3.1 | Brain | family |
| b.44.1.0 | Brain | family |
| c.2.1.3 | Brain | family |
| c.37.1.0 | Brain | family |
| c.55.1.1 | Brain | family |
| c.61.1.0 | Brain | family |
| c.61.1.2 | Brain | family |
| d.122.1.1 | Brain | family |
| d.144.1.0 | Brain | family |
| d.81.1.1 | Brain | family |
| No-Fold | Brain | family |

| | | |
|---|---|---|
| a.39.1.5 | Colon | family |
| c.37.1.0 | Colon | family |
| No-Fold | Colon | family |
| a.25.1.0 | Esophagus | family |
| a.39.1.5 | Esophagus | family |
| a.45.1.1 | Esophagus | family |
| b.29.1.0 | Esophagus | family |
| b.34.5.7 | Esophagus | family |
| b.40.4.5 | Esophagus | family |
| b.43.3.1 | Esophagus | family |
| b.44.1.0 | Esophagus | family |
| b.6.1.3 | Esophagus | family |
| b.88.1.2 | Esophagus | family |
| c.21.1.1 | Esophagus | family |
| c.37.1.0 | Esophagus | family |
| c.47.1.5 | Esophagus | family |
| c.55.1.1 | Esophagus | family |
| c.61.1.0 | Esophagus | family |
| c.61.1.2 | Esophagus | family |
| d.141.1.1 | Esophagus | family |
| d.144.1.0 | Esophagus | family |
| d.41.4.1 | Esophagus | family |
| d.55.1.1 | Esophagus | family |
| d.58.46.1 | Esophagus | family |
| d.79.3.1 | Esophagus | family |
| f.17.2.1 | Esophagus | family |
| f.18.1.1 | Esophagus | family |
| f.21.1.2 | Esophagus | family |
| f.25.1.1 | Esophagus | family |
| f.32.1.1 | Esophagus | family |

| | | |
|---|---|---|
| g.41.8.4 | Esophagus | family |
| No-Fold | Esophagus | family |
| a.39.1.0 | Heart | family |
| b.1.1.0 | Heart | family |
| b.43.3.1 | Heart | family |
| b.44.1.0 | Heart | family |
| c.37.1.8 | Heart | family |
| c.37.1.9 | Heart | family |
| c.55.1.1 | Heart | family |
| d.211.1.1 | Heart | family |
| No-Fold | Heart | family |
| b.43.3.1 | Kidney | family |
| b.44.1.0 | Kidney | family |
| c.37.1.0 | Kidney | family |
| d.64.1.1 | Kidney | family |
| No-Fold | Kidney | family |
| a.126.1.1 | Liver | family |
| a.24.1.1 | Liver | family |
| a.25.1.0 | Liver | family |
| b.1.1.2 | Liver | family |
| b.47.1.2 | Liver | family |
| b.60.1.0 | Liver | family |
| b.60.1.1 | Liver | family |
| c.37.1.0 | Liver | family |
| c.55.1.10 | Liver | family |
| c.62.1.1 | Liver | family |
| d.171.1.1 | Liver | family |
| d.92.1.9 | Liver | family |
| e.1.1.0 | Liver | family |
| f.18.1.1 | Liver | family |

| | | |
|---|---|---|
| g.18.1.1 | Liver | family |
| g.45.1.1 | Liver | family |
| g.8.1.0 | Liver | family |
| No-Fold | Liver | family |
| c.37.1.0 | Lung | family |
| d.64.1.1 | Lung | family |
| No-Fold | Lung | family |
| a.238.1.0 | Muscle | family |
| a.26.1.2 | Muscle | family |
| a.39.1.0 | Muscle | family |
| a.40.1.1 | Muscle | family |
| a.45.1.1 | Muscle | family |
| a.83.1.0 | Muscle | family |
| b.1.1.0 | Muscle | family |
| b.15.1.1 | Muscle | family |
| b.29.1.0 | Muscle | family |
| b.34.12.1 | Muscle | family |
| b.40.4.5 | Muscle | family |
| b.43.3.1 | Muscle | family |
| b.44.1.0 | Muscle | family |
| b.49.1.1 | Muscle | family |
| b.6.1.3 | Muscle | family |
| b.68.11.0 | Muscle | family |
| b.88.1.2 | Muscle | family |
| c.1.10.1 | Muscle | family |
| c.2.1.3 | Muscle | family |
| c.21.1.1 | Muscle | family |
| c.37.1.0 | Muscle | family |
| c.37.1.8 | Muscle | family |
| c.47.1.5 | Muscle | family |

| | | |
|---|---|---|
| c.55.1.1 | Muscle | family |
| c.61.1.0 | Muscle | family |
| c.61.1.2 | Muscle | family |
| c.66.1.42 | Muscle | family |
| d.122.1.1 | Muscle | family |
| d.128.1.2 | Muscle | family |
| d.144.1.0 | Muscle | family |
| d.41.4.1 | Muscle | family |
| d.42.1.0 | Muscle | family |
| d.58.46.1 | Muscle | family |
| d.58.62.1 | Muscle | family |
| d.64.1.1 | Muscle | family |
| d.81.1.1 | Muscle | family |
| f.17.2.1 | Muscle | family |
| f.21.1.2 | Muscle | family |
| f.25.1.1 | Muscle | family |
| f.32.1.1 | Muscle | family |
| No-Fold | Muscle | family |
| a.16.1.2 | Ovary | family |
| a.25.1.0 | Ovary | family |
| a.4.5.28 | Ovary | family |
| a.4.7.1 | Ovary | family |
| a.45.1.1 | Ovary | family |
| b.1.1.0 | Ovary | family |
| b.1.1.2 | Ovary | family |
| b.29.1.0 | Ovary | family |
| b.34.5.7 | Ovary | family |
| b.39.1.1 | Ovary | family |
| b.40.4.5 | Ovary | family |
| b.43.3.1 | Ovary | family |

| | | |
|---|---|---|
| b.43.3.3 | Ovary | family |
| b.44.1.0 | Ovary | family |
| b.6.1.3 | Ovary | family |
| b.88.1.2 | Ovary | family |
| c.2.1.3 | Ovary | family |
| c.21.1.1 | Ovary | family |
| c.37.1.0 | Ovary | family |
| c.47.1.5 | Ovary | family |
| c.55.1.1 | Ovary | family |
| c.61.1.0 | Ovary | family |
| c.61.1.2 | Ovary | family |
| c.62.1.1 | Ovary | family |
| c.66.1.42 | Ovary | family |
| d.122.1.1 | Ovary | family |
| d.141.1.1 | Ovary | family |
| d.144.1.0 | Ovary | family |
| d.19.1.1 | Ovary | family |
| d.41.4.1 | Ovary | family |
| d.47.1.1 | Ovary | family |
| d.55.1.1 | Ovary | family |
| d.58.12.1 | Ovary | family |
| d.58.46.1 | Ovary | family |
| d.58.62.1 | Ovary | family |
| d.58.7.1 | Ovary | family |
| d.64.1.1 | Ovary | family |
| d.66.1.4 | Ovary | family |
| d.79.3.1 | Ovary | family |
| d.81.1.1 | Ovary | family |
| e.1.1.0 | Ovary | family |
| e.24.1.1 | Ovary | family |

| | | |
|---|---|---|
| f.17.2.1 | Ovary | family |
| f.18.1.1 | Ovary | family |
| f.21.1.2 | Ovary | family |
| f.25.1.1 | Ovary | family |
| f.32.1.1 | Ovary | family |
| g.28.1.0 | Ovary | family |
| g.41.8.4 | Ovary | family |
| No-Fold | Ovary | family |
| a.39.1.5 | Prostate | family |
| a.45.1.1 | Prostate | family |
| b.34.5.7 | Prostate | family |
| b.40.4.5 | Prostate | family |
| b.43.3.1 | Prostate | family |
| b.44.1.0 | Prostate | family |
| c.2.1.3 | Prostate | family |
| c.37.1.0 | Prostate | family |
| c.47.1.5 | Prostate | family |
| c.55.1.1 | Prostate | family |
| c.61.1.0 | Prostate | family |
| c.61.1.2 | Prostate | family |
| d.122.1.1 | Prostate | family |
| d.144.1.0 | Prostate | family |
| d.58.46.1 | Prostate | family |
| d.58.62.1 | Prostate | family |
| d.64.1.1 | Prostate | family |
| d.79.3.1 | Prostate | family |
| d.81.1.1 | Prostate | family |
| No-Fold | Prostate | family |
| a.25.1.0 | SmallIntestine | family |
| a.39.1.5 | SmallIntestine | family |

| | | |
|---|---|---|
| a.45.1.1 | SmallIntestine | family |
| b.29.1.0 | SmallIntestine | family |
| b.43.3.1 | SmallIntestine | family |
| b.44.1.0 | SmallIntestine | family |
| b.88.1.2 | SmallIntestine | family |
| c.37.1.0 | SmallIntestine | family |
| c.47.1.5 | SmallIntestine | family |
| c.55.1.1 | SmallIntestine | family |
| d.144.1.0 | SmallIntestine | family |
| f.25.1.1 | SmallIntestine | family |
| g.28.1.0 | SmallIntestine | family |
| No-Fold | SmallIntestine | family |
| a.45.1.1 | Spleen | family |
| b.1.1.2 | Spleen | family |
| b.43.3.1 | Spleen | family |
| b.44.1.0 | Spleen | family |
| b.88.1.2 | Spleen | family |
| c.21.1.1 | Spleen | family |
| c.37.1.0 | Spleen | family |
| c.55.1.1 | Spleen | family |
| d.144.1.0 | Spleen | family |
| d.19.1.1 | Spleen | family |
| d.58.7.1 | Spleen | family |
| d.64.1.1 | Spleen | family |
| f.25.1.1 | Spleen | family |
| No-Fold | Spleen | family |
| b.43.3.1 | Stomach | family |
| b.44.1.0 | Stomach | family |
| c.21.1.1 | Stomach | family |
| c.37.1.0 | Stomach | family |

| | | |
|---|---|---|
| c.55.1.1 | Stomach | family |
| d.144.1.0 | Stomach | family |
| d.41.4.1 | Stomach | family |
| d.79.3.1 | Stomach | family |
| f.18.1.1 | Stomach | family |
| f.21.1.2 | Stomach | family |
| f.25.1.1 | Stomach | family |
| f.32.1.1 | Stomach | family |
| g.41.8.4 | Stomach | family |
| No-Fold | Stomach | family |
| a.25.1.0 | Thyroid | family |
| a.45.1.1 | Thyroid | family |
| b.1.1.2 | Thyroid | family |
| b.34.5.7 | Thyroid | family |
| b.40.4.5 | Thyroid | family |
| b.43.3.1 | Thyroid | family |
| b.44.1.0 | Thyroid | family |
| b.6.1.3 | Thyroid | family |
| b.88.1.2 | Thyroid | family |
| c.2.1.3 | Thyroid | family |
| c.21.1.1 | Thyroid | family |
| c.37.1.0 | Thyroid | family |
| c.37.1.8 | Thyroid | family |
| c.47.1.10 | Thyroid | family |
| c.47.1.5 | Thyroid | family |
| c.55.1.1 | Thyroid | family |
| c.61.1.0 | Thyroid | family |
| c.61.1.2 | Thyroid | family |
| c.66.1.42 | Thyroid | family |
| d.122.1.1 | Thyroid | family |

| | | |
|---|---|---|
| d.144.1.0 | Thyroid | family |
| d.41.4.1 | Thyroid | family |
| d.55.1.1 | Thyroid | family |
| d.58.46.1 | Thyroid | family |
| d.58.62.1 | Thyroid | family |
| d.64.1.1 | Thyroid | family |
| d.66.1.4 | Thyroid | family |
| d.79.3.1 | Thyroid | family |
| d.81.1.1 | Thyroid | family |
| f.17.2.1 | Thyroid | family |
| f.18.1.1 | Thyroid | family |
| f.21.1.2 | Thyroid | family |
| f.25.1.1 | Thyroid | family |
| f.32.1.1 | Thyroid | family |
| No-Fold | Thyroid | family |
| a.39.1.5 | Vagina | family |
| a.45.1.1 | Vagina | family |
| a.75.1.1 | Vagina | family |
| b.15.1.1 | Vagina | family |
| b.29.1.0 | Vagina | family |
| b.34.5.7 | Vagina | family |
| b.40.4.5 | Vagina | family |
| b.43.3.1 | Vagina | family |
| b.43.3.3 | Vagina | family |
| b.44.1.0 | Vagina | family |
| b.6.1.3 | Vagina | family |
| b.88.1.2 | Vagina | family |
| c.2.1.3 | Vagina | family |
| c.21.1.1 | Vagina | family |
| c.37.1.0 | Vagina | family |

| | | |
|---|---|---|
| c.47.1.5 | Vagina | family |
| c.55.1.1 | Vagina | family |
| c.55.4.1 | Vagina | family |
| c.61.1.0 | Vagina | family |
| c.61.1.2 | Vagina | family |
| c.66.1.42 | Vagina | family |
| d.141.1.1 | Vagina | family |
| d.144.1.0 | Vagina | family |
| d.355.1.1 | Vagina | family |
| d.41.4.1 | Vagina | family |
| d.55.1.1 | Vagina | family |
| d.58.46.1 | Vagina | family |
| d.58.62.1 | Vagina | family |
| d.58.7.1 | Vagina | family |
| d.64.1.1 | Vagina | family |
| d.66.1.4 | Vagina | family |
| d.79.3.1 | Vagina | family |
| d.81.1.1 | Vagina | family |
| e.24.1.1 | Vagina | family |
| f.17.2.1 | Vagina | family |
| f.18.1.1 | Vagina | family |
| f.21.1.2 | Vagina | family |
| f.25.1.1 | Vagina | family |
| f.32.1.1 | Vagina | family |
| g.28.1.0 | Vagina | family |
| g.41.8.4 | Vagina | family |
| No-Fold | Vagina | family |
| a.25.1.0 | WholeBlood | family |
| b.1.1.0 | WholeBlood | family |
| b.1.1.2 | WholeBlood | family |

| | | |
|---|---|---|
| c.37.1.0 | WholeBlood | family |
| c.55.1.1 | WholeBlood | family |
| No-Fold | WholeBlood | family |
| a.135.1 | Adipose | superfamily |
| a.25.1 | Adipose | superfamily |
| a.39.1 | Adipose | superfamily |
| a.45.1 | Adipose | superfamily |
| b.1.1 | Adipose | superfamily |
| b.29.1 | Adipose | superfamily |
| b.34.5 | Adipose | superfamily |
| b.40.4 | Adipose | superfamily |
| b.43.3 | Adipose | superfamily |
| b.44.1 | Adipose | superfamily |
| b.6.1 | Adipose | superfamily |
| b.88.1 | Adipose | superfamily |
| c.2.1 | Adipose | superfamily |
| c.21.1 | Adipose | superfamily |
| c.37.1 | Adipose | superfamily |
| c.47.1 | Adipose | superfamily |
| c.55.1 | Adipose | superfamily |
| c.61.1 | Adipose | superfamily |
| c.66.1 | Adipose | superfamily |
| d.144.1 | Adipose | superfamily |
| d.41.4 | Adipose | superfamily |
| d.58.46 | Adipose | superfamily |
| d.58.62 | Adipose | superfamily |
| d.64.1 | Adipose | superfamily |
| d.79.3 | Adipose | superfamily |
| f.17.2 | Adipose | superfamily |
| f.18.1 | Adipose | superfamily |

| | | |
|---|---|---|
| f.21.1 | Adipose | superfamily |
| f.25.1 | Adipose | superfamily |
| f.32.1 | Adipose | superfamily |
| g.68.1 | Adipose | superfamily |
| No-Fold | Adipose | superfamily |
| a.118.7 | Brain | superfamily |
| a.25.1 | Brain | superfamily |
| a.39.1 | Brain | superfamily |
| a.77.1 | Brain | superfamily |
| b.17.1 | Brain | superfamily |
| b.43.3 | Brain | superfamily |
| b.44.1 | Brain | superfamily |
| c.2.1 | Brain | superfamily |
| c.37.1 | Brain | superfamily |
| c.55.1 | Brain | superfamily |
| c.61.1 | Brain | superfamily |
| d.122.1 | Brain | superfamily |
| d.144.1 | Brain | superfamily |
| d.81.1 | Brain | superfamily |
| g.37.1 | Brain | superfamily |
| No-Fold | Brain | superfamily |
| a.39.1 | Colon | superfamily |
| c.37.1 | Colon | superfamily |
| No-Fold | Colon | superfamily |
| a.25.1 | Esophagus | superfamily |
| a.39.1 | Esophagus | superfamily |
| a.45.1 | Esophagus | superfamily |
| b.1.1 | Esophagus | superfamily |
| b.15.1 | Esophagus | superfamily |
| b.29.1 | Esophagus | superfamily |

| | | |
|---|---|---|
| b.34.5 | Esophagus | superfamily |
| b.40.4 | Esophagus | superfamily |
| b.43.3 | Esophagus | superfamily |
| b.44.1 | Esophagus | superfamily |
| b.6.1 | Esophagus | superfamily |
| b.88.1 | Esophagus | superfamily |
| c.2.1 | Esophagus | superfamily |
| c.21.1 | Esophagus | superfamily |
| c.37.1 | Esophagus | superfamily |
| c.47.1 | Esophagus | superfamily |
| c.55.1 | Esophagus | superfamily |
| c.61.1 | Esophagus | superfamily |
| d.141.1 | Esophagus | superfamily |
| d.144.1 | Esophagus | superfamily |
| d.41.4 | Esophagus | superfamily |
| d.55.1 | Esophagus | superfamily |
| d.58.46 | Esophagus | superfamily |
| d.79.3 | Esophagus | superfamily |
| f.17.2 | Esophagus | superfamily |
| f.18.1 | Esophagus | superfamily |
| f.21.1 | Esophagus | superfamily |
| f.25.1 | Esophagus | superfamily |
| f.32.1 | Esophagus | superfamily |
| g.41.8 | Esophagus | superfamily |
| No-Fold | Esophagus | superfamily |
| a.39.1 | Heart | superfamily |
| b.1.1 | Heart | superfamily |
| b.43.3 | Heart | superfamily |
| b.44.1 | Heart | superfamily |
| c.37.1 | Heart | superfamily |

| | | |
|---|---|---|
| c.47.1 | Heart | superfamily |
| c.55.1 | Heart | superfamily |
| d.144.1 | Heart | superfamily |
| d.211.1 | Heart | superfamily |
| No-Fold | Heart | superfamily |
| b.43.3 | Kidney | superfamily |
| b.44.1 | Kidney | superfamily |
| c.37.1 | Kidney | superfamily |
| d.64.1 | Kidney | superfamily |
| No-Fold | Kidney | superfamily |
| a.126.1 | Liver | superfamily |
| a.24.1 | Liver | superfamily |
| a.25.1 | Liver | superfamily |
| b.1.1 | Liver | superfamily |
| b.29.1 | Liver | superfamily |
| b.43.3 | Liver | superfamily |
| b.47.1 | Liver | superfamily |
| b.6.1 | Liver | superfamily |
| b.60.1 | Liver | superfamily |
| c.2.1 | Liver | superfamily |
| c.37.1 | Liver | superfamily |
| c.47.1 | Liver | superfamily |
| c.55.1 | Liver | superfamily |
| c.62.1 | Liver | superfamily |
| c.66.1 | Liver | superfamily |
| c.69.1 | Liver | superfamily |
| d.171.1 | Liver | superfamily |
| d.92.1 | Liver | superfamily |
| e.1.1 | Liver | superfamily |
| f.18.1 | Liver | superfamily |

| | | |
|---|---|---|
| g.18.1 | Liver | superfamily |
| g.45.1 | Liver | superfamily |
| g.8.1 | Liver | superfamily |
| No-Fold | Liver | superfamily |
| a.39.1 | Lung | superfamily |
| c.37.1 | Lung | superfamily |
| d.64.1 | Lung | superfamily |
| No-Fold | Lung | superfamily |
| a.238.1 | Muscle | superfamily |
| a.26.1 | Muscle | superfamily |
| a.39.1 | Muscle | superfamily |
| a.40.1 | Muscle | superfamily |
| a.45.1 | Muscle | superfamily |
| a.83.1 | Muscle | superfamily |
| b.1.1 | Muscle | superfamily |
| b.1.18 | Muscle | superfamily |
| b.15.1 | Muscle | superfamily |
| b.29.1 | Muscle | superfamily |
| b.34.12 | Muscle | superfamily |
| b.40.4 | Muscle | superfamily |
| b.43.3 | Muscle | superfamily |
| b.44.1 | Muscle | superfamily |
| b.49.1 | Muscle | superfamily |
| b.6.1 | Muscle | superfamily |
| b.68.11 | Muscle | superfamily |
| b.88.1 | Muscle | superfamily |
| c.1.10 | Muscle | superfamily |
| c.2.1 | Muscle | superfamily |
| c.21.1 | Muscle | superfamily |
| c.37.1 | Muscle | superfamily |

| | | |
|---|---|---|
| c.47.1 | Muscle | superfamily |
| c.55.1 | Muscle | superfamily |
| c.61.1 | Muscle | superfamily |
| c.66.1 | Muscle | superfamily |
| d.122.1 | Muscle | superfamily |
| d.128.1 | Muscle | superfamily |
| d.144.1 | Muscle | superfamily |
| d.41.4 | Muscle | superfamily |
| d.42.1 | Muscle | superfamily |
| d.58.46 | Muscle | superfamily |
| d.58.62 | Muscle | superfamily |
| d.64.1 | Muscle | superfamily |
| d.81.1 | Muscle | superfamily |
| f.17.2 | Muscle | superfamily |
| f.21.1 | Muscle | superfamily |
| f.25.1 | Muscle | superfamily |
| f.32.1 | Muscle | superfamily |
| No-Fold | Muscle | superfamily |
| a.16.1 | Ovary | superfamily |
| a.25.1 | Ovary | superfamily |
| a.39.1 | Ovary | superfamily |
| a.4.5 | Ovary | superfamily |
| a.4.7 | Ovary | superfamily |
| a.45.1 | Ovary | superfamily |
| b.1.1 | Ovary | superfamily |
| b.29.1 | Ovary | superfamily |
| b.34.5 | Ovary | superfamily |
| b.39.1 | Ovary | superfamily |
| b.40.3 | Ovary | superfamily |
| b.40.4 | Ovary | superfamily |

| | | |
|---|---|---|
| b.43.3 | Ovary | superfamily |
| b.44.1 | Ovary | superfamily |
| b.47.1 | Ovary | superfamily |
| b.6.1 | Ovary | superfamily |
| b.88.1 | Ovary | superfamily |
| c.2.1 | Ovary | superfamily |
| c.21.1 | Ovary | superfamily |
| c.37.1 | Ovary | superfamily |
| c.47.1 | Ovary | superfamily |
| c.55.1 | Ovary | superfamily |
| c.61.1 | Ovary | superfamily |
| c.62.1 | Ovary | superfamily |
| c.66.1 | Ovary | superfamily |
| d.122.1 | Ovary | superfamily |
| d.141.1 | Ovary | superfamily |
| d.144.1 | Ovary | superfamily |
| d.19.1 | Ovary | superfamily |
| d.41.4 | Ovary | superfamily |
| d.47.1 | Ovary | superfamily |
| d.55.1 | Ovary | superfamily |
| d.58.12 | Ovary | superfamily |
| d.58.46 | Ovary | superfamily |
| d.58.62 | Ovary | superfamily |
| d.58.7 | Ovary | superfamily |
| d.64.1 | Ovary | superfamily |
| d.66.1 | Ovary | superfamily |
| d.79.3 | Ovary | superfamily |
| d.81.1 | Ovary | superfamily |
| e.1.1 | Ovary | superfamily |
| e.24.1 | Ovary | superfamily |

| | | |
|---|---|---|
| f.17.2 | Ovary | superfamily |
| f.18.1 | Ovary | superfamily |
| f.21.1 | Ovary | superfamily |
| f.25.1 | Ovary | superfamily |
| f.32.1 | Ovary | superfamily |
| g.28.1 | Ovary | superfamily |
| g.41.8 | Ovary | superfamily |
| No-Fold | Ovary | superfamily |
| a.39.1 | Prostate | superfamily |
| a.45.1 | Prostate | superfamily |
| b.29.1 | Prostate | superfamily |
| b.34.5 | Prostate | superfamily |
| b.40.4 | Prostate | superfamily |
| b.43.3 | Prostate | superfamily |
| b.44.1 | Prostate | superfamily |
| c.2.1 | Prostate | superfamily |
| c.37.1 | Prostate | superfamily |
| c.47.1 | Prostate | superfamily |
| c.55.1 | Prostate | superfamily |
| c.61.1 | Prostate | superfamily |
| d.122.1 | Prostate | superfamily |
| d.144.1 | Prostate | superfamily |
| d.58.46 | Prostate | superfamily |
| d.58.62 | Prostate | superfamily |
| d.64.1 | Prostate | superfamily |
| d.79.3 | Prostate | superfamily |
| d.81.1 | Prostate | superfamily |
| No-Fold | Prostate | superfamily |
| a.25.1 | SmallIntestine | superfamily |
| a.39.1 | SmallIntestine | superfamily |

| | | |
|---|---|---|
| a.45.1 | SmallIntestine | superfamily |
| b.1.1 | SmallIntestine | superfamily |
| b.29.1 | SmallIntestine | superfamily |
| b.43.3 | SmallIntestine | superfamily |
| b.44.1 | SmallIntestine | superfamily |
| b.88.1 | SmallIntestine | superfamily |
| c.2.1 | SmallIntestine | superfamily |
| c.37.1 | SmallIntestine | superfamily |
| c.47.1 | SmallIntestine | superfamily |
| c.55.1 | SmallIntestine | superfamily |
| c.66.1 | SmallIntestine | superfamily |
| d.144.1 | SmallIntestine | superfamily |
| f.25.1 | SmallIntestine | superfamily |
| g.28.1 | SmallIntestine | superfamily |
| No-Fold | SmallIntestine | superfamily |
| a.39.1 | Spleen | superfamily |
| a.45.1 | Spleen | superfamily |
| b.1.1 | Spleen | superfamily |
| b.1.18 | Spleen | superfamily |
| b.43.3 | Spleen | superfamily |
| b.44.1 | Spleen | superfamily |
| b.88.1 | Spleen | superfamily |
| c.21.1 | Spleen | superfamily |
| c.37.1 | Spleen | superfamily |
| c.55.1 | Spleen | superfamily |
| d.144.1 | Spleen | superfamily |
| d.19.1 | Spleen | superfamily |
| d.58.7 | Spleen | superfamily |
| d.64.1 | Spleen | superfamily |
| f.25.1 | Spleen | superfamily |

| | | |
|---|---|---|
| No-Fold | Spleen | superfamily |
| b.1.1 | Stomach | superfamily |
| b.43.3 | Stomach | superfamily |
| b.44.1 | Stomach | superfamily |
| c.21.1 | Stomach | superfamily |
| c.37.1 | Stomach | superfamily |
| c.55.1 | Stomach | superfamily |
| d.144.1 | Stomach | superfamily |
| d.41.4 | Stomach | superfamily |
| d.79.3 | Stomach | superfamily |
| f.18.1 | Stomach | superfamily |
| f.21.1 | Stomach | superfamily |
| f.25.1 | Stomach | superfamily |
| f.32.1 | Stomach | superfamily |
| g.41.8 | Stomach | superfamily |
| No-Fold | Stomach | superfamily |
| a.25.1 | Thyroid | superfamily |
| a.39.1 | Thyroid | superfamily |
| a.45.1 | Thyroid | superfamily |
| b.1.1 | Thyroid | superfamily |
| b.29.1 | Thyroid | superfamily |
| b.34.5 | Thyroid | superfamily |
| b.40.4 | Thyroid | superfamily |
| b.43.3 | Thyroid | superfamily |
| b.44.1 | Thyroid | superfamily |
| b.6.1 | Thyroid | superfamily |
| b.88.1 | Thyroid | superfamily |
| c.2.1 | Thyroid | superfamily |
| c.21.1 | Thyroid | superfamily |
| c.37.1 | Thyroid | superfamily |

| | | |
|---|---|---|
| c.47.1 | Thyroid | superfamily |
| c.55.1 | Thyroid | superfamily |
| c.61.1 | Thyroid | superfamily |
| c.66.1 | Thyroid | superfamily |
| d.122.1 | Thyroid | superfamily |
| d.144.1 | Thyroid | superfamily |
| d.211.1 | Thyroid | superfamily |
| d.41.4 | Thyroid | superfamily |
| d.55.1 | Thyroid | superfamily |
| d.58.46 | Thyroid | superfamily |
| d.58.62 | Thyroid | superfamily |
| d.64.1 | Thyroid | superfamily |
| d.66.1 | Thyroid | superfamily |
| d.79.3 | Thyroid | superfamily |
| d.81.1 | Thyroid | superfamily |
| f.17.2 | Thyroid | superfamily |
| f.18.1 | Thyroid | superfamily |
| f.21.1 | Thyroid | superfamily |
| f.25.1 | Thyroid | superfamily |
| f.32.1 | Thyroid | superfamily |
| No-Fold | Thyroid | superfamily |
| a.39.1 | Vagina | superfamily |
| a.45.1 | Vagina | superfamily |
| a.75.1 | Vagina | superfamily |
| b.1.1 | Vagina | superfamily |
| b.1.18 | Vagina | superfamily |
| b.15.1 | Vagina | superfamily |
| b.29.1 | Vagina | superfamily |
| b.34.5 | Vagina | superfamily |
| b.40.4 | Vagina | superfamily |

| | | |
|---|---|---|
| b.43.3 | Vagina | superfamily |
| b.44.1 | Vagina | superfamily |
| b.47.1 | Vagina | superfamily |
| b.6.1 | Vagina | superfamily |
| b.88.1 | Vagina | superfamily |
| c.2.1 | Vagina | superfamily |
| c.21.1 | Vagina | superfamily |
| c.37.1 | Vagina | superfamily |
| c.47.1 | Vagina | superfamily |
| c.55.1 | Vagina | superfamily |
| c.55.4 | Vagina | superfamily |
| c.61.1 | Vagina | superfamily |
| c.66.1 | Vagina | superfamily |
| d.141.1 | Vagina | superfamily |
| d.144.1 | Vagina | superfamily |
| d.355.1 | Vagina | superfamily |
| d.41.4 | Vagina | superfamily |
| d.55.1 | Vagina | superfamily |
| d.58.46 | Vagina | superfamily |
| d.58.62 | Vagina | superfamily |
| d.58.7 | Vagina | superfamily |
| d.64.1 | Vagina | superfamily |
| d.66.1 | Vagina | superfamily |
| d.79.3 | Vagina | superfamily |
| d.81.1 | Vagina | superfamily |
| e.24.1 | Vagina | superfamily |
| f.17.2 | Vagina | superfamily |
| f.18.1 | Vagina | superfamily |
| f.21.1 | Vagina | superfamily |
| f.25.1 | Vagina | superfamily |

| | | |
|---|---|---|
| f.32.1 | Vagina | superfamily |
| g.28.1 | Vagina | superfamily |
| g.41.8 | Vagina | superfamily |
| No-Fold | Vagina | superfamily |
| a.25.1 | WholeBlood | superfamily |
| a.39.1 | WholeBlood | superfamily |
| b.1.1 | WholeBlood | superfamily |
| b.1.18 | WholeBlood | superfamily |
| c.37.1 | WholeBlood | superfamily |
| c.55.1 | WholeBlood | superfamily |
| d.108.1 | WholeBlood | superfamily |
| d.144.1 | WholeBlood | superfamily |
| No-Fold | WholeBlood | superfamily |
| a.135 | Adipose | fold |
| a.25 | Adipose | fold |
| a.39 | Adipose | fold |
| a.45 | Adipose | fold |
| b.1 | Adipose | fold |
| b.29 | Adipose | fold |
| b.34 | Adipose | fold |
| b.40 | Adipose | fold |
| b.43 | Adipose | fold |
| b.44 | Adipose | fold |
| b.6 | Adipose | fold |
| b.88 | Adipose | fold |
| c.2 | Adipose | fold |
| c.21 | Adipose | fold |
| c.37 | Adipose | fold |
| c.47 | Adipose | fold |
| c.55 | Adipose | fold |

| | | |
|---|---|---|
| c.61 | Adipose | fold |
| c.66 | Adipose | fold |
| d.144 | Adipose | fold |
| d.41 | Adipose | fold |
| d.58 | Adipose | fold |
| d.64 | Adipose | fold |
| d.79 | Adipose | fold |
| f.17 | Adipose | fold |
| f.18 | Adipose | fold |
| f.21 | Adipose | fold |
| f.25 | Adipose | fold |
| f.32 | Adipose | fold |
| g.68 | Adipose | fold |
| No-Fold | Adipose | fold |
| a.118 | Brain | fold |
| a.25 | Brain | fold |
| a.39 | Brain | fold |
| a.77 | Brain | fold |
| b.1 | Brain | fold |
| b.17 | Brain | fold |
| b.34 | Brain | fold |
| b.43 | Brain | fold |
| b.44 | Brain | fold |
| c.2 | Brain | fold |
| c.37 | Brain | fold |
| c.55 | Brain | fold |
| c.61 | Brain | fold |
| d.122 | Brain | fold |
| d.144 | Brain | fold |
| d.81 | Brain | fold |

| | | |
|---|---|---|
| g.37 | Brain | fold |
| No-Fold | Brain | fold |
| a.39 | Colon | fold |
| b.1 | Colon | fold |
| c.37 | Colon | fold |
| c.55 | Colon | fold |
| d.58 | Colon | fold |
| No-Fold | Colon | fold |
| a.25 | Esophagus | fold |
| a.39 | Esophagus | fold |
| a.4 | Esophagus | fold |
| a.45 | Esophagus | fold |
| b.1 | Esophagus | fold |
| b.15 | Esophagus | fold |
| b.29 | Esophagus | fold |
| b.34 | Esophagus | fold |
| b.40 | Esophagus | fold |
| b.43 | Esophagus | fold |
| b.44 | Esophagus | fold |
| b.6 | Esophagus | fold |
| b.88 | Esophagus | fold |
| c.2 | Esophagus | fold |
| c.21 | Esophagus | fold |
| c.37 | Esophagus | fold |
| c.47 | Esophagus | fold |
| c.55 | Esophagus | fold |
| c.61 | Esophagus | fold |
| d.141 | Esophagus | fold |
| d.144 | Esophagus | fold |
| d.41 | Esophagus | fold |

| | | |
|---|---|---|
| d.55 | Esophagus | fold |
| d.58 | Esophagus | fold |
| d.79 | Esophagus | fold |
| f.17 | Esophagus | fold |
| f.18 | Esophagus | fold |
| f.21 | Esophagus | fold |
| f.25 | Esophagus | fold |
| f.32 | Esophagus | fold |
| g.41 | Esophagus | fold |
| No-Fold | Esophagus | fold |
| a.39 | Heart | fold |
| b.1 | Heart | fold |
| b.34 | Heart | fold |
| b.43 | Heart | fold |
| b.44 | Heart | fold |
| c.37 | Heart | fold |
| c.47 | Heart | fold |
| c.55 | Heart | fold |
| d.144 | Heart | fold |
| d.211 | Heart | fold |
| d.58 | Heart | fold |
| No-Fold | Heart | fold |
| b.40 | Kidney | fold |
| b.43 | Kidney | fold |
| b.44 | Kidney | fold |
| c.37 | Kidney | fold |
| c.55 | Kidney | fold |
| d.58 | Kidney | fold |
| d.64 | Kidney | fold |
| No-Fold | Kidney | fold |

| | | |
|---|---|---|
| a.126 | Liver | fold |
| a.24 | Liver | fold |
| a.25 | Liver | fold |
| b.1 | Liver | fold |
| b.29 | Liver | fold |
| b.43 | Liver | fold |
| b.47 | Liver | fold |
| b.6 | Liver | fold |
| b.60 | Liver | fold |
| c.1 | Liver | fold |
| c.2 | Liver | fold |
| c.37 | Liver | fold |
| c.47 | Liver | fold |
| c.55 | Liver | fold |
| c.62 | Liver | fold |
| c.66 | Liver | fold |
| c.69 | Liver | fold |
| d.171 | Liver | fold |
| d.58 | Liver | fold |
| d.92 | Liver | fold |
| e.1 | Liver | fold |
| f.18 | Liver | fold |
| g.18 | Liver | fold |
| g.45 | Liver | fold |
| g.8 | Liver | fold |
| No-Fold | Liver | fold |
| a.39 | Lung | fold |
| b.1 | Lung | fold |
| b.43 | Lung | fold |
| c.37 | Lung | fold |

| | | |
|---|---|---|
| c.55 | Lung | fold |
| d.64 | Lung | fold |
| No-Fold | Lung | fold |
| a.238 | Muscle | fold |
| a.26 | Muscle | fold |
| a.39 | Muscle | fold |
| a.40 | Muscle | fold |
| a.45 | Muscle | fold |
| a.83 | Muscle | fold |
| b.1 | Muscle | fold |
| b.15 | Muscle | fold |
| b.29 | Muscle | fold |
| b.34 | Muscle | fold |
| b.40 | Muscle | fold |
| b.43 | Muscle | fold |
| b.44 | Muscle | fold |
| b.49 | Muscle | fold |
| b.6 | Muscle | fold |
| b.68 | Muscle | fold |
| b.88 | Muscle | fold |
| c.1 | Muscle | fold |
| c.2 | Muscle | fold |
| c.21 | Muscle | fold |
| c.37 | Muscle | fold |
| c.47 | Muscle | fold |
| c.55 | Muscle | fold |
| c.61 | Muscle | fold |
| c.66 | Muscle | fold |
| d.122 | Muscle | fold |
| d.128 | Muscle | fold |

| | | |
|---|---|---|
| d.144 | Muscle | fold |
| d.41 | Muscle | fold |
| d.42 | Muscle | fold |
| d.58 | Muscle | fold |
| d.64 | Muscle | fold |
| d.81 | Muscle | fold |
| f.17 | Muscle | fold |
| f.21 | Muscle | fold |
| f.25 | Muscle | fold |
| f.32 | Muscle | fold |
| No-Fold | Muscle | fold |
| a.16 | Ovary | fold |
| a.25 | Ovary | fold |
| a.39 | Ovary | fold |
| a.4 | Ovary | fold |
| a.45 | Ovary | fold |
| b.1 | Ovary | fold |
| b.29 | Ovary | fold |
| b.34 | Ovary | fold |
| b.39 | Ovary | fold |
| b.40 | Ovary | fold |
| b.43 | Ovary | fold |
| b.44 | Ovary | fold |
| b.47 | Ovary | fold |
| b.6 | Ovary | fold |
| b.88 | Ovary | fold |
| c.2 | Ovary | fold |
| c.21 | Ovary | fold |
| c.37 | Ovary | fold |
| c.47 | Ovary | fold |

| | | |
|---|---|---|
| c.55 | Ovary | fold |
| c.61 | Ovary | fold |
| c.62 | Ovary | fold |
| c.66 | Ovary | fold |
| d.122 | Ovary | fold |
| d.141 | Ovary | fold |
| d.144 | Ovary | fold |
| d.19 | Ovary | fold |
| d.41 | Ovary | fold |
| d.47 | Ovary | fold |
| d.55 | Ovary | fold |
| d.58 | Ovary | fold |
| d.64 | Ovary | fold |
| d.66 | Ovary | fold |
| d.79 | Ovary | fold |
| d.81 | Ovary | fold |
| e.1 | Ovary | fold |
| e.24 | Ovary | fold |
| f.17 | Ovary | fold |
| f.18 | Ovary | fold |
| f.21 | Ovary | fold |
| f.25 | Ovary | fold |
| f.32 | Ovary | fold |
| g.28 | Ovary | fold |
| g.41 | Ovary | fold |
| No-Fold | Ovary | fold |
| a.39 | Prostate | fold |
| a.4 | Prostate | fold |
| a.45 | Prostate | fold |
| b.1 | Prostate | fold |

| | | |
|---|---|---|
| b.29 | Prostate | fold |
| b.34 | Prostate | fold |
| b.40 | Prostate | fold |
| b.43 | Prostate | fold |
| b.44 | Prostate | fold |
| c.2 | Prostate | fold |
| c.37 | Prostate | fold |
| c.47 | Prostate | fold |
| c.55 | Prostate | fold |
| c.61 | Prostate | fold |
| d.122 | Prostate | fold |
| d.144 | Prostate | fold |
| d.58 | Prostate | fold |
| d.64 | Prostate | fold |
| d.79 | Prostate | fold |
| d.81 | Prostate | fold |
| No-Fold | Prostate | fold |
| a.25 | SmallIntestine | fold |
| a.39 | SmallIntestine | fold |
| a.45 | SmallIntestine | fold |
| b.1 | SmallIntestine | fold |
| b.29 | SmallIntestine | fold |
| b.43 | SmallIntestine | fold |
| b.44 | SmallIntestine | fold |
| b.88 | SmallIntestine | fold |
| c.2 | SmallIntestine | fold |
| c.37 | SmallIntestine | fold |
| c.47 | SmallIntestine | fold |
| c.55 | SmallIntestine | fold |
| c.66 | SmallIntestine | fold |

| | | |
|---|---|---|
| d.144 | SmallIntestine | fold |
| d.58 | SmallIntestine | fold |
| f.25 | SmallIntestine | fold |
| g.28 | SmallIntestine | fold |
| No-Fold | SmallIntestine | fold |
| a.39 | Spleen | fold |
| a.4 | Spleen | fold |
| a.45 | Spleen | fold |
| b.1 | Spleen | fold |
| b.43 | Spleen | fold |
| b.44 | Spleen | fold |
| b.88 | Spleen | fold |
| c.21 | Spleen | fold |
| c.37 | Spleen | fold |
| c.55 | Spleen | fold |
| d.144 | Spleen | fold |
| d.19 | Spleen | fold |
| d.58 | Spleen | fold |
| d.64 | Spleen | fold |
| f.25 | Spleen | fold |
| No-Fold | Spleen | fold |
| a.4 | Stomach | fold |
| b.1 | Stomach | fold |
| b.34 | Stomach | fold |
| b.43 | Stomach | fold |
| b.44 | Stomach | fold |
| c.21 | Stomach | fold |
| c.37 | Stomach | fold |
| c.55 | Stomach | fold |
| d.144 | Stomach | fold |

| | | |
|---|---|---|
| d.41 | Stomach | fold |
| d.58 | Stomach | fold |
| d.79 | Stomach | fold |
| f.18 | Stomach | fold |
| f.21 | Stomach | fold |
| f.25 | Stomach | fold |
| f.32 | Stomach | fold |
| g.41 | Stomach | fold |
| No-Fold | Stomach | fold |
| a.25 | Thyroid | fold |
| a.39 | Thyroid | fold |
| a.4 | Thyroid | fold |
| a.45 | Thyroid | fold |
| b.1 | Thyroid | fold |
| b.29 | Thyroid | fold |
| b.34 | Thyroid | fold |
| b.40 | Thyroid | fold |
| b.43 | Thyroid | fold |
| b.44 | Thyroid | fold |
| b.6 | Thyroid | fold |
| b.88 | Thyroid | fold |
| c.2 | Thyroid | fold |
| c.21 | Thyroid | fold |
| c.37 | Thyroid | fold |
| c.47 | Thyroid | fold |
| c.55 | Thyroid | fold |
| c.61 | Thyroid | fold |
| c.66 | Thyroid | fold |
| d.122 | Thyroid | fold |
| d.144 | Thyroid | fold |

| | | |
|---|---|---|
| d.211 | Thyroid | fold |
| d.41 | Thyroid | fold |
| d.55 | Thyroid | fold |
| d.58 | Thyroid | fold |
| d.64 | Thyroid | fold |
| d.66 | Thyroid | fold |
| d.79 | Thyroid | fold |
| d.81 | Thyroid | fold |
| f.17 | Thyroid | fold |
| f.18 | Thyroid | fold |
| f.21 | Thyroid | fold |
| f.25 | Thyroid | fold |
| f.32 | Thyroid | fold |
| No-Fold | Thyroid | fold |
| a.39 | Vagina | fold |
| a.4 | Vagina | fold |
| a.45 | Vagina | fold |
| a.75 | Vagina | fold |
| b.1 | Vagina | fold |
| b.15 | Vagina | fold |
| b.29 | Vagina | fold |
| b.34 | Vagina | fold |
| b.40 | Vagina | fold |
| b.43 | Vagina | fold |
| b.44 | Vagina | fold |
| b.47 | Vagina | fold |
| b.6 | Vagina | fold |
| b.88 | Vagina | fold |
| c.2 | Vagina | fold |
| c.21 | Vagina | fold |

| | | |
|---|---|---|
| c.37 | Vagina | fold |
| c.47 | Vagina | fold |
| c.55 | Vagina | fold |
| c.61 | Vagina | fold |
| c.66 | Vagina | fold |
| d.141 | Vagina | fold |
| d.144 | Vagina | fold |
| d.355 | Vagina | fold |
| d.41 | Vagina | fold |
| d.55 | Vagina | fold |
| d.58 | Vagina | fold |
| d.64 | Vagina | fold |
| d.66 | Vagina | fold |
| d.79 | Vagina | fold |
| d.81 | Vagina | fold |
| e.24 | Vagina | fold |
| f.17 | Vagina | fold |
| f.18 | Vagina | fold |
| f.21 | Vagina | fold |
| f.25 | Vagina | fold |
| f.32 | Vagina | fold |
| g.28 | Vagina | fold |
| g.41 | Vagina | fold |
| No-Fold | Vagina | fold |
| a.25 | WholeBlood | fold |
| a.39 | WholeBlood | fold |
| b.1 | WholeBlood | fold |
| b.34 | WholeBlood | fold |
| c.1 | WholeBlood | fold |
| c.37 | WholeBlood | fold |

| | | |
|---|---|---|
| c.55 | WholeBlood | fold |
| d.108 | WholeBlood | fold |
| d.144 | WholeBlood | fold |
| d.15 | WholeBlood | fold |
| No-Fold | WholeBlood | fold |

**Table S8: GES and sGES of the whole blood tissue type across GTEx and ARCHS4, after outlier removal.** Each signature type is present in 100% of all ARCHS4 and GTEx whole blood samples, after outlier removal from ARCHS4. ‘ID’ refers to either the gene name, interproscan identifier or SCOPe identifier.

| ID | Name | Tissue | Signature Type |
|---|---|---|---|
| ACTB | Actin Beta | WholeBlood | gene |
| FTL | Ferritin Light Chain | WholeBlood | gene |
| IPR000719 | Protein kinase domain | WholeBlood | domain |
| IPR002048 | EF-hand domain | WholeBlood | domain |
| IPR004000 | Actin family | WholeBlood | domain |
| IPR005225 | Small GTP-binding protein domain | WholeBlood | domain |
| IPR007110 | Immunoglobulin-like domain | WholeBlood | domain |
| IPR008331 | Ferritin/DPS protein domain | WholeBlood | domain |
| IPR009040 | Ferritin-like diiron domain | WholeBlood | domain |
| b.1.1.2 | C1 set domains | WholeBlood | family |
| c.37.1.0 | Pyruvate oxidase and decarboxylase PP module | WholeBlood | family |
| c.55.1.1 | ROK | WholeBlood | family |
| No-Fold | N/A | WholeBlood | family |
| a.25.1 | Ferritin-like | WholeBlood | superfamily |
| a.39.1 | EF-hand | WholeBlood | superfamily |
| b.1.1 | Clathrin adaptor appendage domain | WholeBlood | superfamily |
| b.1.18 | Immunoglobulin | WholeBlood | superfamily |
| c.37.1 | P-loop containing nucleoside triphosphate hydrolases | WholeBlood | superfamily |
| c.55.1 | Actin-like ATPase domain | WholeBlood | superfamily |
| d.108.1 | Acyl-CoA N-acyltransferases (Nat) | WholeBlood | superfamily |
| d.144.1 | Protein kinase-like (PK-like) | WholeBlood | superfamily |
| No-Fold | N/A | WholeBlood | superfamily |

| | | | |
|---|---|---|---|
| a.25 | Ferritin-like | WholeBlood | fold |
| a.39 | EF Hand-like | WholeBlood | fold |
| b.1 | Immunoglobulin-like beta-sandwich | WholeBlood | fold |
| b.34 | SH3-like barrel | WholeBlood | fold |
| c.1 | TIM beta/alpha-barrel | WholeBlood | fold |
| c.37 | P-loop containing nucleoside triphosphate hydrolases | WholeBlood | fold |
| c.55 | Ribonuclease H-like motif | WholeBlood | fold |
| d.108 | Acyl-CoA N-acyltransferases (Nat) | WholeBlood | fold |
| d.144 | Protein kinase-like (PK-like) | WholeBlood | fold |
| d.15 | beta-Grasp (ubiquitin-like) | WholeBlood | fold |
| No-Fold | N/A | WholeBlood | fold |

**Table S9: Over-expressed folds in selected gene expression profiles from figure 6.** Each group is derived from Fig. 6. Each fold shown is overrepresented in each group of drugs.

| Group | SCOPe ID | Fold Name | Drug Category | Drug |
|---|---|---|---|---|
| 1 | a.13 | RAP domain-like | Anthracyclines | DOX;DAU;IDA |
| 1 | b.61 | Streptavidin-like | Anthracyclines | DOX;DAU;IDA |
| 2 | d.145 | FAD-binding/transporter-associated domain-like | Anthracyclines | IDA |
| 2 | b.64 | Mannose 6-phosphate receptor domain | Anthracyclines | IDA |
| 2 | c.123 | CoA-transferase family III (CaiB/BaiF) | Anthracyclines | IDA |
| 2 | a.56 | CO dehydrogenase ISP C-domain like | Anthracyclines | IDA |
| 2 | d.133 | Molybdenum cofactor-binding domain | Anthracyclines | IDA |
| 2 | a.42 | SWIB/MDM2 domain | Anthracyclines | IDA |
| 2 | d.41 | alpha/beta-Hammerhead | Anthracyclines | IDA |
| 2 | c.4 | Nucleotide-binding domain | Anthracyclines | IDA |
| 2 | c.31 | DHS-like NAD/FAD-binding domain | Anthracyclines | IDA |
| 2 | c.36 | Thiamin diphosphate-binding fold (THDP-binding) | Anthracyclines | IDA |
| 2 | a.24 | Four-helical up-and-down bundle | Anthracyclines | IDA |
| 2 | d.47 | Ribosomal L11/L12e N-terminal domain | Anthracyclines | IDA |
| 2 | b.35 | GroES-like | Anthracyclines | IDA |
| 2 | g.8 | BPTI-like | Anthracyclines | IDA |
| 3 | g.46 | Metallothionein | Protein Kinase Inhibitor | NIL;VEM;SOR;REG;PAZ |
| 3 | d.230 | Dodecin subunit-like | Protein Kinase Inhibitor | NIL;VEM;SOR;REG;PAZ |
| 4 | c.107 | DHH phosphoesterases | Protein Kinase Inhibitor | RUX;TRA |
| 4 | d.111 | PR-1-like | Protein Kinase Inhibitor | RUX;TRA |
| 4 | b.15 | HSP20-like chaperones | Protein Kinase Inhibitor | RUX;TRA |
| 4 | b.60 | Lipocalins | Protein Kinase Inhibitor | RUX;TRA |
| 4 | a.65 | Annexin | Protein Kinase Inhibitor | RUX;TRA |
| 4 | c.32 | Tubulin nucleotide-binding domain-like | Protein Kinase Inhibitor | RUX;TRA |
| 4 | a.25 | Ferritin-like | Protein Kinase Inhibitor | RUX;TRA |
| 5 | b.118 | FAS1 domain | Anthracyclines | EPI |
| 5 | f.56 | MAPEG domain-like | Anthracyclines | EPI |
| 5 | b.104 | P-domain of calnexin/calreticulin | Anthracyclines | EPI |
| 5 | d.389 | Menin N-terminal domain-like | Anthracyclines | EPI |
| 5 | b.17 | PEBP-like | Anthracyclines | EPI |
| 5 | d.49 | Signal recognition particle alu RNA binding heterodimer | Anthracyclines | EPI |
| 5 | a.144 | PABP domain-like | Anthracyclines | EPI |
| 5 | d.47 | Ribosomal L11/L12e N-terminal domain | Anthracyclines | EPI |

**Table S10: Under-expressed folds in selected gene expression profiles from fig. 6.** Each group is derived from Fig. 6. Each fold shown is under-represented in each group of drugs.

| Group | SCOPe ID | Fold Name | Drug Category | Drug |
|---|---|---|---|---|
| 1 | a.16 | S15/NS1 RNA-binding domain | Anthracyclines | DOX;DAU |
| 1 | a.27 | Anticodon-binding domain of a subclass of class I aminoacyl-tRNA synthetases | Anthracyclines | DOX;DAU |
| 1 | c.26 | Adenine nucleotide alpha hydrolase-like | Anthracyclines | DOX;DAU |
| 1 | b.53 | Ribosomal protein L25-like | Anthracyclines | DOX;DAU |
| 1 | c.51 | Anticodon-binding domain-like | Anthracyclines | DOX;DAU |
| 1 | d.104 | Class II aaRS and biotin synthetases | Anthracyclines | DOX;DAU |
| 2 | a.165 | Myosin phosphatase inhibitor 17kDa protein | Anthracyclines | IDA |
| 2 | d.126 | Pentein | Anthracyclines | IDA |
| 2 | d.389 | Menin N-terminal domain-like | Anthracyclines | IDA |
| 2 | g.75 | TSP type-3 repeat | Anthracyclines | IDA |
| 2 | g.1 | insulin like | Anthracyclines | IDA |
| 2 | g.28 | Thyroglobulin type-1 domain | Anthracyclines | IDA |
| 2 | g.27 | FnI-like domain | Anthracyclines | IDA |
| 3 | f.21 | Heme-binding four-helical bundle | Protein Kinase Inhibitor | SOR;REG;PAZ |
| 3 | f.32 | a domain/subunit of cytochrome bc1 complex (Ubiquinol-cytochrome c reductase) | Protein Kinase Inhibitor | SOR;REG;PAZ |
| 3 | c.49 | Pyruvate kinase C-terminal domain-like | Protein Kinase Inhibitor | SOR;REG;PAZ |
| 3 | a.29 | Bromodomain-like | Protein Kinase Inhibitor | SOR;REG;PAZ |
| 3 | e.6 | Acyl-CoA dehydrogenase NM domain-like | Protein Kinase Inhibitor | SOR;REG;PAZ |
| 3 | c.86 | Phosphoglycerate kinase | Protein Kinase Inhibitor | SOR;REG;PAZ |
| 3 | d.54 | Enolase N-terminal domain-like | Protein Kinase Inhibitor | SOR;REG;PAZ |
| 4 | a.22 | Histone-fold | Anthracyclines | EPI |
| 4 | d.153 | Ntn hydrolase-like | Anthracyclines | EPI |
| 4 | b.39 | Ribosomal protein L14 | Anthracyclines | EPI |
